# Super-Resolution Visual ProteomEx for Hard Tissues and Clinical Samples

**DOI:** 10.64898/2026.07.01.735813

**Authors:** Shuchang Zhao, Wei Tan, Angela Yiu, Tianning Liu, Xun Guo, Guilherme A. Câmara, Haodong Tian, Yuhao Wang, Guanqiao Dong, Cuiji Sun, Jiaxiang Ding, Pratiksha Patra, Andrew D. White, Samuel G. Rodriques, Ludovico Mitchener, Huan Zhou, Tiannan Guo, Yeqiang Liu, Kiryl D. Piatkevich

## Abstract

Correlating nanoscale tissue architecture with unbiased molecular composition within the same specimen remains a fundamental challenge in spatial biology. Current spatial proteomics methods either lack the imaging resolution required to resolve nanoscale tissue architecture or achieve only limited proteome coverage, and untargeted approaches capable of deep protein identification have yet to be integrated with super-resolution imaging within a single workflow. Here, we present microProteomEx, an integrated platform that combines hydrogel-assisted tissue expansion with mass spectrometry-compatible fluorescence imaging, laser capture microdissection, and bottom-up proteomics to achieve simultaneous three-dimensional super-resolution imaging (effective lateral resolution down to ∼47 nm on conventional diffraction-limited microscopes) and spatially resolved proteomics at a scalable lateral resolution of 29–100 µm (0.02–0.28 nL volumetric resolution). By optimizing fixation, protein anchoring, and a secondary re-embedding strategy, we extend the method to mechanically resilient tissues, including mouse kidney and heart, as well as to formalin-fixed, paraffin-embedded clinical specimens. We demonstrate single-glomerulus and single-plaque proteomics (450-1,000 proteins per structure), resolve proteomic differences between malignant melanoma and giant congenital melanocytic nevus in a rare pediatric case, and characterize the morphology-resolved molecular heterogeneity of cored versus diffuse amyloid-beta plaques in a mouse model of Alzheimer’s disease across two disease stages. microProteomEx establishes a broadly accessible framework for correlating tissue ultrastructure with deep spatial proteomics, with direct implications for disease biology, biomarker discovery, and precision medicine.

## Introduction

Deciphering tissue heterogeneity is fundamental to understanding normal physiology and pathological processes^1^. Despite remarkable advances in single-cell and spatially resolved -omics technologies, integrating microscale proteomics with super-resolution cytostructural imaging has remained challenging^2^, particularly for resilient tissues and archived clinical samples.

Hydrogel-assisted tissue magnification became a popular approach for volumetric multicolor super-resolution imaging under conventional diffraction-limited microscopes^3^, providing increasingly detailed images of subcellular ultrastructures across large tissue samples^4^. Despite a large diversity of developed protocols and corresponding chemical strategies, all sample magnification methods share a common framework^5–8^. First, a preserved biological sample is treated with chemical anchors to selectively modify biomolecules of interest, including proteins, nucleic acids, lipids, and carbohydrates/glycans, with functional groups that can be covalently integrated into acrylate-based polymer networks^9^. Then, the functionalized sample is infused with hydrogel monomers, followed by an *in situ* free-radical polymerization reaction that incorporates biomolecules into polymer chains. The formed tissue-hydrogel composite is mechanically homogenized by either enzymatic or physical-chemical treatment to facilitate isotropic magnification in aqueous buffers or pure water. Since its invention, hydrogel-assisted sample magnification was mainly used to achieve 3D super-resolution imaging with diffraction-limited microscopes, establishing Expansion Microscopy (ExM) as a new method for nanoscale imaging^7^. The retention of functional nucleic acids and proteins within expanded hydrogels enabled highly multiplexed imaging of DNA, RNA, and proteins using targeted approaches such as FISH and antibody labeling^10–12^. Later, the hydrogel milieu was found to be compatible with enzymatic reactions, enabling untargeted imaging-based *in situ* sequencing of the genome and transcriptome while mapping sequenced DNA and RNA molecules within their cellular context with nanoscale precision^13,14^. More recently, the ExM framework was employed to increase the spatial resolution of other imaging modalities, such as mass spectrometry imaging, enabling sub-micrometer spatial lipidomics^15,16^, histopathology^17^, and proteomics^18^ of preserved tissues. These applications demonstrate the versatility of the ExM workflow to increase the spatial resolution of various molecular techniques beyond fluorescence microscopy.

Recently, we extended the applicability of hydrogel-based tissue magnification to spatial proteomics by integrating it with bottom-up mass spectrometry (MS) proteomics^19^. According to the developed workflow, named ProteomEx, the preserved biological tissue sample is expanded up to ∼6-fold in linear dimensions and stained with a colorimetric dye, enabling precise manual microdissection of regions of interest based on the gross anatomy of tissue. Subsequently, excised pieces of tissue-hydrogel composite are processed using an in-gel digestion protocol, and recovered peptides are analyzed by liquid chromatography coupled with tandem mass spectrometry (LC-MS/MS). ProteomEx was applied to analyze subregion-specific proteome variability in healthy and pathological mouse brains. However, the spatial resolution of ProteomEx is currently limited to 160 µm due to challenges in manually handling small, transparent hydrogel samples during peptide recovery. Moreover, the sample preparation process typically requires approximately 54 hours (excluding MS data acquisition) and involves multiple manual steps, hindering the parallel processing of samples and limiting the feasibility of large-scale analyses. Mechanically tough tissues, such as the kidney and heart, which require more aggressive treatment for isotropic expansion, were not verified in the ProteomEx workflow. Furthermore, the super-resolution imaging capabilities of expanded samples were not realized in the ProteomEx method.

To address these limitations, we here report the development and validation of the enhanced ProteomEx method, referred to as microProteomEx, which allows 3D fluorescence super-resolution imaging of biological samples followed by MS-based proteome profiling of the targeted region of interest at a lateral resolution of 29-100 µm while retaining complete spatial information of tissue. Super-resolution imaging capabilities of microProteomEx were achieved using a conventional diffraction-limited microscope by staining expanded tissue samples with MS-analysis-compatible fluorescent dyes and/or commercial antibodies. Compared to the original protocol, the microProteomEx workflow is twice as fast and suitable for processing a wide range of mammalian tissues, including mechanically tough tissues such as the heart and kidney, due to optimized fixation and homogenization procedures. For instance, microProteomEx isolates single glomeruli from the kidney and micro-sized regions from heart tissue, yielding 3D structural imaging alongside proteomic data (∼600 protein identifications per ROI). We further validated microProteomEx on diverse pathological samples, including clinical archived melanoma tissues to distinguish molecular signatures between giant congenital melanocytic nevus (GCMN) and malignant melanoma (MM), and 5xFAD mouse brains to characterize molecular heterogeneity between morphologically distinct types of amyloid-β plaques. These applications demonstrate that microProteomEx uniquely combines multicolor 3D super-resolution imaging (down to 60 nm on conventional microscopes) with deep spatial proteomics at a lateral resolution of 29–100 µm. Compared to emerging platforms such as Deep Visual Proteomics (DVP)^20,21^ and Filter-Aided Expansion Proteomics (FAXP)^22^, microProteomEx offers superior nanoscale imaging capabilities and accessibility while maintaining competitive proteomic depth from micro-volume samples. Altogether, microProteomEx extends the expansion proteomics framework to super-resolution 3D imaging, opening up unprecedented capabilities for correlating proteome composition with nanoscale cytoarchitecture of intact tissue.

## Results

### Optimization of the ProteomEx workflow

We started by optimizing the original ProteomEx protocol to increase spatial resolution and streamline its workflow. The original ProteomEx protocol achieved a lateral/volumetric resolution of approximately 160 µm/0.6 nL before tissue expansion, corresponding to a 1 mm punch on a 30 µm tissue slice, magnified with a linear expansion factor (LEF) of ∼6. However, recovering peptides from expanded hydrogel samples smaller than 1 mm in diameter posed significant technical challenges that the original study did not overcome. This difficulty was attributed to the substantial shrinkage of hydrogel particles during peptide retrieval, caused by multiple buffer exchanges, which also resulted in the loss of approximately 10% of microdissected samples, as reported in the original study^19^. To address this shortcoming, we developed a re-embedding strategy in which excised hydrogel samples were embedded into a secondary, non-swelling polyacrylamide gel. This procedure effectively stabilized the hydrogel size during peptide recovery, thus enabling processing of samples smaller than 1 mm while preventing sample loss during in-gel digestion (**Supplementary Figure 1a**). To streamline sample processing of re-embedded samples, reduction and alkylation were performed on the entire expanded tissue slice before staining and microdissection, rather than on individual excised hydrogel pieces as in the original protocol (**Supplementary Figure 1b**). To evaluate the efficacy of peptide recovery from sub-nanoliter tissue samples via re-embedding, we used a laser capture microdissection (LCM) system to accurately isolate microscale tissue specimens. Correspondingly, expanded and Coomassie-stained mouse liver tissue slices were dehydrated without shrinkage (laser microdissection does not work on fully hydrated hydrogels), imaged with a bright field microscope, and dissected using LCM (**Supplementary Figure 1c**). Tissue fragments, with pre-expansion volumes of 0.0235 nL and 0.0975 nL, corresponding to approximately three and twelve hepatocytes^23^, respectively, were excised, re-embedded, and digested with trypsin (see Methods for details). MS analysis in data-dependent acquisition (DDA) mode identified 600/173 peptides/proteins from the 0.0235 nL sample and 4,493/1,064 peptides/proteins from the 0.0975 nL sample, with coefficients of variation (CVs) <17% (**Supplementary Figure 1d,e**). In conclusion, the re-embedding procedure enabled peptide recovery from sub-nanoliter tissue volumes that were 25 times lower than those of the original protocol, while preserving the integrity of the extracted peptides.

Encouraged by the ability to efficiently process sub-nanoliter volume samples via secondary re-embedding, we further optimized other critical steps of the original ProteomEx protocol to simplify and accelerate the sample preparation workflow. To minimize proteome variability arising from tissue-specific properties, mouse liver tissue was chosen for optimization due to its relatively low heterogeneity^24,25^. We found that N-Succinimidyl Acrylate (NSA) anchoring time, homogenization time, and Coomassie staining/destaining steps could be reduced by several-fold compared to the original protocol without compromising peptide or protein identifications^19^ (**Fig. 1a-c**). Specifically, the anchoring time was reduced by half, from 12 hours to 6 hours, the homogenization time was shortened by a factor of 8, from 12 hours to 1.5 hours, and the duration of the staining and expansion steps was decreased by 3.3-fold, from 10 hours to 3 hours (see Methods for details). These optimizations maintained high reproducibility and stability of protein identifications and quantification in microsamples (2.5 nL and 5.9 nL tissue volumes; R = 0.88-0.99), along with low CVs across different sample volumes (CVs of ∼16%), and Venn analysis demonstrated substantial overlap in identified proteins (**Supplementary Fig. 2a-c**). We also confirmed that workflow optimization resulted in an average distortion of 3.4% at a 1,250 µm scale for liver tissue (**Fig. 1d,e**), approximately half that of the original protocol^19^. In summary, we optimized the ProteomEx protocol by reducing the tissue volume suitable for efficient peptide recovery by re-embedding expanded samples in a non-swelling hydrogel, thus improving the achievable lateral resolution of microdissection from ∼160 µm to ∼28 µm. In addition, we streamlined and significantly accelerated the workflow, reducing the processing time by half (from 65.5 hours to 33 hours, excluding MS data acquisition; **Fig. 1f,g**) while improving the depth of proteome identification compared to the original protocol. Furthermore, peptide quality assessment revealed optimal length distributions (predominantly 7-20 amino acids) and reproducible digestion with ∼18-20% missed cleavage rates across both sample volumes (**Supplementary Fig. 2e,f**) Notably, protein identifications were improved, with an average of 4,054 proteins identified from 5.9 nL of mouse liver tissue (**Supplementary Fig. 2d**), compared to an average of 2,607 proteins identified from 5.6 nL of tissue in the original ProteomEx protocol^19^. We referred to the optimized protocol as microProteomEx (or µProteomEx for short) to emphasize its ability to efficiently process tissue microsamples.

**Figure 1.**
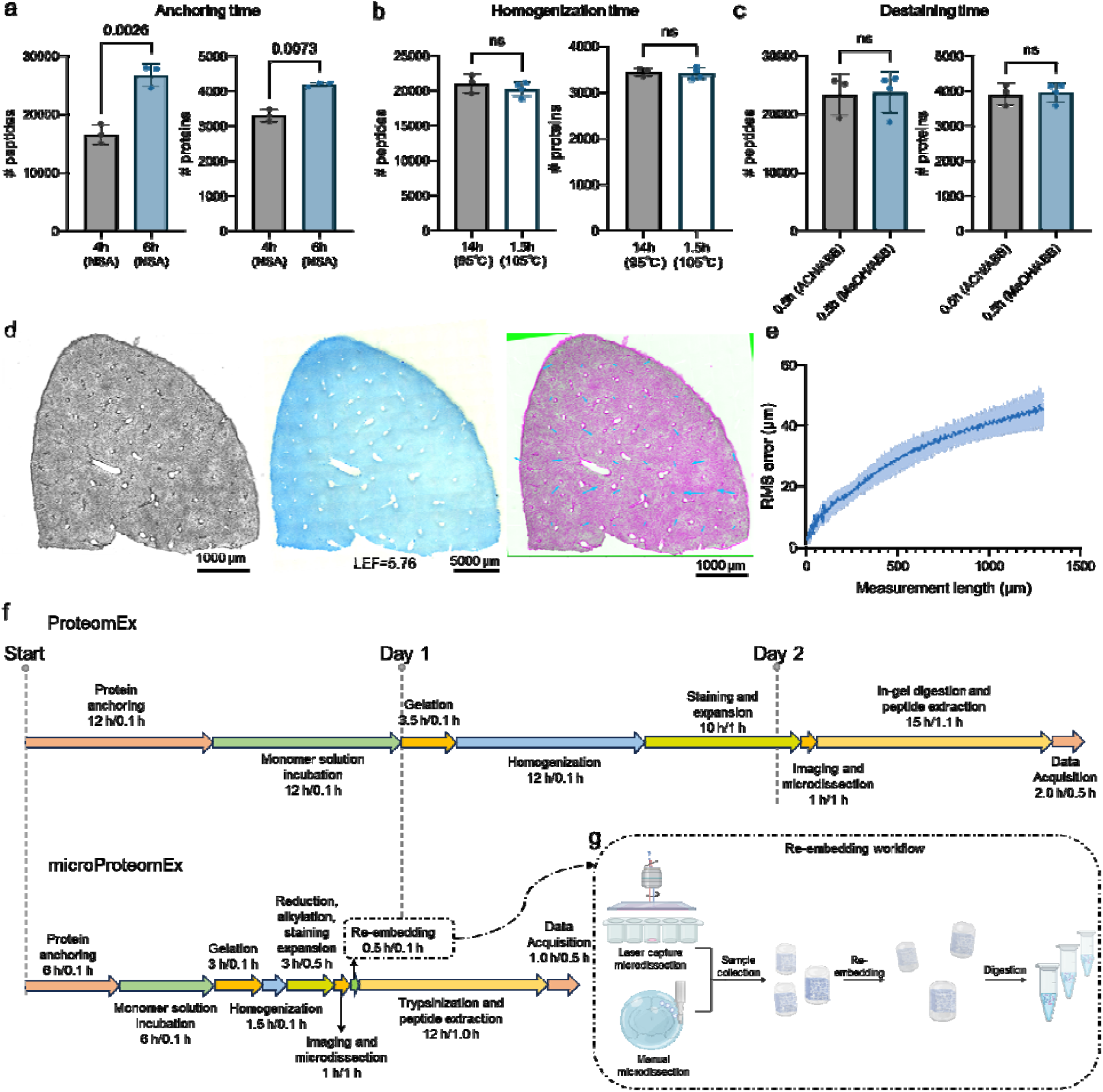
Acceleration of the ProteomEx workflow. (a) Number of peptide and protein identifications in 4 h and 6 h anchoring samples under 95□ homogenization condition and MeOH destaining condition (n = 3 punches per group from one slice each from one mouse; 5.9 nL of tissues per punch). Dot, individual data point; bar, mean; whiskers, standard deviation (SD), throughout Fig. 1a-c. Data in Fig 1a-c were analyzed by DDA. (b) Number of peptide and protein identifications in 95□ and 105□ homogenization condition samples under 6 h anchoring condition and MeOH destaining condition (n = 3 and 4 punches per group from one slice each for 95□ and 105□, respectively, from one mouse; 2.5 nL of tissues per punch). (c) Number of peptide and protein identification in 50% ACN/ 50% ABB and 50% MeOH/ 50% ABB destaining samples under 6 h anchoring and 95□ homogenization condition (n = 3 punches per group each from one mouse; 5.9 nL of tissues per punch). (d) Representative bright-field images of liver slice pre-expansion (left) and post-expansion (middle; Coomassie-stained) and overlay (right column) of pre-expansion image (green pseudo-color) and registered post-expansion image (magenta pseudo-color; n=3 slices from one mouse from one experiment). Arrows represent the deformation vector field, LEF=5.76. (e) The root-mean-square (RMS) measurement length error for pre- versus post-expansion liver slice images for the experiments shown in d (n=3 liver slices from one mouse from one experiment; blue line, mean; shaded area, SD; average LEF=5.92). (f). Comparison of the timeline of ProteomEx and microProteomEx, indicating total duration and hands-on time of each step (total duration/hands-on time). (g) Schematic workflow of re-embedding processing for manual and laser capture microdissection samples.

### Integration of super-resolution imaging into microProteomEx workflow

Physical magnification of biological samples enables 3D super-resolution fluorescence imaging under conventional diffraction-limited microscopes. However, this feature was not realized in the original study. To fully leverage the ProteomEx workflow, we utilized fluorescence staining of expanded tissue-hydrogel composites with MS-compatible small-molecule fluorescence dyes. We selected the Sypro series dyes due to their well-established use in PAGE-based proteomics and their specificity for labeling proteins under denaturing conditions^26–28^. Since the homogenization process involves SDS-mediated protein denaturation, these dyes are ideally suited for post-homogenization staining, without any risk of quenching the dyes during the polymerization or homogenization steps.

Since Sypro dyes had not previously been used in ExM, we first validated their staining patterns in liver tissue. For comparison, we chose NHS ester dyes, which are widely used for pan-protein staining of expanded samples in several ExM protocols^29–31^. Homogenized mouse liver samples were stained with SyproRed or Cy5-NHS ester, expanded, and imaged under a confocal microscope. Interestingly, the Cy5-NHS ester and SyproRed produced distinct labeling patterns at both the cellular and subcellular levels with high fluorescence contrast in standard near-infrared and red channels, respectively (**Supplementary Figure 3a,b**). The Cy5-NHS ester dye predominantly labeled nuclear and certain cytoplasmic proteins, whereas SyproRed primarily stained the cytoplasm while leaving the nucleus distinctly visible. Notably, SyproRed enabled clear delineation of cell boundaries, even under a relatively low-magnification objective (20×; **Figure 2a-c**), thereby facilitating the precise dissection of individual cells and even nuclei by LCM. We next compared the proteomic yield obtained from tissues stained with different dyes. Collected tissue-hydrogel composite pieces stained with SyproRed group yielded an average of 16,970/3,284 peptides/proteins per 3.23 nL of tissue, which was 1.96-fold (p = 0.004) and 1.42-fold (p = 0.017) higher than that from Cy5-NHS ester-stained samples per 4.04 nL of tissue (**Supplementary Fig. 3c**). Venn analysis demonstrated substantial overlap in the identified proteins, with less than 1% uniquely detected in the Cy5-NHS ester-stained group, indicating the enhanced identification rate in SyproRed-stained samples (**Supplementary Fig. 3d,e**). The significantly lower protein identifications observed in Cy5-NHS-stained samples can likely be attributed to the covalent modification of side-chain amino groups of lysine by the dye, as in the NHS-stained sample, we observed lower contribution of peptides with high numbers of Lys 3 and 4 and lower abundance compared to SyproRed-stained samples (**Supplementary Fig. 3f,g,h**)^32,33^. In contrast, Sypro dyes do not covalently modify proteins, thereby minimizing interference with subsequent MS identifications. SyproRed was selected as the preferred fluorescent dye for proteomic analysis of expanded tissue samples due to its superior protein yield and consistent staining pattern.

**Figure 2.**
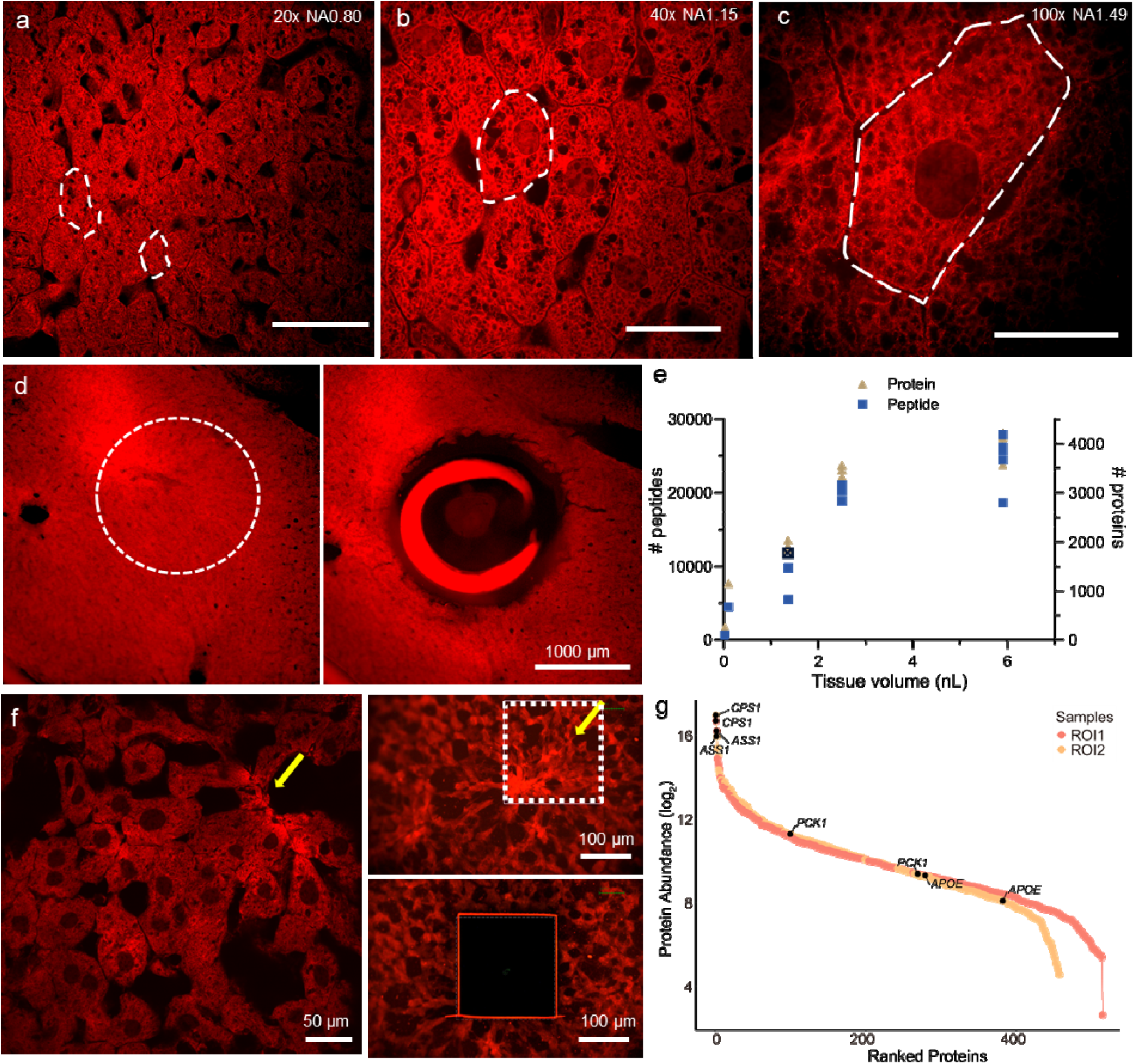
Combination of super-resolution imaging of mouse liver tissue with microProteomEx. (a-c) Representative confocal images (SyproRed-stained) of post-expansion mouse liver tissue. LEF = 5.26 in a-c. Effective spatial resolution of imaging: 76.7 nm in a, 58.4 nm in b, 41.2 nm in c. Scale bars, 9.5 μm (a-c (physical size post-expansion, 50 μm). (d) Fluorescence images of manual dissection mouse liver ROI (right, before microdissection; left, after microdissection; white circle indicated ROI selected for microdissection; left, after microdissection). (e) Number of peptide and protein identification for varied tissue volumes of mouse liver tissues. Black data point represents the sample in e (analyzed by DDA mode). (f) Confocal image and LCM pre- and post-cut images of mouse liver ROI. (g) Rank intensity of identified proteins from LCM samples. Analyzed by DIA mode.

In mouse liver tissue, with a routinely achieved LEF of ∼5.3, we attained effective lateral resolutions of 76.7 nm under 20x NA0.8 objective, 58.4 nm (40x, NA1.15), and 41.2 nm (100x, NA1.49) on a confocal microscope (effective resolution was calculated as theoretical resolution divided by LEF; **Fig. 2a-c**; **Supplementary Movie 1**). Next, LC-MS/MS analysis was performed on the stained and microdissected samples. After whole-slice imaging, regions of interest (ROIs) were selected based on tissue/cellular morphology. These ROIs were highlighted using an excitation beam from the inverted microscope’s objective lens to guide precise manual microdissection, with images captured before and after dissection to confirm precise excision (**Fig. 2d**; see **Supplementary Fig. 4** for more representative images). The number of identified peptides and proteins increased with tissue volume, reaching a plateau between 3.0-6.0 nL tissue size (or 480 µm of lateral resolution; **Fig. 2e**). In addition, SyproRed staining retained robust fluorescence and consistent patterns in dried samples, facilitating fluorescence-guided microdissection under LCM (**Fig. 2f**). This enabled the identification of approximately 500 proteins from 0.4 nL, corresponding to approximately 50 hepatocytes^23^, of expanded mouse liver tissue (**Fig. 2g**). Among the identified proteins were key liver metabolic enzymes, including PCK1 (phosphoenolpyruvate carboxykinase 1, critical for gluconeogenesis), CPS1 (carbamoyl-phosphate synthase 1, essential for urea cycle function), ASS1 (argininosuccinate synthase 1, involved in arginine biosynthesis), and APOE (apolipoprotein E, a regulator of lipid metabolism; **Fig. 2g**). In summary, integrating fluorescence imaging into the microProteomEx workflow provides an effective strategy for microscale tissue proteomics, enabling precise, imaging-guided targeting of ROIs based on ultrastructural features.

### Optimizing microProteomEx for mechanically tough tissues

To broaden the applicability of ProteomEx to diverse mammalian tissue types, we extended its use to mechanically tough tissues, such as the kidney and heart. However, application of the standard protocol to mouse heart and kidney tissues resulted in significant distortion and cracking post-expansion, possibly due to a higher extracellular matrix content (**Supplementary Fig. 5a,b**). The common approach to enhancing isotropic expansion is to increase the temperature or duration of homogenization, as well as add specific proteases (*e.g.*, collagenase)^34^. However, this may compromise protein identification due to increased protein degradation or hydrolysis. An alternative strategy involves reducing cross-linking during fixation, as previously employed in techniques such as U-ExM^35^. This can be easily achieved by reducing the PFA concentration while simultaneously incorporating protein-anchoring agents during fixation. Correspondingly, we reduced the paraformaldehyde (PFA) concentration during tissue fixation from the standard 4% to 1% and supplemented the fixation solution with the NSA protein anchor. Furthermore, pH of the homogenization buffer was adjusted to be slightly acidic (pH∼5) to prevent premature expansion before the tissue-hydrogel composite gets fully homogenized. We began by evaluating the effects of different fixation methods on proteomic analysis by comparing the proteomic output of 1% PFA and 1% PFA + NSA with that of 4% PFA, using mouse liver tissue to align with previous optimizations. Proteomic analysis of ∼5.9 nL tissue volumes revealed that the 1% PFA + NSA fixation exhibited the highest peptide/protein identifications with a nearly 3-fold improvement in the number of peptides/proteins identified compared to the 4% PFA samples under identical processing conditions (**Fig. 3a**). A significant fraction of proteins (2,559, 56%) was identified across all three fixation conditions (**Fig. 3b**). Additionally, 1,244 proteins (27%) were identified in both the 1% PFA and 1% PFA + NSA groups. These results suggest that although the 1% PFA + NSA fixation method does not substantially alter the composition of identified proteins, it improves overall protein retention. The CV analysis indicated similar variability in protein abundance between three groups (∼12-16%; **Supplementary Fig. 5c**). Moreover, the 1% PFA + NSA fixation method demonstrated higher reproducibility between biological replicates at both 2.5 nL and 5.9 nL tissue volumes, with correlation values ranging from 0.84 to 0.98 (**Supplementary Fig. 5d,e**). The subcellular localization and functional classification of identified proteins across three fixation methods showed that the 1% PFA and 1% PFA + NSA groups had a higher proportion of nuclear proteins compared to the 4% PFA group, suggesting fixation-induced biases, while the overall distribution of functional protein categories remained consistent, indicating that 1% PFA + NSA preserves a diverse range of functional proteins (**Supplementary Fig. 5f,g**). Selecting the 1% PFA + NSA fixation method due to the highest proteomics yield, we conducted the isotropic analysis on the kidney and heart confirming excellent preservation of tissue gross anatomy during tissue expansion up to a LEF of ∼6, with RMS length measurement errors of ∼2.4% and ∼3.0% for heart and kidney tissues, respectively, at length scales up to 1,250 μm (**Fig. 3d-g**). These findings indicate that the 1% PFA + NSA fixation method effectively preserved morphological integrity, enabling the application of the protocol to mechanically resilient tissues such as the mouse heart and kidney.

**Figure 3.**
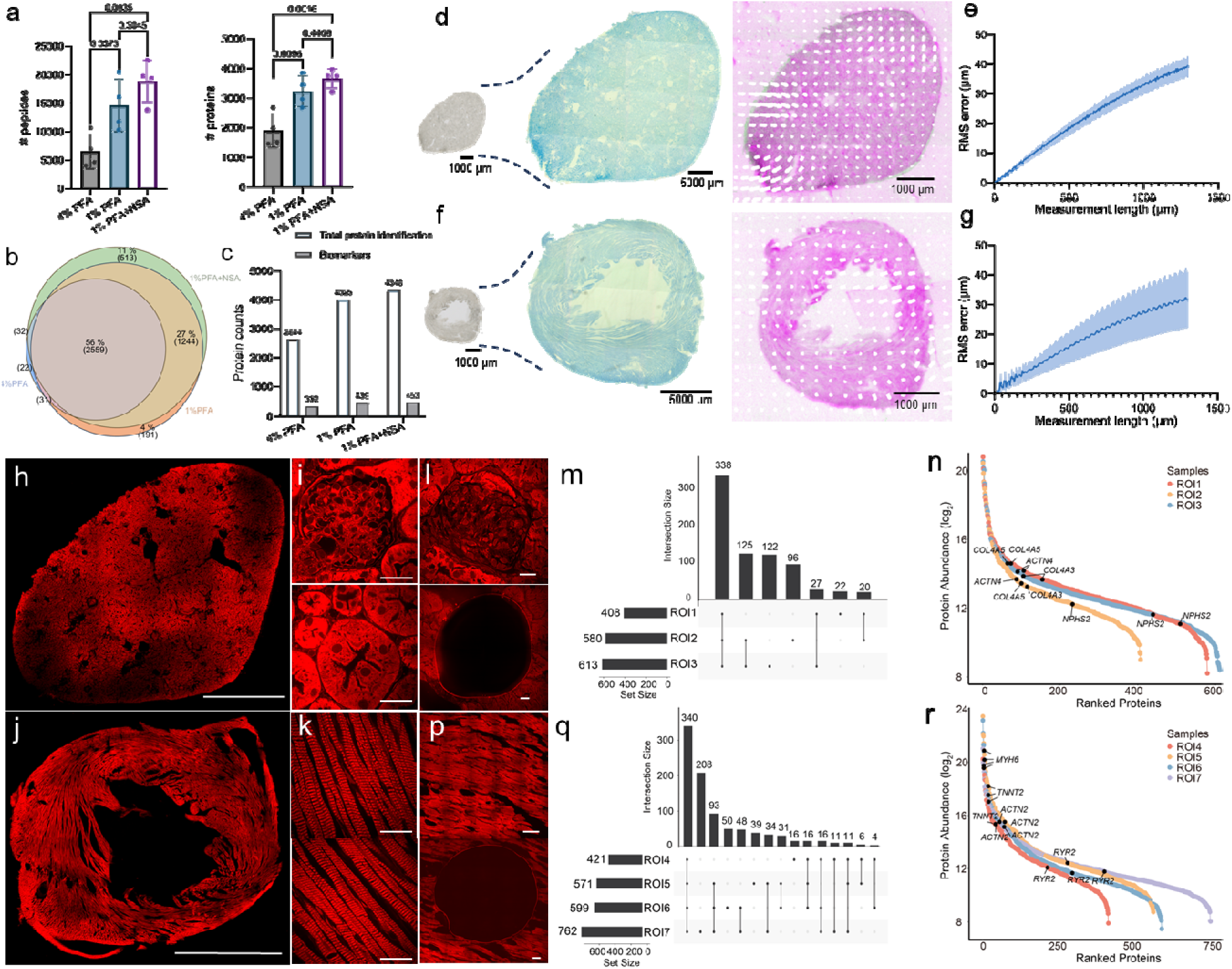
The optimized fixation protocol for the isotropic expansion of tough tissues using microProteomEx. (a) Number of peptide and protein identifications in 4% PFA, 1% PFA, and 1% PFA + NSA fixed mouse liver samples (n = 4 punches per group from one slice each from one mouse; ∼5.89 nL of tissues per punch; data were analyzed by DDA). (b) A Venn diagram of the total proteins identified for the samples is shown in a. (c) The total number of protein identifications and biomarkers for the samples shown in a. (d) Representative bright-field image of mouse kidney slice pre-expansion (left) and post-expansion (middle; Coomassie-stained) and overlay (right column) of pre-expansion image (green pseudo-color) and registered post-expansion image (magenta pseudo-color; n=3 slices from 2 mice from 2 independent experiments). Arrows represent the deformation vector field, LEF= 5.93. (e) The RMS measurement length error for pre- versus post-expansion kidney slice images for the experiment shown in a (n=3 slices from 2 mice from 2 independent experiments; blue line, mean; shaded area, SD; average LEF=6.11. (f) Representative bright-field images of mouse heart slice pre-expansion (left) and post-expansion (middle; Coomassie-stained) and overlay (right column) of pre-expansion image (green pseudo-color) and registered post-expansion image (magenta pseudo-color; n=3 slices from one mouse from one experiment). Arrows represent the deformation vector field; LEF=5.93. (g) The RMS measurement length error for pre- versus post-expansion heart slice images for the experiments shown in a (n=3 slices from one mouse from one experiment; blue line, mean; shaded area, SD; average LEF=5.42). (h-i) Representative confocal single-plane fluorescence images of expanded kidney tissue stained with SyproRed showing glomerulus (left) and renal tubules (right; n = 3 slices from 1 mouse; imaging under 40x NA1.15 at effective lateral resolution of LEF = 5.79). (j-k) Representative confocal single-plane fluorescence images of expanded heart tissue stained with SyproRed (n = 3 slices from 1 mouse; imaging under 40x NA1.15 at effective lateral resolution of LEF = 5.36). (l) Confocal image and LCM pre- and post-cut images of mouse glomeruli ROI. (m) The number of shared and unique sets of proteins in **3** ROIs from kidney tissue. (n) Rank order of core protein abundance in 3 ROIs from kidney tissue. (o) Confocal image and LCM pre- and post-cut images of mouse heart tissue ROI. (p) The number of shared and unique set of proteins in 4 ROIs from heart tissue. (q) Pearson correlation among ROIs from heart tissue. (q) Rank order of core protein abundance in ROIs from heart tissue. Scale bars: (physical size post expansion) 5000 μm in h and j; 100 μm in i, l, k and p.

Then we employed super-resolution imaging to evaluate nanoscale structures under the optimized fixation conditions. SyproRed staining confirmed excellent preservation of tissue gross anatomy during expansion up to a LEF of ∼6 (**Fig. 3h-k**). High-NA confocal imaging resolved well-defined nanoscale morphological details, including individual glomeruli and tubular structures in the mouse kidney (**Fig. 3i**) and intact myocytes with clearly delineated sarcomeres in heart tissue (**Fig. 3k**), enabling super-resolution 3D rendering of a single glomerulus and myocyte structures (**Supplementary Movies 2–3**). Following super-resolution imaging, we isolated three individual glomeruli, about 0.278 nL of tissue each, from expanded mouse kidney slices using manual dissection with biopsy punch or LCM (**Fig. 3l; Supplementary Fig. 6**). These glomeruli yielded approximately 600 protein identifications each, sharing 338 proteins defining a core proteome of mouse glomeruli (**Fig. 3m**). Several glomeruli-associated markers were among the identified proteins, including NPHS2 (podocin, a key component of the glomerular filtration barrier), COL4A3 (collagen type IV alpha 3 chain, essential for glomerular basement membrane integrity), COL4A5 (collagen type IV alpha 5 chain, involved in basement membrane structure), and ACTN4 (alpha-actinin-4, a cytoskeletal protein critical for podocyte function; **Fig. 3n**). Similarly, we isolated four ROIs, about 1.642 nL of tissue each, from expanded mouse heart samples (**Fig. 3p; Supplementary Fig. 6**), identifying in total 924 proteins, with 338 shared proteins (45.1% of detected proteins) defining a core proteome of mouse heart tissue (**Fig. 3q**). Identified proteins included those involved in core cardiac pathways, such as MYH6 (myosin heavy chain 6, the alpha isoform predominant in cardiac muscle contraction), TNNT2 (troponin T type 2, a regulator of cardiac muscle thin filament function), RYR2 (ryanodine receptor 2, mediating calcium release in cardiomyocytes), and ACTN2 (alpha-actinin-2, a sarcomeric cross-linking protein essential for myofibril organization; **Fig. 3q**). Pearson correlation analysis showed strong correlations between each pair individual microsamples for kidney and heart tissues (0.80–0.92; **Supplementary Fig. 6d,e**). These results demonstrated that microProteomEx enables the correlation of nanoscale imaging of tissue cytoarchitecture with corresponding proteomic information, providing a novel method for investigating ultrastructural-molecular heterogeneity in biological systems.

### Application of microProteomEx to archived clinical samples

Having successfully applied microProteomEx to mechanically tough tissues, such as heart and kidney, we extended this method to archived clinical skin samples, which exhibit similar mechanical properties due to fibrous and collagen-rich structures. These structures may also pose significant hurdles for proteomic analysis due to their resistance to homogenization and protein extraction^36,37^. We selected formalin-fixed paraffin-embedded (FFPE) skin samples from a pediatric patient diagnosed with malignant melanoma (MM) arising in a giant congenital melanocytic nevus (GCMN)^38^. This clinical case represents an exceptionally rare pathological entity^39^. GCMN occurs in approximately 1 in 20,000 newborns, with malignant transformation to melanoma documented in fewer than 0.5% of cases within the first year of life^40^. This unique case enabled us to focus on the stage transition between two distinct regions within the same lesion, specifically GCMN and MM, allowing for direct comparative proteomic analysis without inter-patient or inter-tissue variability.

Skin slices from the patient’s lesion were collected and stained with hematoxylin and eosin (H&E) for annotation by expert pathologists (**Fig. 4a,b**). The slices were designated as containing primarily GCMN or MM regions based on histopathological features. Two adjacent FFPE slices were processed using the microProteomEx workflow, optimized for tough tissue as described above, which included dewaxing, protein anchoring, gelation, homogenization, fluorescent staining, expansion, imaging, manual microdissection, and MS analysis (**Fig. 4a,c,d**). After expansion, we performed 3D super-resolution imaging of the cytoarchitecture in both regions, visualized with SyproRed staining, which revealed significant cytostructural differences between the MM and GCMN regions (**Fig. 4e** and **Supplementary Movies 4** and **5** for GCMN and MM, respectively). The GCMN area exhibited a more organized, nested cellular arrangement typical of benign nevi, while the MM region showed disrupted architecture with clustered cellular density and irregularity. These morphological differences align with established histological features, including the structured and adnexal involvement in GCMN^41,42^ and the atypical, invasive proliferation in MM^43,44^.

**Figure 4.**
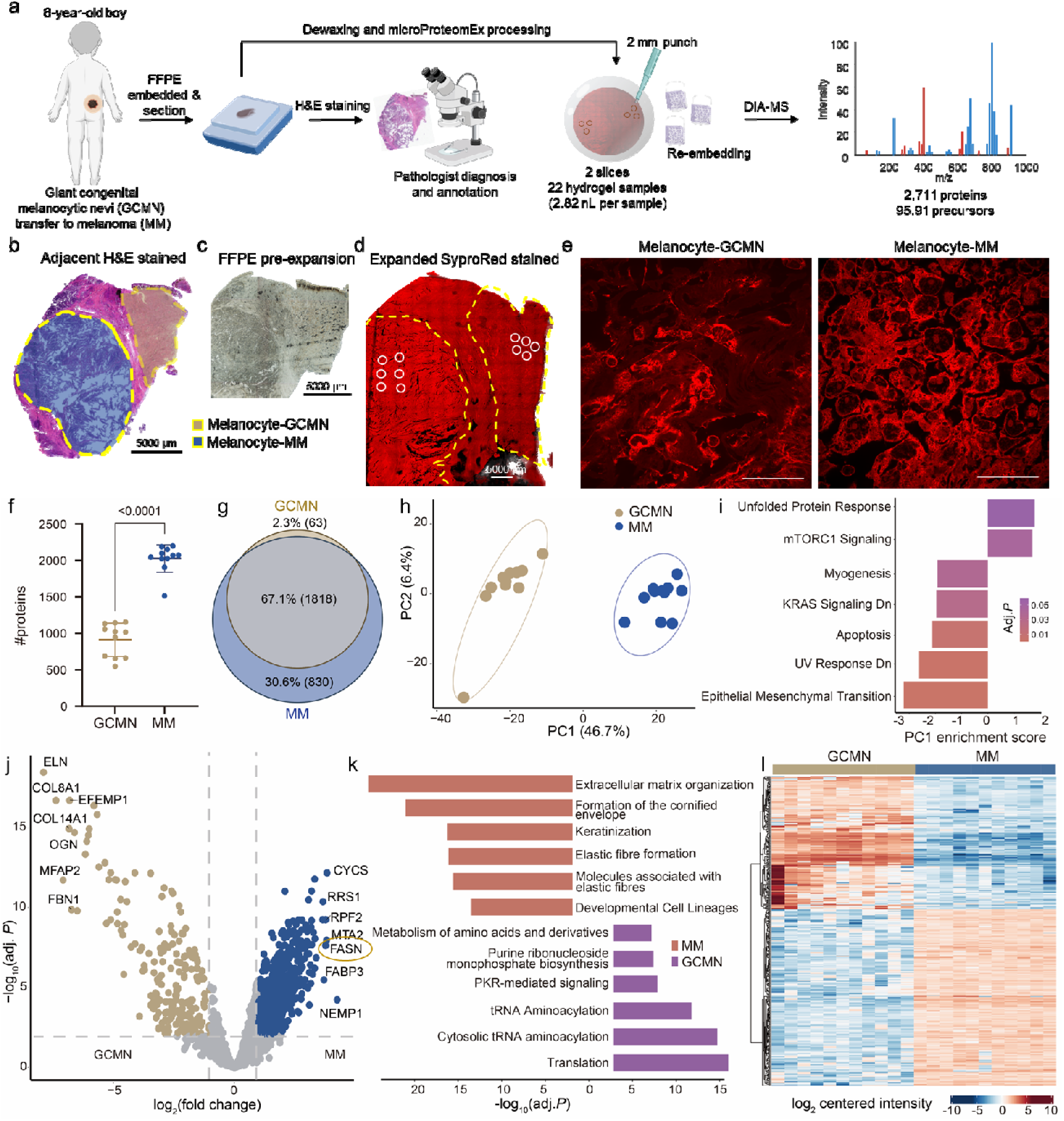
Application of microProteomEx for archived FFPE clinical skin samples. (a) Schematic workflow of sample processing. (b) Bright-field image of H&E-stained skin sample annotated by an expert pathologist, indicating regions of melanocyte-GCMN and melanocyte-MM tissue. (c) pre-expansion Bright-field image of the FFPE skin slice, adjacent to the slice shown in b, used for microProteomEx analysis. (d) Post-expansion single-plane confocal fluorescence image of SyproRed-stained skin sample showing excised ROIs indicated by white circles. (e) Representative super-resolution images of melanocyte-GCMN and melanocyte-MM from the expanded skin sample shown in d (n= 5 FOVs from GCMN and MM from 2 slices). Physical size post-expansion, 50 μm; LEF = 3.65 in d and e. (f) Number of proteins identified in melanocyte-GCMN and melanocyte-MM tissue. *n*□=□11 individuals per group. Analyzed by an unpaired t-test with Welch’s correction. (g) The overlap of identified proteins in two groups. (h) Principal component (PC) analysis of 22 skin samples. (i) Gene set enrichment analysis (GSEA) (MSigDB Hallmark) of PC1 component shown in (h). (j) Volcano plot indicating proteins upregulated in MM and GCMN regions. Claybank and blue colors represent proteins with BH adjusted *P* value (< 0.01), and more than 2-fold change. Unsignificant proteins are colored in gray. (k) Enriched Reactome pathways for differential proteins shown in (j). Red bars indicate pathways enriched in the downregulated proteins in MM region. Purple bars indicate pathways enriched in the downregulated proteins in GCMN region. (l) Heatmap showing the 338 significantly differentially expressed proteins (DEPs) across MM and GCMN regions. Adj. *P*, adjusted *P* value.

Following imaging, manual dissection was performed using 2 mm punches (corresponding to a ∼549 µm spot in diameter or ∼1.1 nL tissue volume), yielding 11 independent samples per region, each analyzed separately in DIA mode on the timsTOF HT mass spectrometer. The analysis identified approximately 1,881 proteins in the GCMN samples and about 2,648 proteins in the MM samples, with 990 ± 236 and 2,212 ± 209 proteins identified per sample in GCMN and MM, respectively, and 1818 shared proteins (**Fig. 4f, g**). Both GCMN and MM regions showed comparable technical reproducibility assessed by calculating Pearson correlation coefficients and CV for protein quantification across samples (**Supplementary Fig. 7**). Notably, 96.7 of proteins identified in GCMN were also present in MM, indicating a shared core proteome alongside an expanded protein repertoire in MM consistent with melanoma progression, which has enhanced metabolic and proliferative demands in melanoma progression.

Principal component analysis (PCA) of the proteomic data clearly separated the GCMN and MM groups, with PC1 accounting for approximately 46.7% of the variance, highlighting distinct proteomic profiles between the two stages (**Fig. 4h**). To further interpret these differences, we performed gene set enrichment analysis (GSEA) on proteins contributing to PC1 using the Hallmark database (**Fig. 4i**). Enrichment analysis revealed that mTORC1 signaling was significantly enriched in MM, reflecting enhanced anabolic and proliferative signaling. In contrast, epithelial-mesenchymal transition (EMT), myogenesis, apoptosis, UV response, and KRAS signaling were all significantly enriched in GCMN, indicating that these programs are relatively suppressed in MM compared to GCMN. These findings reflect the loss of stromal and structural identity in MM relative to the organized nevus tissue of GCMN, as well as enhanced translational and metabolic activity in the malignant stage.

Differential expression analysis identified 197 upregulated and 141 downregulated proteins in MM versus GCMN (**Fig. 4j**). Among the top differentially expressed proteins was fatty acid synthase (FASN), which was upregulated in the advanced MM stage. FASN is a key enzyme in lipid metabolism, catalyzing *de novo* fatty acid synthesis, and its overexpression is well documented in melanoma, where it supports membrane biogenesis and energy demands during rapid proliferation^45–47^. The most strongly downregulated proteins in MM included elastin (ELN), COL8A1, MFAP2, COL14A1, and EFEMP1, all of which are components of the extracellular matrix and elastic fiber network. Reactome pathway enrichment of the downregulated proteins revealed that extracellular matrix organization, formation of the cornified envelope, keratinization, elastic fiber formation, and molecules associated with elastic fibers were all significantly enriched among proteins downregulated in MM (**Fig. 4k**), reflecting a broad collapse of the structural ECM and skin differentiation programs that characterize the benign nevus. Conversely, pathways enriched among upregulated proteins in MM included translation, tRNA aminoacylation, and amino acid metabolism, consistent with the elevated biosynthetic demands of malignant proliferation (**Fig. 4k**). Hierarchical clustering of the proteomic data further confirmed clear separation between samples from the GCMN and MM regions, with consistent intra-group clustering as expected (**Fig. 4l**).

To further explore the biological significance of the acquired proteomic dataset, we utilized Kosmos^48^, a novel AI agent that automates data-driven discovery, to identify key proteins and pathways driving malignancy and propose mechanistic hypotheses. After five iterations of analysis, each consisting of 3-5 individual data analysis or literature search tasks, Kosmos synthesized four key discoveries (**Supplementary Data 1**). The results highlighted a proteome-wide collapse of the extracellular matrix pathway (GO:0062023, adj p=5.90x10, **Supplementary Dataset 1**), consistent with the ECM downregulation identified in our independent bioinformatic analysis (**Fig. 4k**). Among other Kosmos’s discoveries worth mentioning is a L1-regularized logistic regression (LASSO) analysis of the proteomic data, which uncovered a small 13-protein panel. This panel comprises 12 downregulated proteins involved in the extracellular matrix and cell adhesion, including PRELP, COL3A1, COL14A1, and GPNMB, as well as one upregulated chaperone, HSP90AB1. This panel achieved perfect separation of MM from GCMN samples (**Supplementary Data 1**), with all panel members occupying central hub positions in the dysregulated protein interaction network. In addition, Kosmos identified a notable reprogramming of immune pathways, specifically a coordinated downregulation of proteins associated with innate immune activity and neutrophil-mediated immunity (GO:0002446, adj p=1.48x10 ¹). Kosmos went further, proposing mechanistic hypotheses based on network analysis and a literature search, linking the observed loss of cell adhesion to a compensatory stress-adaptation program coordinated by HSP90AB1 via shared focal adhesion partners.

To validate these additional Kosmos-generated findings, we acquired orthogonal experimental data using chromatin immunoprecipitation sequencing (ChIP-seq) for the repressive H3K27me3 mark to investigate gene expression suppression. Fingerprint and correlation analyses confirmed broad H3K27me3 enrichment and high reproducibility between samples (**Supplementary Fig. 8a,b**). Genome browser inspection of representative downregulated genes, including *ELN* and *EHD2*, revealed elevated promoter-associated H3K27me3 signal in MM relative to GCMN (**Supplementary Fig. 8c**); ELN is involved in elastic fiber assembly and extracellular matrix integrity, while EHD2 regulates membrane dynamics and cell migration, both potentially processes relevant to melanoma progression. Metagene analysis centered on TSSs (±3 kb) indicated that downregulated proteins showed elevated H3K27me3 signals in MM relative to GCMN (**Supplementary Fig. 8d**). These results suggest that promoter-associated H3K27me3 contributes to transcriptional repression of a subset of downregulated proteins, providing orthogonal support for the proteomics findings. This validation dataset was input into Kosmos with the same research objective to compare MM and GCMN. Kosmos’s analysis of the ChIP-seq data independently converged on the same core themes (**Supplementary Data 2**). It identified promoter-centric H3K27me3 reprogramming, an epigenetic silencing mechanism, as the driver for the collapse of cell adhesion networks, including “Focal Adhesion” (GO:0005925, adj p=9.79x10□□) and “Cell-Substrate Junction” (GO:0030055, adj p=9.26x10□□). Results also suggested reprogramming of immune pathways, characterized by the loss of H3K27me3 at promoters (derepression) of pro-inflammatory cytokines and chemokines (e.g., IL1B, IL18, CCL3, CCL4, CCL20), effectively activating a pro-inflammatory microenvironment. The independent confirmation of these complex biological themes across two orthogonal datasets demonstrates Kosmos’s utility beyond standard data analysis methods, offering a broad range of plausible hypotheses for further experimental exploration.

### Single-Plaque Proteomics and Proteomic Profiles of Different Types of Amyloid Plaques in 5xFAD Mouse Model

Having demonstrated that microProteomEx enabled straightforward proteomic profiling guided by super-resolution imaging across diverse tissue types, we next applied this approach to characterize micro-pathological structures in an AD mouse model. One of the most significant morphological changes in AD is the accumulation of amyloid-beta (Aβ) protein deposits. The size of individual Aβ plaques is typically 30-100 µm^49,50^ in diameter, which falls within the lateral resolution achieved with microProteomEx. To facilitate image-guided microdissection of Aβ plaques in expanded brain tissue, we first tested the compatibility of antibody staining with microProteomEx protocol. By using the 5xFAD mouse model of AD, we confirmed that anti-Aβ aa 1-16 antibodies could efficiently stain post-expansion samples, producing patterns identical to those observed with conventional immunohistochemistry of pre-expansion samples (**Supplementary Fig. 9**). In addition, we noted that SyproRed staining of expanded brain tissue facilitated visualization of distinct neuroanatomical features across various brain regions (**Supplementary Fig. 10**). Major brain regions, including the cortex (i), striatum (ii), corpus callosum (iii), and anterior commissure (iv), were clearly visualized in expanded brain tissue using SyproRed staining (**Supplementary Fig. 10**). The high spatial resolution allowed for the observation of intricate protein distributions across these regions, providing valuable insights into the brain’s cellular architecture. As a result, confocal imaging of the expanded samples enabled super-resolution 3D imaging of Aβ plaques within the intact tissue context, with SyproRed clearly delineating neuropil structures in the cortex and DAPI highlighting cell nuclei (**Fig. 5a-e**, **Supplementary Movie 6-7**).

**Figure 5.**
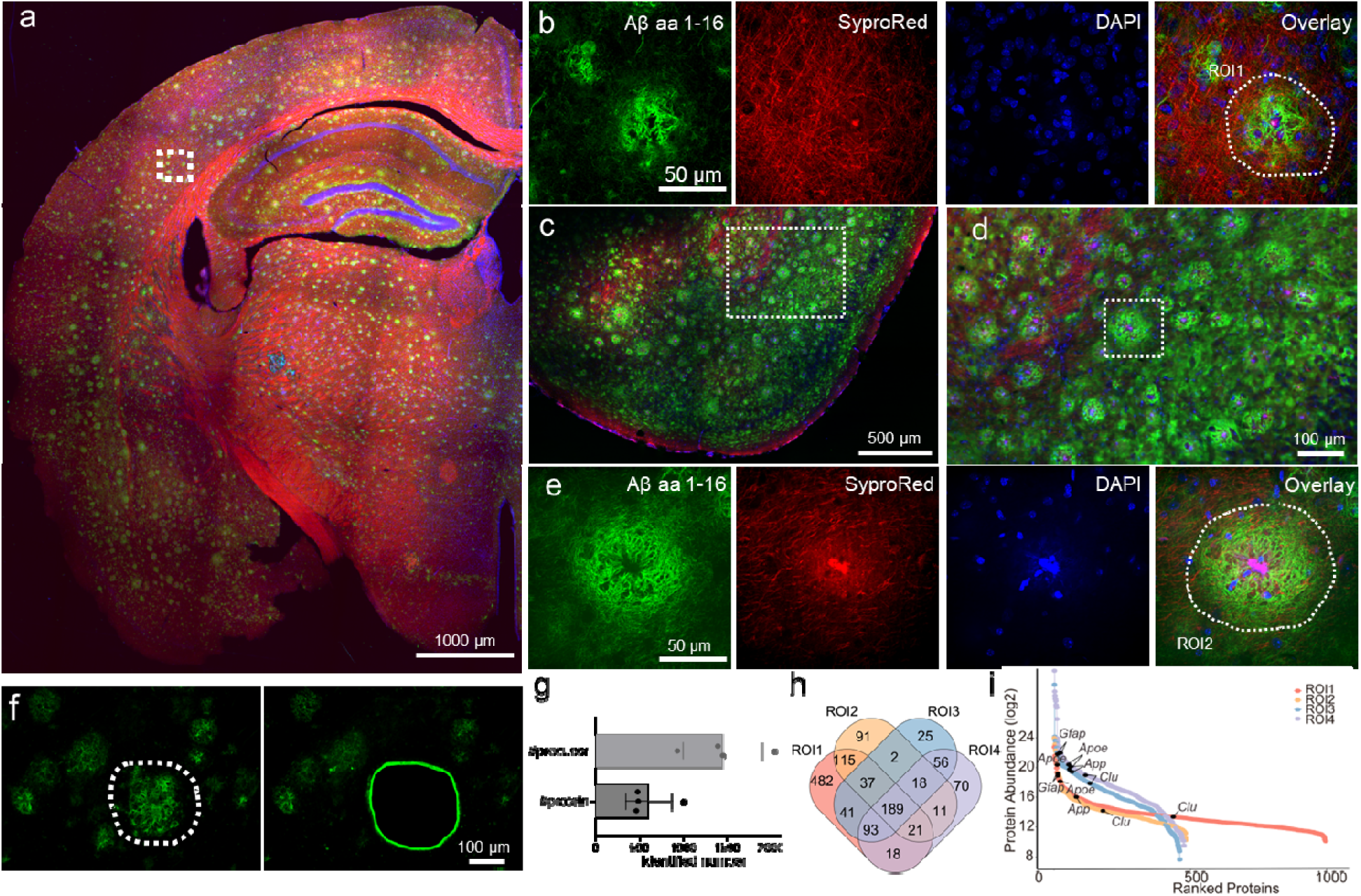
Visual spatial proteomic profiling of amyloid plaque in 5x FAD model. AD brain samples were expanded and stained with DAPI, Sypro Red, and an antibody against the Aβ 1-16 peptide. Following super-resolution imaging to capture individual plaques, targeted regions were microdissected using LCM, and the corresponding peptide samples were analyzed by MS. (a) Merged composite image of the left hemisphere of the AD mouse brain. The boxed region denotes ROI 1 corresponding to a single plaque. (b-c) Merged composite images of the left hemisphere of the AD mouse brain. c, the magnified view of the region captured in b. The boxed region marks ROI 2, which is shown in greater detail in panel (d). (d) Single-channel images of each staining modality. The circular region indicates the LCM-targeted area for ROI2. (e) LCM images before and after microdissection of the targeted ROI2. (f-h) Proteome of ROIs is acquired by DIA mode and analyzed by DIA-NN. f: Number of identified precursors and peptides for ROI 1 and 2 proteomes. g: Rank plot showing protein intensity for proteomes of ROI 1 and 2. AD related protein and ranked highest and lowest 3 proteins are labeled in plot. h: Venn diagram depicting the overlap of identified proteins between ROI 1 and ROI 2 proteomes. (i) study design of proteomic analysis of two types amyloid plaque in 5x FAD model. (j) High-resolution images of diffuse plaque and classic cored plaque in a 10-month 5x FAD mouse brain cortex. (k) Upset diagram showing the identification depth and overlap for each group. (l) Principal component analysis showing the sampl clusters based on the prototype. Brightness and contrast settings: each channel was first set by the automatic adjustment function in Nikon Elements analysis, and then manually adjusted to improve contrast. LEF: 2.5; Scale bars labeled in Figure are in expansion units. Effective spatial resolution of imaging: 113 nm in d and j.

Next, using the LCM method, we precisely isolated four selected ROIs, each corresponding to a single amyloid plaque, from the left-hemisphere cortex and subjected them to MS-based analysis (**Fig. 5a-f**). Across the four ROIs, approximately 450–1000 proteins were identified, among which 189 proteins were consistently detected across all four ROIs (**Fig. 5g, h**). A rank plot of protein intensities highlighted the relative abundance of key AD-related proteins in the following order glial fibrillary acidic protein (GFAP), Apolipoprotein E (ApoE), amyloid-beta precursor protein (APP), and clusterin (Clu) (**Fig. 5i**). Further functional categorization of the top abundant proteins consistently detected across all four ROIs revealed diverse cellular processes, including cytoskeletal components (β-actin/γ-actin (Actb; Actg1), α-tubulin 4a (Tuba4a), and β-tubulin 5 (Tubb5), cell adhesion molecules (desmoplakin (Dsp) and annexin A2 (Anxa2), glycolytic enzymes (Gapdh, Eno1, Pkm), mitochondrial proteins (Atp5f1a, Slc25a4, Ogdh), elongation factors (Eef2, Eef1a1) and ribosomal/ubiquitin proteins (Rps27a;Uba52;Ubb;Ubc), collectively indicating metabolically active tissue with robust structural organization. In summary, these results demonstrate that microProteomEx, integrated with super-resolution fluorescence imaging, enabled the precise identification, isolation, and untargeted proteomic profiling of individual amyloid plaques in the 5xFAD AD mouse model.

During super-resolution 3D imaging of the cortex regions of the 5xFAD mouse brains stained with SyproRed, DAPI, and anti-Aβ aa 1-16 antibodies, we observed two morphologically distinct types of plaque depositions. The first type, classified as a cored plaque (also known as a dense core or classic cored plaque)^51–53^, displayed a well-defined, compact Aβ core with radially oriented fibrillar structures, accompanied by a concentrated SyproRed signal, indicative of dense protein aggregation. In contrast, the second type of plaques, which we referred to as diffuse plaques (aka coarse-grained plaque)^54–56^, lacked a compact core, showing a more dispersed and loosely organized Aβ deposition pattern with less prominent SyproRed colocalization, reflecting an early and less structured stage of amyloid accumulation (see **Supplementary Figure 11** for representative images and clear labeling of the described structures).

To leverage the unique ability of image-guided classification and isolation of distinct Aβ plaque types, we investigated proteomic differences between cored and diffuse plaques at two stages of AD pathology progression: moderate AD pathology (5-month-old female 5xFAD) and late-stage disease (10-month-old female 5xFAD). Six brain slices were processed using microProteomEx workflow, including post-homogenization plaque immunostaining, super-resolution 3D imaging via confocal microscopy, image analysis, and microdissection of the selected plaques using LCM, followed by re-embedding and peptide retrieval for MS data acquisition in DIA mode (**Fig. 6a**).

**Figure 6.**
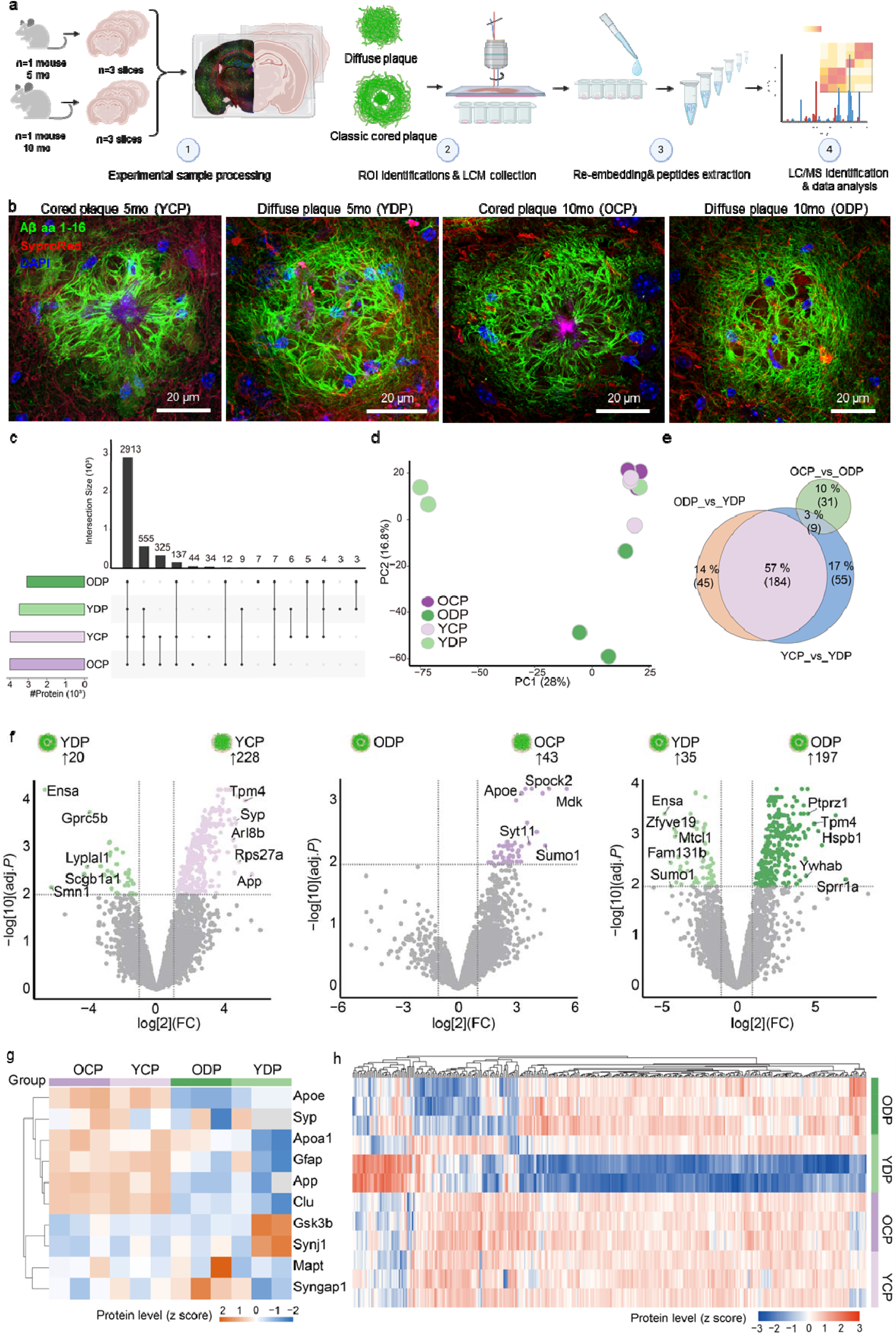
Spatial proteomic profiling of two types of amyloid plaque in 5-month-old and 10-month-old 5x FAD mice. (a) Study design of proteomics analysis of two types of amyloid plaque in the 5x FAD model. (b) High-resolution images of diffuse plaque and classic cored plaque in 10-month 5x FAD mouse brain cortex. (c) Upset diagram showing the identification depth and overlap for each group. (d) number of protein intensity across all samples. (e) Principal component analysis showing the sample clusters based on the prototype. (f) Multi-group volcano plot showing the differential protein expression analysis results for four groups: YDP, YCP, ODP, and OCP. (g) Heatmap displaying z-score normalized protein levels for a representative set of differentially enriched proteins across four plaque groups. (h) Number of differentially expressed proteins in different types of plaque. (i) Venn diagram showing the overlaps between DEPs. Color scale in g and h represents z-score (red, high; blue, low). n = 3 per group.

In total, we excised four sample groups in triplicate: cored and diffuse plaques from 5-month-old mice and 10-month-old mice, denoted as YCP, YDP, OCP, and ODP, respectively (**Fig. 6b; Supplementary Movie 8-9)**. Each group contained approximately 30 individual plaques with 10 plaques collected per brain slice, all sampled from the cortex of the left hemisphere (**Supplementary Fig. 12**). To verify microsampling precision for each excised plaque, expanded tissue samples were imaged before and after microdissection (**Supplementary Fig. 12**).

Subsequent MS data analysis using the DIA-NN software identified 21,322 peptide precursors corresponding to 4,064 different proteins across all collected samples, with 2,913 proteins constituting a common core proteome (**Fig. 6c**), indicating the high completeness of the collected dataset. PCA revealed distinct clustering of cored plaques (YCP, OCP) and diffuse plaques (YDP, ODP), indicating noticeable proteomic differences between plaque types (**Fig. 6d**). Next, we focused on the DEPs in different plaque types during the moderate and late AD stages (**Fig. 6e,f**). We identified 248 DEPs in the YCP group compared to the YDP group, and 232 DEPs were detected in the ODP group compared to the YDP group. Among these, 45 proteins were consistently enriched in diffuse plaques across both age-group comparisons, including neuronal RNA-binding proteins (Elavl3, Stau2), synaptic vesicle components (Syt3), and autophagy regulators (Atg9a, Wdfy3). However, comparisons in 10-month-old mice revealed limited plaque-type differences (ODP vs. OCP: 43 DEPs) and no age-related changes within cored plaques (YCP vs. OCP: 0 DEPs; **Fig. 6f**). The dataset of ODP_vs_YDP and YCP_vs_YDP shared 57% of DEPs, corresponding to 184 proteins (**Fig. 6e**). Pearson correlation analysis revealed high correlations among cored plaques regardless of age (YCP-OCP: r = 0.94–0.96, consistent with 0 DEPs) but markedly lower correlations between cored and diffuse plaques (r = 0.71–0.76), while CV cumulative curves demonstrated that young diffuse plaques (YDC) exhibited the highest proteomic variation (**Supplementary Fig. 13**). These observations suggest that moderate-stage diffuse plaques represent a molecularly distinct niche compared to cored plaques, wherein aging minimally affects cored plaque proteomic composition but progressively stabilizes diffuse plaque proteomes, thereby reducing inter-plaque-type differences through proteomic convergence over time. A detailed quantitative analysis of 10 well-characterized AD biomarkers across all four experimental groups revealed striking differential expression patterns between cored and diffuse plaques, which were largely preserved across age groups (**Fig. 6g**). Specifically, App, Apoe, and Clu, proteins involved in Aβ metabolism and clearance, showed ∼3- 8.9-fold higher expression in cored plaques versus diffuse plaques across both age groups (**Supplementary Fig. 14**). In turn Gfap, a marker of reactive astrocytosis, and Apoa1, the major high-density lipoprotein component with anti-inflammatory properties, demonstrated 2.0- to 2.7-fold and 1.5- to 3.0-fold enrichment, respectively, in cored plaques (**Supplementary Fig. 14**). In contrast, tau-related proteins (Mapt and Gsk3b) and synaptic proteins (Syp, Syngap1, and Synj1) exhibited minimal plaque-type specificity, with no consistent pattern observed between plaque types. Extended proteomic analysis identified an additional 7 proteins, namely phospholipase D3 (PLD3), inositol polyphosphate-5-phosphatase D (SHIP1), platelet-activating factor acetylhydrolase (PAF-AH vC1q), tumor necrosis factor related protein 4 (CTRP4/C1QTNF4) Midkine (MDK), Palmitoyl-protein thioesterase 1 (PPT1), and nucleosome assembly protein 1-like 4 (NAP1L4), showing consistent core enrichment across both age groups, suggesting core structural proteins that define plaque architecture independent of temporal progression (**Fig. 6h**). Further investigation of the biological implications of the proteomic changes using GO biological process enrichment analysis of DEPs between YDP and YCP groups revealed significant enrichment in metabolic processes (including amyloid precursor protein metabolism) and transport pathways (peptide and neurotransmitter transport; **Supplementary Fig. 14**).

Further exploration with Kosmos provided additional biological findings that both corroborated and extended the above analysis (**Supplementary Data 3**). Consistent with our PCA results (**Figure 6d**), Kosmos confirmed that plaque type, rather than age, represents the dominant source of variance in the proteome. It also identified App and Apoe as key proteins that distinguish the plaque types, aligning with their differential expression observed in our analysis (**Fig. 6f**). Kosmos further proposed a model in which cored and diffuse plaques represent functionally distinct molecular states established early in the disease process. It suggested a progression from an early metabolic-stress state, as indicated by pathway enrichment at 5 months, to a distinct late-stage phenotype by 10 months, characterized by heightened oxidative stress, immune activation, and apoptosis. Kosmos linked this late cored plaque state, involving potential signaling changes such as an H-Ras-JNK switch, directly to the observed compact morphology and associated nuclear distortion, offering a molecular hypothesis for the observed localized cytotoxicity. This contrasts with diffuse plaque development, which is associated with the maturation of a keratin-based structural program and the exclusion of neuronal proteins, rather than with escalating toxicity. While these hypotheses are yet to be experimentally validated, they offer valuable directions for future investigations in the potential mechanisms underlying plaque heterogeneity and differential toxicity across age.

In summary, by integrating super-resolution 3D imaging with image-guided spatial proteomics, we resolved the molecular heterogeneity of distinct amyloid plaque subtypes in 5xFAD mice, revealing that morphologically distinct core and diffuse plaques represent functionally divergent pathological entities. Specifically, cored plaques exhibit significant enrichment of apolipoproteins and immune mediators that drive plaque consolidation and neuroinflammation, suggesting a two-stage pathogenic model in which initial diffuse Aβ deposition transitions to compact core formation via apolipoprotein-mediated aggregation and immune activation.

## Discussion

Technologies that enable simultaneous assessment of nanoscale structures and molecular profiles within intact tissue samples are essential for elucidating structure-function relationships in biological systems. The ExM framework has proven invaluable in this context, having been successfully applied to enable untargeted *in situ* genomics and transcriptomics in intact samples with nanoscale precision^13,14^. While protein retention ExM has been combined with intact tissue for untargeted bottom-up MS-based proteomics analysis, earlier methods, including proExM-MS^57^, ProteomEx^19^, and FAXP^22^, did not fully realize the potential for 3D super-resolution imaging prior to proteomic analysis. By integrating MS-compatible fluorescence staining of expanded tissue, microProteomEx uniquely combines 3D super-resolution imaging on conventional microscopes with subsequent deep proteomics at scalable spatial resolution from 28 to 100 µm, enabling comprehensive structural and molecular profiling of spatially defined regions of interest within the same sample.

In addition, microProteomEx offers several key advantages over similar spatial proteomics methods, particularly DVP^20^ and FAXP^22^. Compared to DVP, which employs confocal microscopy for sample imaging, microProteomEx enables 3D super-resolution fluorescence imaging prior to proteomic analysis, allowing direct visualization and analysis of subcellular structures at nanoscale resolution under conventional microscopes. Second, while DVP relies on laser capture microdissection (LCM) and specialized automated instrumentation for sample handling, microProteomEx can be performed using only off-the-shelf reagents and simple accessories such as biopsy punches and conventional diffraction-limited microscopes. Critically, microdissection in microProteomEx can be performed manually under image guidance; for example, we successfully microdissected individual kidney glomeruli (∼0.278 nL each) by hand using a biopsy punch. In this regard, microProteomEx represents a significantly more accessible method than DVP, requiring minimal specialized equipment while enabling nanoscale imaging with conventional microscopes. It should be noted, however, that these approaches are not mutually exclusive, as the DVP platform can, in principle, be integrated with expanded samples, enabling complementary analysis workflows that combine the strengths of both methods.

Compared to FAXP^22^, microProteomEx introduced several important technical and methodological innovations. First, we optimized the fixation and protein-anchoring protocol to improve compatibility with mechanically challenging tissues such as the heart and kidney (**Figure 7**). By performing fixation and protein anchoring simultaneously, we reduced the harshness of the subsequent homogenization step while preserving tissue integrity, which is particularly critical for mechanically tough tissues. This improved fixation method increased proteomic yield by nearly 3-fold compared with conventional fixation with 4% PFA and homogenization. Second, the integration of SyproRed staining provides a fast and simple way to visualize nanoscale morphological details (*e.g.*, sarcomeres in the heart, glomeruli in the kidney, and the core of beta-amyloid plaques in the 5xFAD mouse brain) compared to immunostaining used in FAXP. Third, microProteomEx extracts peptides from small microdissected regions without requiring custom-built accessories, such as the in-tip digestion device used in FAXP. Instead, we employ a simple re-embedding strategy in a polyacrylamide hydrogel, which can be easily prepared from the same monomers used for the initial expansion hydrogel. This approach further simplifies the technical requirements and improves accessibility. Fourth, we demonstrated microProteomEx across a wide range of sample types, including kidney, heart, liver, brain, and human skin, and combined it with 3D super-resolution fluorescence imaging. Altogether, these technical innovations make microProteomEx more versatile and accessible than FAXP while maintaining high proteomic depth and spatial resolution.

**Figure 7.**
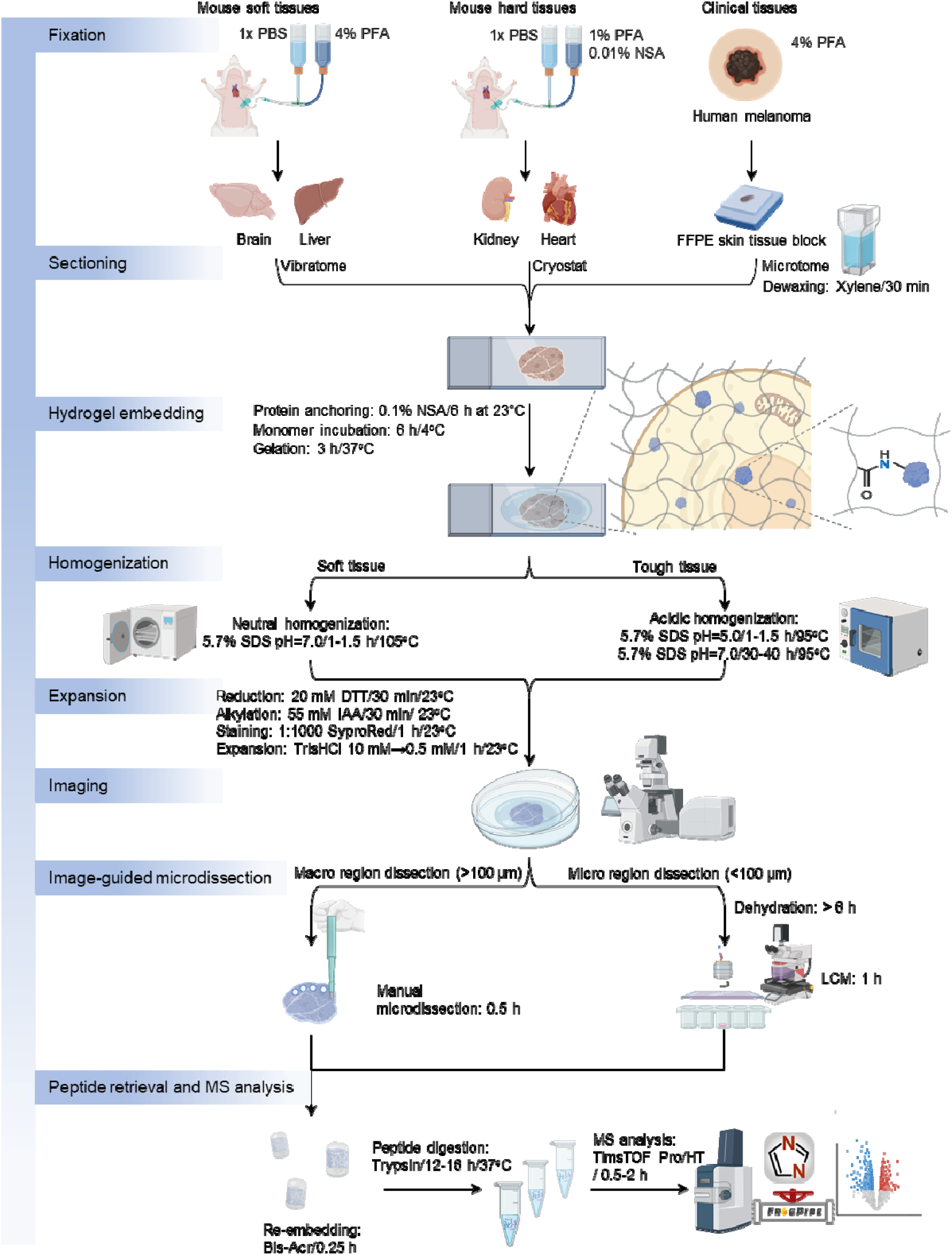
Schematic workflow of microProteomEx applied to three tissue categories, including soft tissues (brain, liver), mechanically tough tissues (kidney, heart), and archived clinical tissues (human skin FFPE blocks). The workflow consists of i) fixation (soft and hard tissues are perfused and fixed with paraformaldehyde; FFPE clinical samples are processed directly from formalin-fixed paraffin-embedded blocks); ii) sectioning (tissue sections are prepared according to tissue type; FFPE sections undergo dewaxing after sectioning); iii) hydrogel embedding (tissue sections are anchored to a polyacrylamide hydrogel matrix via in situ gelation to form a tissue-hydrogel composite); iv) homogenization (sample is treated with SDS-based buffer under tissue-type-specific conditions to ensure complete and uniform homogenization); v) expansion (homogenized sample undergo reduction, alkylation, and fluorescent staining, followed by isotropic expansion in aqueous buffer. vi) imaging (expanded samples are imaged by confocal fluorescence microscopy); vii) image-guided microdissection (ROIs are identified from fluorescence images and isolated by either manual microdissection with biopsy punches for >100 µm regions or LCM for <100 µm regions); viii) peptide retrieval and MS analysis (dissected hydrogel fragments are re-embedded, digested with trypsin, and the resulting peptides are analyzed by nano-LC-MS/MS, with data processed using FragPipe).

The application of microProteomEx to archival FFPE melanoma samples demonstrates its direct translational utility. By coupling super-resolution confocal imaging with spatially guided proteomics, we resolved distinct proteomic landscapes between GCMN and MM regions within the same tissue section. The proteomic transition from GCMN to MM identified in this study was characterized by a broad downregulation of ECM components, alongside upregulation of FASN and the biosynthetic machinery. These findings are consistent with several previous independent melanoma proteomics and transcriptomics studies. For example, Naimy et al. reported decreased levels of laminins, collagens, and extracellular molecules, as well as FASN upregulation, in melanoma relative to nevi across 77 FFPE melanoma samples^58^. The Xiang et al. study also identified a “cell proliferation subtype” characterized by elevated biosynthetic and translational activity, consistent with the mTORC1-driven anabolic program and upregulation of translation and tRNA aminoacylation pathways we observe in MM^59^. The transcriptomic meta-analysis by Zia et al. demonstrated that the transition from benign to early-stage melanoma involves processes that reflect active ECM remodeling rather than ECM retention^60^. Furthermore, ECM downregulation in MM is consistent with the broader cancer ECM literature^42–44^. Degradation of the organized ECM during malignant transformation supports invasion and proliferation^44^. In melanoma specifically, the loss of MFAP2 and EFEMP1 reflects the dismantling of the elastic fiber network that normally constrains cell migration^58^. The enrichment of H3K27me3 at the promoters of ELN and EHD2 in MM relative to GCMN, as revealed by our ChIP-seq data, implicates Polycomb-mediated epigenetic silencing as a mechanism driving ECM collapse. In this study, we employed an AI scientist, Kosmos^48^, to evaluate its capabilities for large dataset analysis and interpretation. Kosmos analysis of two orthogonal datasets (proteomics and ChIP-seq) independently converged on the same core biological findings as our conventional bioinformatic pipeline, identifying ECM/adhesion collapse and promoter-centric Polycomb reprogramming as major discoveries. This result provides initial validation of the AI agent’s analytical coherence, which can complement human-driven analysis of large spatial omics datasets to surface mechanistic relationships not immediately apparent from conventional pathway enrichment alone. Altogether, these results demonstrate the utility of microProteomEx for spatially resolved proteomics of large clinical FFPE archives that have historically been underexploited by deep proteomic methods. When coupled with AI-driven analysis tools such as Kosmos, microProteomEx enables systematic, hypothesis-generating interrogation of clinical tissue collections, with direct implications for biomarker discovery, diagnostic stratification, and the identification of metabolically regulated vulnerabilities amenable to therapeutic targeting.

Elucidating the molecular composition of amyloid plaques beyond Aβ alone represents a fundamental yet unresolved challenge in AD research. Prior investigations have been restricted to proteomic profiling of individual plaques, without differentiating between cored and diffuse plaque subtypes^61–63^. Using the microProteomEx workflow, we achieved precise structure-guided isolation of cored and diffuse plaques at two disease stages in the 5xFAD model, thus revealing morphology- and age-resolved proteomic landscapes within a unified experimental framework for the first time. The number of identified proteins (4,064 across all conditions) matches or exceeds that reported in prior plaque-focused studies^61–63^. The proteomic landscape of cored versus diffuse plaques supports a two-stage compositional model in which early diffuse deposits are enriched for cytoskeletal regulators (microtubule crosslinking factor 1, Mtcl1; tubulin folding cofactor C, Tbcc), vesicle trafficking components (sorting nexin 15, Snx15; Rab GTPase effector Rufy2), and neuronal RNA-binding proteins (Stau2, Elavl3), while cored plaques accumulate a distinct machinery of apolipoproteins (Apoe, Clu), reactive astrocyte markers (glial fibrillary acidic protein, Gfap), growth factors (pleiotrophin, Ptn; midkine, Mdk; and vitronectin, Vtn), proteases (HtrA serine peptidase 1), and autophagy-related proteins (Rag GTPase B, Rragb, and the E3 ubiquitin ligase Lrsam1).

Notably, 5 of our 13 consensus core-enriched proteins, namely Apoe, App, Clu, Ptn, and Htra1, have been independently reported in mouse and human plaque proteomics datasets^62,63^, providing cross-study validation of microProteomEx fidelity. Among consensus core-enriched proteins, Apoe and Clu showed the strongest co-enrichment in both age groups, suggesting a coordinated apolipoprotein response at the fibrillar core. For example, Jackson et al. showed that Clu accumulates specifically in the synaptic compartment in AD, with the highest levels in APOE4 carriers, and that synapses containing both clusterin and oligomeric Aβ are most abundant near plaques in APOE4 cases^64^. Based on these findings, Jackson et al. suggested that Clu and ApoE may work together to drive synaptic damage^64^. Gfap showed cored-plaque enrichment, consistent with the established preferential association of GFAP-positive reactive astrocytes with dense-core and neuritic plaques over diffuse deposits in human AD^65,66^. In turn, pleiotrophin (Ptn), a heparin-binding growth factor independently identified by Levites et al.^63^, showed the highest mean fold-change of any consensus protein. Furthermore, midkine (Mdk), the closest structural paralog of Ptn, was also detected as a consensus core-enriched protein in our dataset, showing the highest fold-enrichment in cored versus diffuse plaques among the additional core-enriched proteins identified by extended proteomic analysis (**Fig. 6h**). However, the exact role of the Mdk/Ptn family at the fibrillar interface remains mechanistically unresolved. For example, Zaman et al. demonstrated that recombinant Mdk inhibits Aβ40 and Aβ42 fibril assembly in vitro and that Mdk knockout in 5xFAD mice increases amyloid burden, supporting a protective function^67^. Conversely, Levites et al. showed that overexpression of either Mdk or Ptn in CRND8 mice accelerates amyloid deposition in plaques and cerebrovascular amyloid, and that both proteins bind fibrillar Aβ in vitro, suggesting they may also act as amyloid scaffolds that promote pathological accumulation□³. Thus, the net effect of Ptn enrichment within cored plaques remains to be further investigated. Htra1, a secreted serine protease enriched in human EOAD and DS plaques^62^, was directionally consistent across both comparisons, potentially reflecting a failed proteolytic clearance response within compact deposits.

Among proteins not previously reported in plaque proteomics, such as Rag GTPase B (Rragb), which regulates mTORC1 signaling at the lysosomal surface and amino acid sensing, and Lrsam1, an E3 ubiquitin ligase involved in ubiquitin-mediated protein degradation and autophagy, their co-enrichment may implicate lysosomal-autophagic dysfunction as a feature of cored plaque consolidation^68^. Synaptoporin (Synpr), a synaptic vesicle protein, may reflect dystrophic neurite accumulation within the fibrillar core. We should also note that canonical microglial markers, including Trem2, Tyrobp, Spp1, Aif1, and Lgals3bp, were not detected in our dataset, most likely because these proteins are concentrated in the peri-plaque microglial halo rather than within the plaque structure proper^69,70^. Their absence is thus a feature of microProteomEx’s spatial selectivity rather than a limitation of sensitivity. In diffuse plaques, the 45 proteins consistently enriched relative to cored plaques across both age groups included cytoskeletal regulators (Mtcl1, Tbcc), vesicle trafficking components (Snx15, Rufy2), neuronal RNA-binding proteins (Stau2, Elavl3), and autophagy regulators (Atg9a, Wdfy3), consistent with early synaptic and proteostatic involvement prior to fibrillar consolidation. Tau (Mapt) and Gsk3β showed practically no fold change across comparisons, indicating that tau pathology is not yet plaque-subtype-specific at these stages, consistent with chemical imaging evidence that pyroglutamation of Aβx-42 precedes Aβ1-40 deposition during the diffuse-to-cored transition^71^.

Beyond this core, we identify compositional differences suggesting that cored and diffuse plaques are molecularly distinct entities rather than sequential morphological variants of a common substrate. Cored plaques were enriched for proteins associated with amyloidogenic processing (App, Apoe), lipid metabolism, Aβ clearance, and lysosomal function, indicating engagement of a dynamic proteostatic and clearance-related machinery that may reflect both the structural maturity of fibrillar deposits and compensatory mechanisms responding to sustained Aβ burden. In contrast, diffuse plaques at early disease stages were enriched for synaptic organization and vesicle trafficking proteins, including annexins and neurofilament-associated components, suggesting that these early deposits may actively disrupt synaptic maintenance prior to fibrillar maturation. Pathway-level analysis reinforced this interpretation: cored plaques at later stages showed disproportionate enrichment in lysosomal dysfunction and Aβ cleavage pathways, consistent with a shift from early synaptic involvement to late-stage proteostatic failure. We acknowledge that limited sample sizes necessitate cautious interpretation and that conclusions are hypothesis-generating pending validation in larger cohorts and orthogonal systems. Nevertheless, multiple lines of internal consistency, including cross-subtype reproducibility, cross-stage proteomic trajectories, and concordance with prior published datasets, support the biological relevance of the observed differences, establishing a spatially resolved proteomic foundation for mechanistic investigation and subtype-specific therapeutic targeting.

Despite these advances, microProteomEx has limitations. The current spatial resolution of 25–100 µm, while a substantial improvement over conventional workflows, remains above single-cell scale (∼10 µm), potentially limiting dissection of proteomic heterogeneity in highly complex cellular niches. Proteome depth does not yet match that of single-cell proteomics platforms, in part reflecting the fundamental challenges of mass spectrometry sensitivity at sub-nanoliter input volumes, rather than limitations intrinsic to the workflow itself (see FAXP method). The multi-step workflow, though reduced to ∼27 hours, may still pose challenges for high-throughput applications, and performance in other tissue types and sample preparations remains to be evaluated. Looking forward, integration of automated microdissection with machine learning–guided image annotation holds promise for pushing resolution below 25 µm, and coupling with spatial transcriptomics approaches such as ExST would enable integrated multi-omic molecular mapping within the same expanded tissue section. Prospective validation across larger FFPE cohorts and additional disease models will be essential to establish diagnostic utility and support integration into precision medicine workflows.

The integration of AI-driven analysis tools, such as Kosmos, represents an important yet underexplored dimension of spatially resolved proteomics workflows. As demonstrated here, Kosmos independently corroborated key findings from our standard analysis pipeline — including plaque-type proteomic dominance and the centrality of App and Apoe — while generating novel mechanistic hypotheses not immediately apparent from conventional pathway enrichment, such as a potential Hras–JNK signaling axis driving late-stage cored plaque phenotypes. While AI-generated hypotheses require experimental validation, their ability to synthesize across large datasets and surface non-obvious mechanistic relationships offers meaningful acceleration of the hypothesis-generation cycle, and we anticipate that pairing spatially resolved proteomic platforms with AI-assisted analysis agents will become an increasingly productive paradigm for navigating molecular complexity in both basic and translational research settings.

In summary, microProteomEx establishes a new benchmark for spatially resolved tissue proteomics by uniquely combining 3D super-resolution imaging with deep, morphology-guided proteomic profiling within a single, accessible experimental framework. By overcoming the core limitations of prior expansion proteomics methods, microProteomEx enables interrogation of structure-function relationships at spatial scales and tissue types previously inaccessible to untargeted proteomics. Its applicability across diverse biological systems and compatibility with multi-omic and AI-assisted analysis frameworks position microProteomEx as a versatile and broadly deployable platform, with far-reaching implications for disease biology, biomarker discovery, and the advancement of precision medicine.

## Methods

### Chemicals, reagents, antibodies, and buffer compositions

The complete list of solutions and buffers used for tissue processing and staining, including their purposes, specific names, compositions, and storage conditions, is provided in **Supplementary Table 1**. Suppliers and lot numbers for the chemicals and reagents used to prepare solutions and buffers are provided in **Supplementary Table 2**. The antibody concentrations used for immunofluorescence (IF) staining of brain slices and expanded gels are described in the immunostaining method sections below.

### Animal care and procedures

All mice were housed in strict barrier facilities with macroenvironmental temperatures and humidity ranging from 20-26□ and 40-70%, respectively. Food and water were provided ad libitum. Mouse rooms had a 12 h light/12 h dark cycle (7 AM to 7 PM). The housing conditions were closely monitored and controlled. All animal maintenance and experimental procedures were conducted in accordance with Westlake University Animal Care guidelines, and all animal studies were approved by the Institutional Animal Care and Use Committee (IACUC) of Westlake University, Hangzhou, China, under animal protocol no. 19-044-KP-2.

A total of 20 C57BL/6J mice at the age of 2-7 months, regardless of sex, were provided by the Westlake University Animal facility. C57BL/6J Tg (APPSwFlLon, PSEN1*M146L*L286V) 6799Vas/Mmjax mice in the C57BL/6J genetic background (5xFAD mice) were obtained from The Jackson Laboratory (strain 008730, RRID:MMRRC_034848-JAX). One female 5xFAD mouse (5-month-old) and one female 5xFAD (10-month-old) were used for this study. 5xFAD mice were genotyped by PCR analysis of genomic DNA extracted from tail biopsies. Specific primers targeting human APP and PSEN1 transgenes were used to identify the presence of mutations, with an internal control included to ensure DNA quality. PCR products were separated on a 1% agarose gel and visualized using a gel imaging system (GenoSens S2, Clinx, China). The expected band sizes confirmed the presence of the transgenes, ensuring accurate genotyping for subsequent experiments.

### Mouse tissue preparation

To harvest desired organs, the mice were deeply anesthetized with 1% sodium pentobarbital. After transcardial perfusion with 50 ml of 1x phosphate-buffered saline (PBS) at 20-23°C to ensure complete blood washout, the mice were perfused with 40 ml of 4% paraformaldehyde (PFA) or 1% PFA or 1% PFA with 0.1 mg/ml NSA buffer (all buffers at 20-23°C) depending on the required experimental design (see Results section and Figure legends for exact conditions of perfusion for reported samples). Right after perfusion, the whole liver, kidney, heart, and brain organs were dissected and stored in 1x PBS at 4°C until sliced. Tissues perfused with 4% PFA were sectioned at 30 μm using a vibratome (Leica 1200S, Germany) unless otherwise stated. Tissues perfused with 1% PFA or 1% PFA with 0.1 mg/ml NSA buffer were immersed in 30% sucrose buffer until buffer equilibration for 12-24 h at 4°C. Right before slicing, the collected tissues were frozen at -80°C (60 min on average) and sectioned at 30 μm using a cryostat (Leica CM3050S, Germany) following standard protocol. For the fixation method comparison shown in Figure 3, tissues perfused with 4% PFA were sectioned the same way as 1% PFA fixed tissue described above. For the AD application, the mice were perfused with 4% PFA, following the above methods, and the mouse brains were harvested. A small hole was punctured in the right hemisphere of the mouse brain to distinguish between the left and right hemispheres using a fine needle. Mouse brain tissue was sectioned at 40 μm using a vibratome (Leica 1200S, Germany). The brain slices were stored in 1x PBS buffer at 4°C until used.

### Immunostaining

Slice samples were blocked with the blocking solution for 2 hours at room temperature (20–26°C). Following blocking, the sections were incubated with the primary antibody diluted in antibody staining solution at the recommended concentration for 14-16 h at 4°C. After primary antibody incubation, the sections were washed five times with 1x PBS for 5 minutes each. The sections were then incubated with secondary antibody, diluted in antibody staining solution at the recommended concentration, for 2 hours at room temperature (20–26°C) in the dark. The sections were washed three times with 1x PBS for 5 minutes each and then mounted using ProLong™ Diamond Antifade Mountant and cover-slipped.

### Human Sample Source and processing

Human samples in this study were collected from Shanghai Children’s Medical Center and Shanghai Skin Disease Hospital. Written informed consent was obtained from the patient’s legal guardian. This study was approved by the institutional ethics committee of Shanghai Children’s Medical Center (Approval No.: SCMCIRB—K2022164-1) and conducted in accordance with the Declaration of Helsinki Principles. Specimens from the GCMN patients were fixed with formalin and processed into paraffin blocks for further experimental analysis.

This study included a single pediatric participant (internal patient ID: GCMN1)^38^ diagnosed with giant congenital melanocytic nevus (GCMN). The patient was a 6-year-old male presenting with malignant melanoma transformation. According to the 7b rule classification system, the lesion was categorized as body type, with Krengel classification parameters of G2 S3 C0 R2 N2 H1. Molecular genetic analysis revealed a pathogenic mutation in the SMARCA2 gene. The patient had no documented history of other systemic diseases or prior therapeutic interventions at the time of study enrollment.

### H&E Staining

Paraffin-embedded tissue blocks were sectioned at approximately 4-5 μm thickness using a microtome. The resulting sections were flattened, mounted onto glass microscope slides, and baked at 60°C for 1-2 hours to ensure firm adhesion. Deparaffinization was achieved by sequential immersion in xylene I and xylene II (10-15 minutes each). Sections were then rehydrated through a graded ethanol series: 100% ethanol I (5 min), 100% ethanol II (5 min), 95% ethanol (5 min), 90% ethanol (5 min), 80% ethanol (5 min), and 70% ethanol (5 min). Nuclear staining was performed by immersing sections in hematoxylin for 5-10 minutes, followed by rinsing under running tap water to remove excess dye. Sections were subsequently differentiated in hydrochloric acid-alcohol solution for 3-5 seconds and counterstained in eosin for cytoplasmic visualization for 3-5 minutes. Following eosin staining, sections were dehydrated through a reverse graded ethanol series (70%, 80%, 90%, 95%, 100% I, and 100% II; 5 minutes each) and cleared in xylene I and xylene II (10-15 minutes each). Morphological analysis of tissue cellular structure was conducted by light microscopy observation.

### Human Slice Preprocessing for Hydrogel-tissue

Paraffin-embedded tissue blocks were sectioned at approximately 12 μm thickness using a microtome. The tissue sections were flattened, mounted onto a custom-designed gelation chamber, a conventional glass slide with a round or square well (with a diameter of 15 mm or 15 mm x 15 mm square) of 100-200 µm deep (custom-manufactured by Shilaide, China). Dewaxing was achieved by immersion in xylene at least two times (10-15 minutes each). Sections were then rehydrated through a graded ethanol series: 100% ethanol I (5 min), 100% ethanol II (5 min), 95% ethanol (5 min), 90% ethanol (5 min), 80% ethanol (5 min), and 70% ethanol (5 min).

### Hydrogel-tissue composite processing

Tissue sections prepared as described above were briefly washed three times with 100 mM MES buffer (pH = 6.0) and then incubated with protein anchoring solution (0.1 mg/mL NSA in 100 mM MES at pH = 6.0) at room temperature (20–26°C) for 6 h. Then, the samples were briefly rinsed three times with washing buffer (100 mM MOPS at pH = 7.0) and transferred to a custom-designed gelation chamber as described above. The excess wash buffer was removed, and the tissue slice was flattened with a gentle brush. Activated Monomer Solution (ATMS; Monomer Solution with APS and TEMED) was carefully added onto the slice to fill the well, then covered with a coverslip (100 µm, custom-manufactured by Shilaide, China) and incubated at 4°C for 6 h. The gelation chamber was transferred to a vacuum oven (DZF-6000, Shanghai Sunrise Instrument, China) for polymerization reaction at 37°C in an N_2_ atmosphere for 3 h. After polymerization, the tissue-hydrogel composite was carefully removed from the gelation chamber and submerged in the homogenization buffer 2 in a 10 cm Petri dish (150466, Thermo Scientific, USA). The Petri dish was incubated in a humidified environment at 95°C for 10-14 h (brain: 10 h; liver: 14 h) in an oven, or at 105°C for 1-1.5 h (brain: 1 h; liver: 1.5 h) in an autoclave. For kidney and heart samples, the tissue-hydrogel composite was submerged in the homogenization buffer 1 at first at 95°C for 3-5 h (kidney: 3 h; skin: 3.5 h heart: 4 h) using the oven. Then buffer 1 is replaced by homogenization buffer 2 at 95°C for 30-40 h (kidney: 30 h; skin: 36 h; heart: 40 h) using the oven. For immunostaining samples, the homogenization process was performed at 95°C for 40 min using homogenization buffer 2. After homogenization, homogenized samples were transferred to a new 10 cm dish and washed 3 times with 100 mM Tris (pH=8.0) for 10 min per wash at room temperature (20–26°C). Reduction and alkylation of proteins were performed on the whole tissue-hydrogel samples by first submerging them in the DTT solution at 20-22°C for 30 minutes. Following the reduction step, the DTT solution was replaced with the IAA solution for the alkylation. The dish containing the hydrogel composite and alkylation buffer was wrapped in aluminum foil and incubated at room temperature (20–26°C) for 30 minutes. After reduction and alkylation, the samples were washed three times with 100 mM Tris (pH 8.0) for 10 minutes per wash at room temperature (20–26°C).

### Hydrogel-tissue composite immunostaining

Immunostaining of the hydrogel-tissue composite was performed following sample homogenization, with minor modifications to the previously described protocol. The composite was washed three times with 1× PBS, 5 minutes per wash. Subsequently, it was blocked in a blocking solution for 2 hours at room temperature (20–26°C). After blocking, the composite was incubated with the primary antibody diluted in antibody staining solution at the recommended concentration for 14–16 hours at 4°C. Following primary antibody incubation, the sections were washed three times with 1× PBS, 5 minutes per wash, and then incubated with the secondary antibody, diluted in antibody staining solution at the recommended concentration, for 2 hours at room temperature in the dark. The sections were washed again three times with 1× PBS, 5 minutes per wash, before proceeding to dye staining. Triton was excluded from the blocking buffer and the antibody staining solution to prevent potential blockages in the liquid chromatography (LC) system.

### Tissue-hydrogel composite staining with dyes

For Coomassie brilliant blue staining (CBB), tissue-hydrogel samples after homogenization were immersed into Coomassie blue staining work solution and incubated for 1.5 h at 95°C using the oven. For fluorescent dye staining, samples were immersed in the fluorescent dye staining solutions listed in Supplementary Table 1.

### Expansion of homogenized tissue-hydrogel composites

After staining, samples were expanded using gradient buffers: 10 mM Tris, 5 mM Tris, 1 mM Tris, and 0.5 mM Tris (all buffers pH= 7.4) for 15 min each at room temperature (20–26°C). For immunofluorescence stained samples, tissue-hydrogel composites were expanded in 1x PBS for 20 min at room temperature (20–26°C).

### Bright-field and fluorescence imaging

Imaging of whole tissue slice samples before expansion (mounted on a glass slide) and after expansion (mounted on the 10 cm Petri dish) was performed using Zeiss Fluorescence Stereo Zoom Microscope (Axio Zoom.V16, Zeiss, Germany) in brightfield mode controlled by ZEN v3.1 (blue edition) software. Fluorescence imaging of gel samples mounted on a 50 mm confocal dish (P50G-1.5-30-F-Sleeve, scicell, China) or a 384-well multiwell dish (Tko-p383099-408, Mattek, USA), depending on the size of the actual sample. The imaging process was performed using a Zeiss LSM980 confocal microscope (10x NA0.45, 20x NA0.8, 40x NA1.2) controlled by ZEN v.3.5 (blue edition) or Yokogawa CSU-W1 SoRa imaging setup of Nikon Field Scanning Spinning-Disk Confocal System (Nikon Instruments Inc., Japan) with objectives (4x NA0.2, 10x NA0.45, 20x NA0.8, 40x NA1.15, 100x NA1.49) controlled by NIS Elements v.5.21.00.

### Expansion factor measurements

To determine the expansion factors for each expanded sample, we measured the distance between two landmarks selected within tissue sample both before and after expansion. This measurement was conducted using a vernier caliper for direct physical assessment or, alternatively, by imaging with a stereomicroscope for digital quantification. These measurements allowed for precise calculation of the expansion factor by comparing pre- and post-expansion distances, providing a reliable metric for sample isotropy across experimental conditions.

### Laser capture microdissection

An expanded and fully hydrated stained gel samples were placed on a PET membrane slide (MMI, Glattbrugg, Switzerland) overnight at 20-23°C to dry out. The next day, the sample was imaged under the MMI CellCut Plus system (MMI, Glattbrugg, Switzerland) under 4x, 10x, and 20x objectives to identify regions of interest and microdissected using specific settings (the laser under the 10x objective was optimized to have a Power of 60, laser focus of 42 μm, cut velocity of 10 μm/s). Dissected samples were collected on the adhesive lids of 200 µl transparent tubes (MMI, Glattbrugg, Switzerland). Re-embedding activated gel buffer (15 µl) was added onto the adhesive cap of LCM sample collection tubes to cover the sample. Then the tube was carefully closed and placed at 50°C for polymerization reaction for 15 min using the oven. Then, the re-embedding gel is removed from the cap into 1.5 ml Protein LoBind tubes (Eppendorf tubes, USA) for later peptide digestion reaction as described below.

For AD tissue microdissection, six mice brain slices were selected (young AD replicate 1-3, old AD replicate 1-3) for fluorescence imaging and microdissection with the Leica LMD7 Laser Microdissection System (Germany). The glass slide with the dried gel of expanded mouse brain slices prepared as described above was placed on the microscope with the tissue side down. Four empty 200 µl collection tubes (Axygen, USA) were loaded into the collector at once, and the collection tubes were checked for contaminants under the bright field. The overview of each sample was created under bright-field mode with a UVl 5x/0.12 magnification objective lens for ease of subsequent operations. It was then switched to a 10x/0.32 magnification objective to search for region of interest. The laser under the 10x objective was optimized to efficiently cut the gel. The following settings were used a power of 60, aperture of 12, speed of 13, specimen balance of 16, head current of 100%, pulse frequency of 1131, and offset of 50. The contour around aβ-amyloid plaque was drawn and cut at 10x magnification under the GFP channel to ensure that only the desired samples were included, and then the image was captured as a pre-LCM record. Once the laser cuts the contour, the selected protein sample will fall into the collection tube due to gravity. The camera is switched to the collector to confirm the collection of the sample, and a post-LCM image is captured as a record. Re-embedding activated gel buffer (30 µl) was carefully added to the cap of the collection tube to cover the sample. Then the tube was closed and placed at 50°C for polymerization reaction for 10 min. The re-embedding gel is removed from the cap into 1.5 ml Protein LoBind tubes (Eppendorf tubes, USA) for later peptides digestion reaction as described below.

### Manual microdissection and proteomic sample preparation

Microdissections were performed manually using commercially available biopsy punches with diameters of 1.0 mm, 1.5 mm, 2.0 mm, and 3.0 mm (Integra Miltex, USA) from target areas of the expanded samples. The excised gel particles were transferred into PCR tubes (Axygen, USA) using fine tweezers. The re-embedding activated gel solution was added to cover the excised samples. The excised gel size determined the added volume re-embedding activated gel solution, specifically 8 μl for 1.0-1.5 mm; 9 μl for 2.0 mm; 10 μl for 3.0 mm). PCR tubes containing excised samples immersed with re-embedding gel solution were incubated in the heat block of thermomixer (Miulab, China) at 50□ for 15 min for polymerization. To wash out salt, re-embedded gel pieces were washed two times with 200 μl of 100 mM ABB buffer in the original PCR tube agitated at 600 rpm in the thermomixer at 20□ for 5 min. For CBB destaining, 200 μl 50%/50% (v/v) methanol/100 mM ABB were used to wash gel pieces twice under shaking at 600 rpm in the thermomixer, at 60□ for 5 min. 200 μl acetonitrile (ACN) was added to the tube to dehydrate gel pieces twice for 5 minutes under shaking at 600 rpm in the thermomixer at 25□ (gel pieces reduced in size and turned white during this step, which was normal).

To fully saturate the dehydrated gel, the sample was moved to 1.5 ml Protein LoBind tubes (Eppendorf tubes, USA) and 15 µl of trypsin buffer was added to completely cover gel particles and incubated on the ice for 2 h. For the peptide digestion reaction, 10 µl of 100 mM ABB buffer was added to the tube and incubated at 37°C for 12 h. After trypsinization, the peptides were extracted from the gel in the following steps and obtained solution were combined: 1) collect the original supernatant; 2) add 100 µL of 25 mM ABB, shake for 10 min at 25°C and collect supernatant; 3) add 100 µl 10%/90% (v/v) ACN/100 mM ABB, shake for 10 min, collect supernatant; 4) add 100 µL 50%/50% (v/v) ACN/5% formic acid and shake for 10 min at 25°C and collect supernatant; 5) add 100 µL 70%/30% (v/v) ACN/5% formic acid and shake for 10 min at 25°C and collect supernatant; 6) add 100 µL ACN, shake for 10 min at 25°C and collect supernatant (during these step gel pieces underwent shrinking).

Peptide collection buffers were dried in a SpeedVac (MS-SP004, Christ, Germany). 10 µL of reconstitute buffer was added, and the peptide solution was desalted using C18 spin columns (Pierce™ C18 Spin Tips, Thermo Fisher Scientific, US) and dried in a SpeedVac. The purified peptides were stored at –80°C until further MS analysis.

### LC/MS analysis under DDA mode

The peptides were resuspended in 0.1% formic acid automatically injected using Bruker’s nanoElute UHPLC System onto the 25□ cm column equipped with emitter (Aurora series, CSI, 25□cm□×□75 µm ID, 1.6 □µm C18, IonOpticks). Bound peptides were then eluted over 120 min at a constant flow rate of 300 nl/min and the mobile phases water/0.1% FA and ACN/0.1% FA (A and B, respectively) were applied in the linear gradients starting from 2% B and increasing to 22% in 90□min, followed by an increase to 35% B in 100□min, 80% B in 110□min, 80% B in 110–120□min. For low concentration sample, 60 min gratitude is applied. Bound peptides were then eluted over 60 min at a constant flow rate of 300 nl/min and the mobile phases water/0.1% FA and ACN/0.1% FA (A and B, respectively) were applied in the linear gradients starting from 2% B and increasing to 22% in 45□min, followed by an increase to 35% B in 50□min, 80% B in 55□min, 80% B in 55–60□min.

Eluted peptides were sprayed into a TIMS quadrupole time-of-flight instrument (timsTOF Pro 2, Bruker Daltonics) using a nano-electrospray source (CaptiveSpray source, Bruker Daltonics). Samples were measured in dda-PASEF mode, and the dda-PASEF windows scheme ranged in dimension m/z from 300 to 1500 and in dimension 1/K0 from 0.75 to 1.3, with Ramp Time 166□ms.

### LC/MS analysis under DIA mode

For AD application samples, samples were measured with a timsTOF HT mass spectrometer under DIA mode in 30 min gradient. The peptides were resuspended in 0.1% formic acid and automatically injected using Bruker’s nanoElute UHPLC System onto the 25□cm column equipped with emitter (Aurora series, CSI, 25□cm□×□75□µm ID, 1.6□µm C18, IonOpticks). Bound peptides were then eluted over 30 min at a constant flow rate of 300 nl/min and the mobile phases water/0.1% FA and ACN/0.1% FA (A and B, respectively) were applied in the linear gradients starting from 4% B and increasing to 24% in 22.5□min, followed by an increase to 37% B in 25□min, 80% B in 27.5□min, 80% B in 27.5–30 min. Eluted peptides were sprayed into a TIMS quadrupole time-of-flight instrument (timsTOF HT, Bruker Daltonics) using a nano-electrospray source (CaptiveSpray source, Bruker Daltonics). Samples were measured in DIA-PASEF mode and the DIA-PASEF windows scheme ranged in dimension m/z from 300 to 1500 and in dimension 1/K0 0.6–1.4, with Ramp Time 50 ms and window width 25Da.

### Analysis of mass spectrometric data

The DDA data were analyzed using FragPipe (version 21.1) platform with the MSFragger (version 4.1) search engine against a target-decoy Uniprot mouse database. The workflow “SpecLib” was used. The “IM-MS” was selected as input LC/MS file. Precursor mass tolerance was set by -15-15 PPM and the fragment mass tolerance was set by 0.05 Da. The digestive enzyme was trypsin, with cutting after “KR” but not before “P”. The MS1 quant was selected, and the other settings were set to default.

All MS files based on DIA mode were searched together in DIA-NN (version 2.1) against the mouse brain library (including AD and WT) generated in PrroteomEx. False discovery rates of precursors and proteins were set at 1%. Algorithm using a mass and MS1 mass accuracy of 10.0, scan windows of 0, and activated isotopologues, heuristic protein inference, no shared spectra, in single-pass mode. Protein inference was set off, and library generation was set as ‘smart profiling’, and ‘Robust LC (high precision)’ as the quantification strategy. Other settings were used as default parameters.

After DIA-NN analysis, the output result report.pg_matrix.tsv was used for later analysis. The contamination items were deleted, and the protein without a corresponding gene ID was supplemented using Protein ID.

### Bioinformatics analysis

Bioinformatics data analysis was carried out using the R statistical computing environment version 4.4.1. For numerical comparisons, Welch’s unpaired t-test (two-sided) was executed for two groups, and one-way ANOVA was executed for three groups. The statistical visualizations were made using PRISM (version 10.4.1). For subsequent statistical analysis, protein or gene intensities were log_2_-transformed. Pearson’s correlation coefficient was calculated after the exclusion of missing values.

Skin samples and AD samples proteomic data were primarily filtered and imputed using Fragpipe Analyst. Differentially expressed genes between each group were identified using a threshold of adjusted P-value < 0.01 and a fold change > 2. P-values were calculated using a two-tailed Student’s t-test and adjusted via the Benjamini–Hochberg method. GO enrichment analysis was conducted using the clusterProfiler package in R. Differentially expressed proteins were input into the enrichGO function to identify enriched GO terms in the categories of biological process (BP). Gene annotation was performed by mapping gene symbols to Entrez IDs (mouse (5xFAD) data were annotated using org.Mm.eg.db, while human skin data were annotated using org.Hs.eg.db). Additional plot customizations, such as axis scaling and legend optimization, were applied to improve clarity. The fast gene set enrichment analysis (GSEA) was performed using clusterProfiler to identify Cellular Component Ontology terms ranging in size from 10 to 300. The Reactome pathway-based analysis was conducted using the ReactomePA (version 1.47.0) package to identify gene sets ranging in size from 5 to 300. Gene sets with an adjusted p-value of less than 0.05, as reported by GSEA, were considered statistically significant.

### ChIP-seq analysis of clinical samples

ChIP-seq was performed on fresh tissue specimens of giant congenital melanocytic nevus (GCMN), and GCMN-melanoma by Shanghai Genefund Biotech Co., Ltd (Shanghai, China). Briefly, tissues were snap-frozen in liquid nitrogen and stored at -80°C until used. Upon analysis, tissues were mechanically dissociated into single-cell suspensions. Cells were cross-linked with 1% formaldehyde for 10 min at 23°C, and the reaction was quenched with 0.125 M glycine. Cross-linked chromatin was sonicated using an ultrasonicator under ice-cold conditions to generate fragments ranging from 200 to 500 bp. The fragment size was verified by agarose gel electrophoresis after reverse cross-linking. Sonicated chromatin was immunoprecipitated overnight with the H3K27me3 antibody (DF6941, Affinity Biosciences, USA) conjugated to Protein A/G magnetic beads (16-663, Magna ChIPTM, Sigma-Aldrich). An input sample was reserved for normalization. After immunoprecipitation, the beads were washed, and the bound chromatin was eluted. The eluates and input samples were reverse cross-linked by incubation with Proteinase K at 65°C overnight. DNA was purified by phenol-chloroform-isoamyl alcohol (25:24:1) extraction and treated with RNase. Libraries were constructed through sequential steps of DNA end repair, adenylation of 3’ ends, adapter ligation, and PCR amplification. The final DNA libraries were size-selected for fragments of 100-300 bp (including adapters). Library quality was assessed, and qualified libraries were sequenced on the Illumina NovaSeq 6000 platform and raw sequencing file were preprocessed by Genefund Biotech Co., Ltd.

### ChIP-seq data processing and quality control

Raw sequencing reads were first assessed for quality using FastQC. Low-quality bases and adapter sequences were removed using standard trimming procedures by fastp. Cleaned reads were then aligned to the human reference genome (hg38) using Bowtie2 with default parameters. PCR duplicates were removed using Picard MarkDuplicates. Sorted and indexed BAM files were generated using SAMtools. To generate normalized genome-wide signal tracks, deepTools bamCoverage was used to convert BAM files into BigWig (bw) format. Global quality of the ChIP-seq data was processed using deepTools. Data quality and enrichment were first assessed using plotFingerprint to evaluate signal distribution across the genome. To examine sample reproducibility and overall similarity, read counts were summarized across the genome using multiBamSummary, followed by Pearson correlation analysis.

For representative genes, ChIP-seq signal tracks were visualized using IGV. BigWig files were generated from aligned reads and displayed for GCMN and MM samples. Two downregulated proteins identified by proteomics, ELN and EHD2, were selected as examples to illustrate locus-specific H3K27me3 enrichment at transcription start sites.

To assess H3K27me3 enrichment at transcription start sites of downregulated proteins identified in spatial proteomics data, log2(IP/Input) BigWig tracks were generated using deepTools bigwigCompare with identical bin sizes and normalization parameters. Transcription start site coordinates (±3 kb) of proteomics-downregulated genes were extracted and analyzed using computeMatrix in reference-point mode.

### Isotropic expansion analysis

To analyze tissue expansion isotropically, we used a B-spline-based MATLAB package to register post-ExM images to pre-ExM images via landmark selection. Images were first converted to grayscale and downsampled to a 2000x2000 pixel maximum to optimize computational time. The pre-ExM image served as the static reference, while the post-ExM image was the moving image for registration. For initial rigid registration, five landmarks were selected and paired manually. Following this, twenty landmarks were placed across the tissue sample for non-rigid registration. The script generated the deformation vector field, root mean square error (RMS) plot, and overlay of pre- and registered post-ExM images. The final RMS plot with standard deviations for each tissue type was compiled from saved workspaces across samples.

### Kosmos analysis

For the melanoma MS dataset, Kosmos^48^ was provided with the pre-processed proteomic dataset, research objective: “Compare the two stages of melanoma, giant congenital melanocytic nevi (GCMN) and melanoma, acquired from the same patient. Identify biomarkers that differentiate the two and propose mechanisms of action.” and data description: “This is a csv file of proteins identified in FFPE samples of melanocyte GCMN (G) and melanoma (MM) tissues from the same patient, with 11 samples per group”. After 5 iterations, Kosmos produced the final report containing 4 micropublications (Supplementary Dataset 1), which were evaluated by researchers, and relevant findings were selected for further interpretation and validation.

For the ChIP-seq dataset, Kosmos was provided with the pre-processed proteomic dataset, research objective: “Compare the two stages of melanoma, giant congenital melanocytic nevi (GCMN) and melanoma (MM), acquired from the same patient. Identify biomarkers that differentiate the two and propose mechanisms of action.” and data description: “This is ChIP-seq broadpeak data for H3K27me3 in melanoma (MM) and giant congenital melanocytic nevi (GCMN) samples from the same patient.”. After 5 iterations, Kosmos produced the final report containing 4 micropublications (Supplementary Dataset 2), which were evaluated by researchers, and relevant findings were selected.

For the AD dataset, Kosmos was provided with the pre-processed proteomic dataset, research objective: “Two morphologically distinct types of plaque depositions were observed in the cortex region of 5xFAD mouse brain: C-plaques had bright and well-defined central parts stained with SyproRed, and had irregularly shaped DAPI stains; while D-plagues had diffused and dim SyproRed fluorescence in the center with unaffected nuclear shape. Investigate the proteome differences between C-plaques and D-plaques at two stages of AD pathology progression: moderate AD pathology (5-month-old female 5xFAD) and late-stage disease (10-month-old female 5xFAD). Propose a mechanistic hypothesis for the morphological differences of the plaques.” and data description: “Proteomic data of brain slices from cortex of left hemisphere of 5xFAD mouse: C-plaques from 10-month-old mice (OCP), D-plaques from 10-month-old mice (ODP), C-plaques from 5-month-old mice (YCP), and D-plaques from 5-month-old mice (YDP)”. After 10 iterations, Kosmos produced the final report containing 4 micropublications (Supplementary Dataset 3), which were evaluated by researchers, and relevant findings were selected.

All data analysis trajectories performed by Kosmos, including Jupyter notebooks and data, as well as literature search trajectories, can be accessed via hyperlinks in the PDF reports.

### Data analysis

The images were analyzed using ImageJ (version 1.53), ZEN software (v3.1; Zeiss), and NIS Elements Advance Research software (v.5.21.00; Nikon). MS data were analyzed using the FragPipe platform (version 21.1) with MSFragger (version 4.1) and DIA-NN (version 1.8.1). Statistical charts were generated using PRISM (version 10.4.1), R (version 4.4.1). Venn diagrams were generated using the Bioladder platform^72^. Isotropic analysis calculation performed with MATLAB (version R2024a).

### Data availability

The mass spectrometry proteomics data generated in this study have been deposited to the ProteomeXchange Consortium via the PRIDE partner repository with the dataset identifier PXD067963. The ChIP-seq data generated in this study are available from FigShare (DOI: 10.6084/m9.figshare.31991328).

## Acknowledgement

We would like to thank Hu Jin and Mingzhu Fan from the Mass Spectrometry & Metabolomics Core Facility at Westlake University for their assistance and discussions on sample preparation and LC-MS/MS analysis. We would like to thank Zhen Dong and Pingping Hu from Westlake University for assistance with processing MS samples. We thank Zhien Rong from Westlake University for help with image analysis. We would like to thank Wangqing Ge for helping with data organization and genotyping validation. The language of the first draft has been polished with ChatGPT and the Grok platform, and then proofread by the co-authors with Grammarly. This study is supported by the National Natural Science Foundation of China (Grant No. W2432024 and 32571589 to K.D.P.), the National Natural Science Foundation of China (Major Research Plan, Grant No. 92259201 to T.G.), the Key Research and Development Program of Zhejiang Province (Grant No. 2022C03037 to T.G.), the National Natural Science Foundation of China (grant number 82473508, 82173396 to Y.L), in part by the Research Center for Industries of the Future of Westlake University (K.D.P.), the São Paulo Research Foundation (FAPESP) (Grant No. 2025/00104-8 to G.A.C), and the Westlake Education Foundation. This work was supported by Michael Buckley, founder of the Violet Research Institute.

## Contribution

S.Z. and K.D.P. initiated the project, made high-level designs and plans, and interpreted the data. S.Z. developed re-embedding workflow with help from C.S. S.Z. developed the optimized fixation workflow with help from T.L. A.Y. performed Kosmos analysis. L.M., with help from A.D.W. and S.G.R., led the design and engineering of Kosmos. S.Z., X.C., and T.L. prepared fixed tissue samples. S.Z., W.T., P.P., H.Z, and Y.L. performed clinical sample preparation and analysis. S.Z., H.T., Y.W., and G.D. performed the AD sample experiment. S.Z. conducted proteomic data analysis and data visualization. G.A.C. performed dye comparison analysis. S.Z. and K.D.P. wrote the manuscript. K.D.P., Y.L., T.G., and H.Z. supervised the project.

