## Supplementary Materials for "Super-Resolution Visual ProteomEx for Hard Tissues and Clinical Samples"

**Supplementary Table 1.** Description of solutions and buffers used for sample processing.

| **Process** | **Specific name** | **Composition^a^** | **Storage condition** | **Purpose** |
| --- | --- | --- | --- | --- |
| Fixation | 4% PFA | 5 ml 16% PFA dissolve in 15 ml 1x PBS | -20℃ | Fix sample |
|  | 1% PFA | 1 ml 16% PFA dissolve in 15 ml 1x PBS | -20℃ | Fix sample |
|  | 1% PFA+ NSA | 1 ml 16% PFA and 150 μl anchor stock solution   dissolve in 15 ml 1x PBS | -20℃ | Fix sample and anchor proteins |
| Dewaxing | Dimethylbenzene | Dimethylbenzene | 22℃ | Dewax the paraffin |
|  | Sequential ethanol solution | 100%/95%/75%/50% ethanol diluted in ddH_2_O | 22℃ | Rehydrate samples |
| Protein anchoring | MES buffer | 5 ml 100 mM MES buffer (pH6.0) dissolved in 5 ml ddH_2_O | 4℃ | Decrease NSA hydrolysis |
|  | Anchor stock solution | 10 mg/ml NSA dissolved in 100 mM MES buffer | -20℃ | Stock solution |
|  | Anchor solution | 0.1 mg/ml NSA dissolved in 100 mM MES buffer | Freshly prepared | Anchor proteins via the primary amino groups |
|  | MOPS buffer | 1 ml 100 mM MOPS buffer (pH7.0) dissolve in 4 ml ddH_2_O | 4℃ | Stop anchoring |
| Monomer infusion and polymerization | PAE solution stock | 0.115g/ml PAE dissolved in THF | 4℃ | Crossing linker |
|  | Stock monomer solution | 30% (w/v) DMAA, 8.6% (w/v) SMA, 0.77% (v/v) PAE stock solution, and 0.35% (v/v) HCI in ddH_2_O | 4℃ | Gelation monomer solution |
|  | APS solution | 10% APS (w/v) dissolve in ddH_2_O | Freshly prepared | Polymerization initiator |
|  | TEMED solution | 10% TEMED (w/v) dissolve in ddH_2_O | Freshly prepared | Catalyst for the polymerization |
|  | Activated monomer solution | 90% Stock monomer solution /5% ddH_2_O/ 3% TEMED solution/ 2% APS solution | Freshly prepared | Sample gelation |
| Homogenization | Homogenization buffer 1 | 0.2M SDS, 40 ml Coomassie stock solution, 100 mM NaCl, 10%(v/v) acetic acid, add ddH2O to 50 ml | 22℃ | Restrain gel expansion during gel transfer and protein denaturation |
|  | Homogenization buffer 2 | 0.2M SDS, 40 ml Coomassie stock solution, 100mM NaCl, adjust pH to 7 with NaOH, add ddH2O to 50ml | 22℃ | Protein denaturation |
| Reduction | DTT solution | 20 mM DTT dissolved in 100 mM Tris-HCl (PH8.0) | Fresh prepared | Reduce disulfide bond |
| Alkylation | IAA solution | 55 mM IAA dissolved in 100 mM Tris-HCl (PH8.0) | Fresh prepared | Alkylate thiol groups |
| Staining | Coomassie stock solution | 8 ml Coomassie Blue Fast Stain Solution dissolved in 992 ml of ddH2O | 22℃ | Stock solution |
|  | Coomassie staining solution | 100 mM Tris-HCl (pH8.0) dissolve in Coomassie stock solution | 22℃ | Neutralizing acidity of Coomassie stock solution |
|  | Sypro red staining solution | Sypro red stock dissolved in PBS in 1:1000 ratio | Freshly prepared | Stain general proteins |
|  | DAPI staining solution | DAPI stock dissolved in PBS in 1:1500-1:2000 ratio | Freshly prepared | Stain DNA |
|  | Cy5 staining solution | Cy5 stock dissolved in PBS in 1:1000 ratio | Freshly prepared | Stain general proteins |
| Re-embedding | Re-embedding activated gel buffer | 50 μl gel solution stock, 2 μl TEMED solution, 1 μl APS solution dissolve in 100 μl ddH2O | Freshly prepared | Re-embed the exercised gel particle |
|  | Gel solution stock | 29% Acrylamide/1% bisacrylamide (w/v) dissolve in ddH2O | 4℃ | Stock solution for long-term storage |
| Digestion | Trypsin stock solution | 500 ng/μl trypsin (dissolved buffer: 50 mM acetic acid in ddH2O) | -80℃ | Stock solution for long-term storage |
|  | Trypsin work solution | 20% trypsin stock solution dissolved in 100 mM ABB solution | Freshly prepared | Sample trypsinization |
| Immunostaining | Blocking buffer | 5% BSA dissolved in 1x PBS (add 30 μl Triton/ 10ml for samples not processed for LC-MS/MS) | Freshly prepared | Improve specificity of immunostaining |
|  | Antibody staining buffer | 1% BSA dissolved in 1x PBS (add 30 μl Triton/ 10ml for samples not processed for LC-MS/MS) | Freshly prepared | Dilution of antibody stock solution |

^a^See Supplementary Table 2 for the reagent full names, suppliers, and lot numbers.

**Supplementary Table 2.** List of chemicals and reagents used for solution and buffer preparation.

| **Reagent name** | **Abbreviation** | **Supplier (Country)** | **Lot number** |
| --- | --- | --- | --- |
| 32% paraformaldehyde | 32% PFA | Electron Microscopy Sciences (USA) | 15710 |
| 1X PBS (pH7.4) | 1x PBS | Servicebio (China) | G4202-500ml |
| N-succinimidyl acrylate | NSA | TCI (Japan) | S0814 |
| Dimethyl sulfoxide | DMSO | Macklin (China) | D806645-500ml |
| 3-(N-morpholino)propanesulfonic acid, 0.5M, pH 7.0 | 0.5M MOPS | Macklin (China) | M885700-100ml |
| 2-Morpholinoethanesulphonic acid, 0.2M, pH 6.0 | 0.2M MES | Macklin (China) | M885671-250ml |
| Pentaerythritol allyl ether | PAE | Sigma-Aldrich (USA) | 251720-100G |
| Tetrahydrofuran | THF | Macklin | T818769-500ml |
| N, N-dimethylacrylamide | DMAA | Sigma-Aldrich (USA) | 274135-500ML |
| Sodium methacrylate | SMA | Sigma-Aldrich (USA) | 408212-250G |
| N, N,N′, N′-  Tetramethylethylenediamine | TEMED | Sigma-Aldrich (USA) | T7024-100ML |
| Ammonium persulfate | APS | Sigma-Aldrich (USA) | A3678 |
| 10% hydrochloric acid | HCl | Made by the Westlake University Uni platform |  |
| Sodium dodecyl sulfate | SDS | Macklin (China) | S817790-500g |
| DL-Dithiothreitol (powder) | DTT | Sigma-Aldrich (USA) | D9163-25G |
| Iodoacetamide (powder) | IAA | Sigma-Aldrich (USA) | I6125-100G |
| Commassie Blue Fast Stain Solution | CBB | Yeasen (China) | C0318051 |
| 4′,6-diamidino-2-phenylindole | DAPI | Abcam (UK) | ab228549-2mL |
| SYPRO™ Red Protein Gel Stain | Sypro red | Thermo Fisher scientific (USA) | S6654 |
| Trypsin (powder) | Trypsin | Promega (USA) | V5111 |
| Ammonium bicarbonate (powder) | ABB | Sigma-Aldrich (USA) | A6141-500G |
| Cyanine5 NHS ester, Amine-reactive red emitting fluorescent dye (powder) | Cy5 NHS ester | Abcam (UK) | 1263093-76-0 |
| ProLong™ Diamond Antifade Mountant |  | Thermo Fisher scientific (USA) | P36970 |
| Acetonitrile | ACN | Thermo Fisher scientific (USA) | 219087 |
| Bovine serum albumin | BSA | Sigma-Aldrich (USA) | B2064-50G |
| Triton™ X-100 | Triton | Sigma-Aldrich (USA) | X100-500ML |
| Acrylamide | AA | Sigma-Aldrich (USA) | A8887-500G |
| N,N′-Methylenebisacrylamide | Bis | Sigma-Aldrich (USA) | M7279-100G |

**Supplementary Figure 1.** Hydrogel re-embedding enables efficient peptide retrieval from LCM-dissected microsamples.

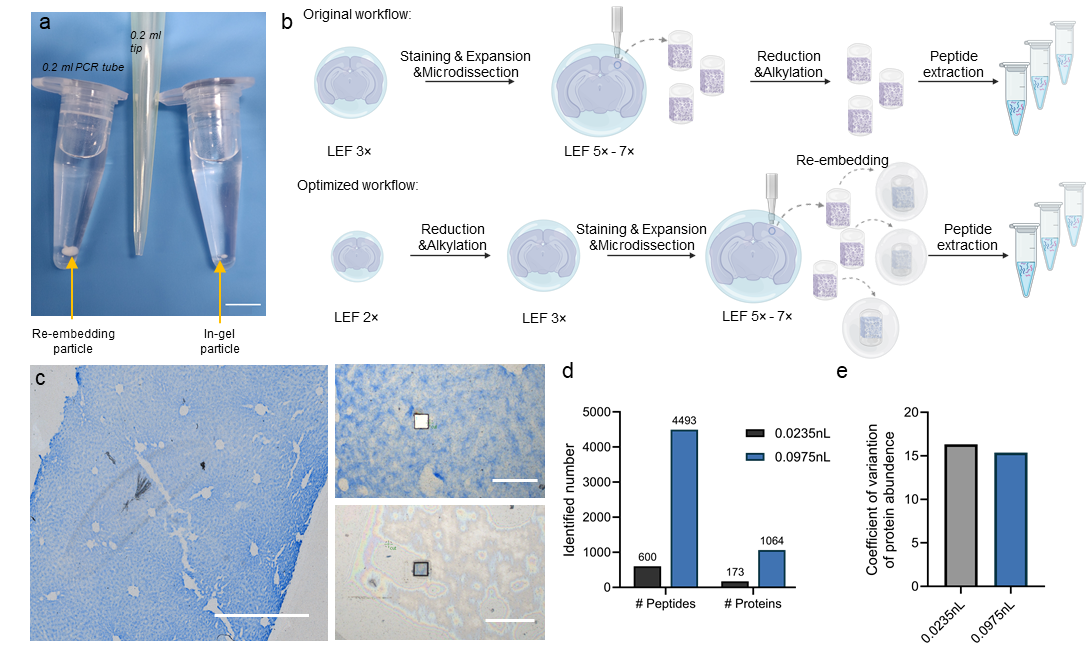

(a) Size comparison of re-embedding tissue-hydrogel particles, 0.2 ml Eppendorf tube, and re-embedded in-gel particle. Scale bar, 5 mm. (b) Schematic representation of the original ProteomEx workflow (up) and optimized workflow (down), including whole gel reduction and alkylation steps and microdissected particles re-embedding (created with BioRender). (c) Representative bright-field images of expanded mouse liver tissue stained with Coomassie brilliant blue (left) gross anatomy of mouse liver expanded samples; (right, up) magnified view of the cut region; (right, bottom) validation image of cut sample on the cap of collection tube; LEF = 3.5; scale bar = 300 µm. (d) Identified peptides and proteins number from LCM cut samples of mouse liver tissue processed using optimized workflow (tissue volume 0.0235 nL and 0.0975 nL, data was analyzed by DDA). (e) Coefficients of variation the identified peptides from LCM cut samples shown in panel d.

**Supplementary Figure 2.** Hydrogel re-embedding provides high reproducibility and stability comparison of protein identifications and quantification for microProteomEx.

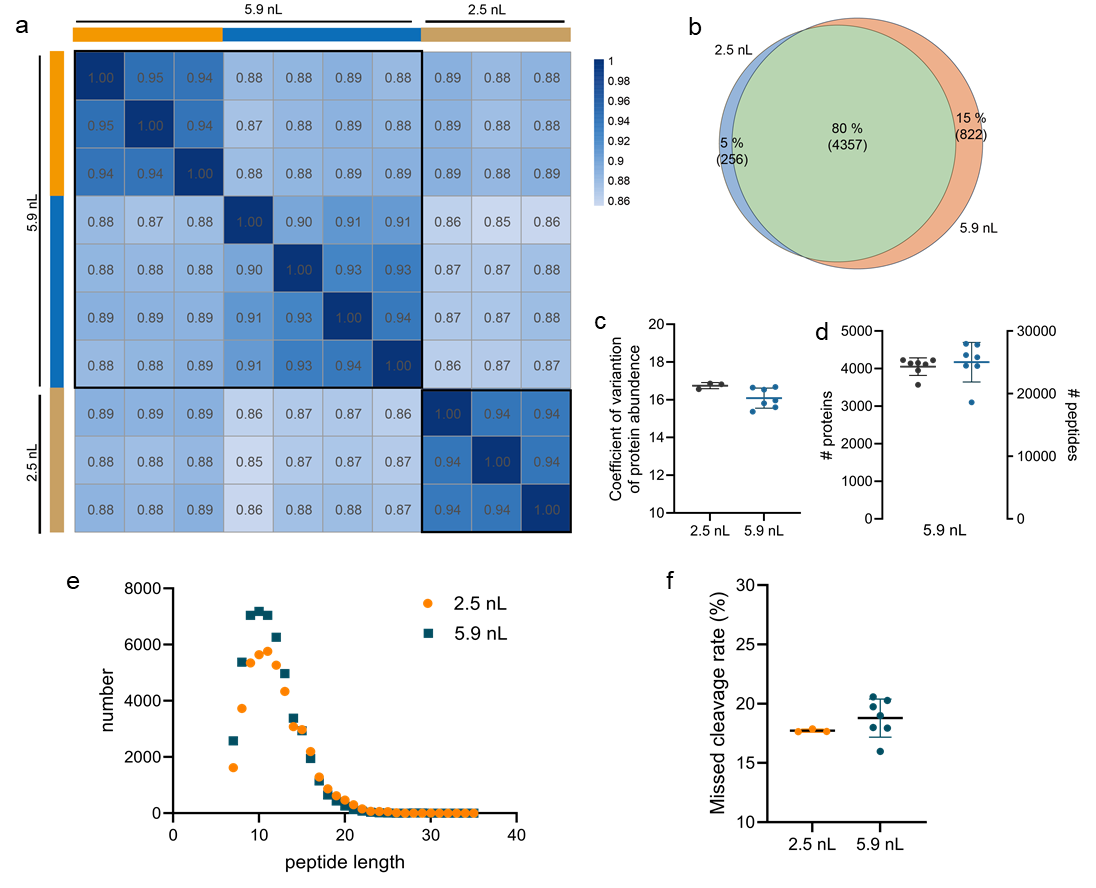

(a) Heatmap of Pearson correlations for protein quantification for each pair of samples from the ten samples shown in Figure 1a,b,c analyzed using the FragPipe software (n=3, 4, 3 punches from one liver tissue slice for each set from one mouse for microProteomEx (5.9 nL. 5.9 nL and 2.5 nL sample), respectively; the MS raw files corresponding to Figure 1a,b,c were used for analysis). The color bars indicate samples from separate tissue slices. (b) Venn diagram of the total protein identification for microProteomEx (2.5 nL and 5.9 nL samples). (c) Coefficient of variation of quantified protein abundance from the two different size volume samples shown in a. (d) Protein and peptide identification numbers in microProteomEx (data from Fig. 1a, c; tissue volume: 5.9 nL in microProteomEx). (e) Peptide length distribution from samples shown in a. (f) Missed cleavage rates of samples shown in a. Data were analyzed under DDA mode with Bruker timsTOF Pro2. Dot, individual data point, bar, mean, whiskers, standard deviation (SD).

**Supplementary Figure 3.** Comparison of Cy5 NHS ester and SyproRed staining for proteomic analysis of mouse liver tissue using microProteomEx.

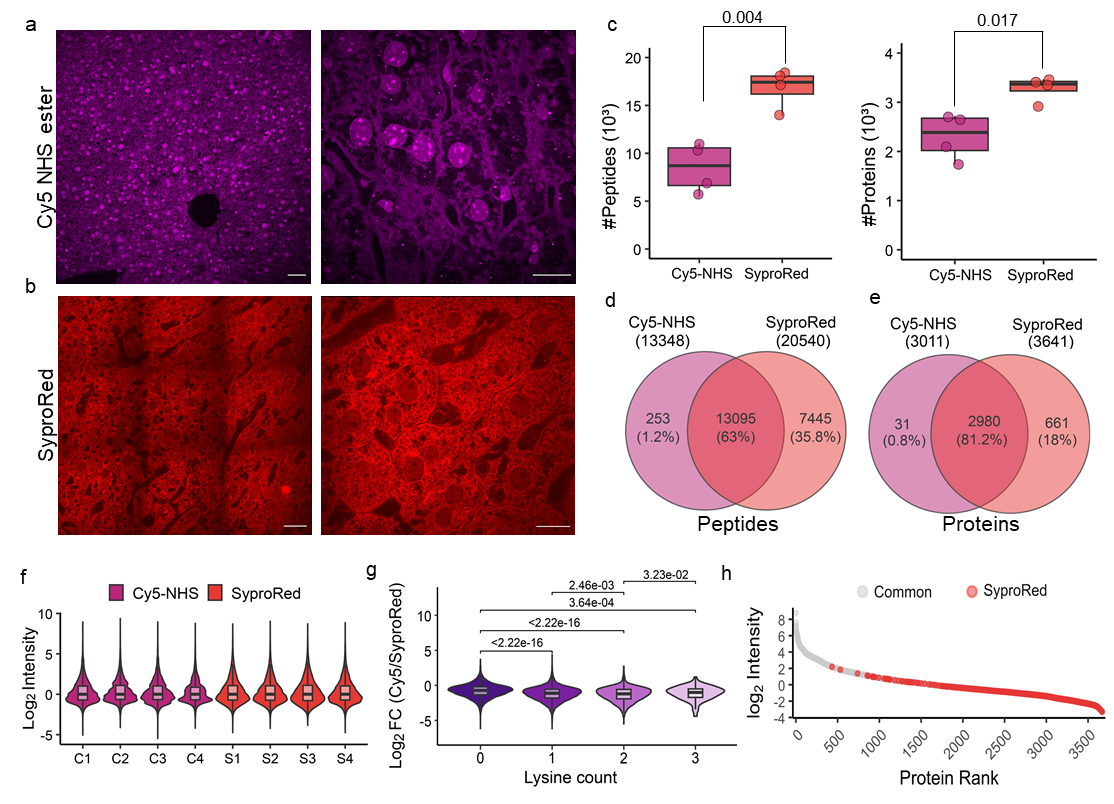

(a) Representative single plane confocal images (Cy5 NHS ester stained) of post-expansion mouse liver tissue (n=3 slices from one mouse). LEF = 3.62. Scale bars, 27.6 μm (left; physical size post-expansion, 100 μm), 13.8 μm (right; physical size post-expansion, 50 μm). (b) Representative single plane confocal images (SyproRed stained) of post-expansion mouse liver tissue (n=3 slices from one mouse). LEF = 4.02. Scale bars, 24.87 μm (left; physical size post-expansion, 100 μm), 12.43 μm (right; physical size post-expansion, 50 μm). (c) Number of peptide and protein identifications in Cy5 NHS ester-stained and SyproRed-stained mouse liver tissue shown in a-b (n=4 samples from 3 slices from one mouse each). (d, e) Venn diagram of the total peptide and protein identifications of Cy5 NHS and SyproRed groups. (f) Protein identification of each of the 4 samples in the Cy5 NHS (C samples) and SyproRed (S samples) groups. (g) Fold change of Cy5 NHS group compared to SyproRed group of different Lysine counts peptides. (h) Rank intensity of common and SyproRed unique proteins of Cy5 NHS and SyproRed groups.

**Supplementary Figure 4.** Image-guided manual and LCM microdissection of expanded mouse liver samples stained and visualized using SyproRed.

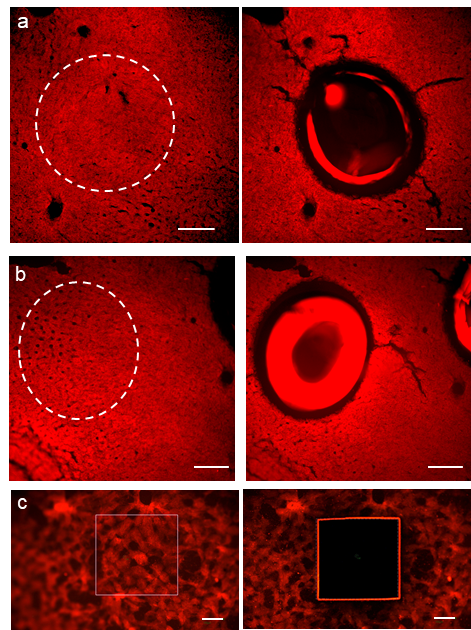

(a,b) Wide-field fluorescence images of mouse liver tissue before (left) and after (right) manual microdissection with biopsy punch (white circles indicate ROIs selected for microdissection). Scale bar, 1000 μm. (c) Wide-field fluorescence images of mouse liver tissue before (left) and after (right) microdissection with LCM. Scale bar, 100 μm.

**Supplementary Figure 5.** Evaluation of fixation methods and proteomic reproducibility across mouse tissues.

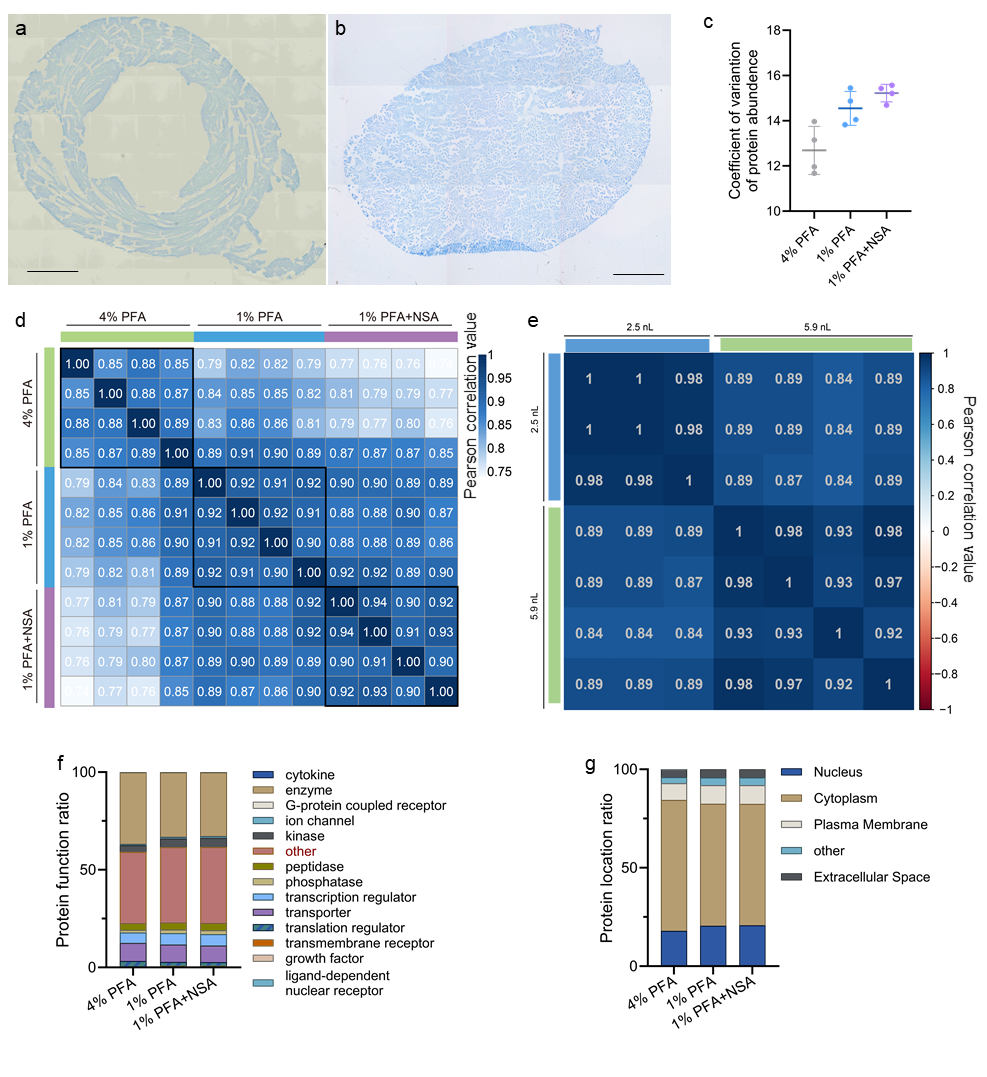

(a) Mouse heart slices stained with CBB. Scale bar, 904.2 μm (physical size post-expansion, 5000 μm). (b) Mouse kidney slices stained with CBB. Scale bar, 904.2 μm (physical size post-expansion, 5000 μm). (c) The coefficient of variation of quantified protein abundance for samples shown in Fig. 3a. Dot, individual data point; bar, mean; error bars, SD. (d) Heatmap of Pearson correlations for protein quantification shown in Fig. 3a. (e). Heatmap of Pearson correlations for protein quantification of 1%PFA+ NSA group. Four samples from Fig. 3a. All samples were analyzed using the FragPipe software (n=3, 4 punches from one, one liver tissue slice from two mice for optimized fixation protocol (2.5 nL and 5.9 nL sample), respectively). The color bars indicate samples from separate tissue slices. (f) The subcellular locations of (a) the identified protein ratio by the different fixation methods for the samples shown in Fig. 3a. Protein ID combination by groups. (g) The types of the identified protein counts ratio by the different fixation methods for the samples shown Fig. 3a. Protein ID combination by groups. (h-i) Representative images of re- and post- manual microdissection of mouse kiney and heart ROIs shown in Fig. 3. Scale bar: 100 μm.

**Supplementary Figure 6.** ROI excision quality and reproducibility of protein identifications in kidney and heart tissue processed with microProteomEx.

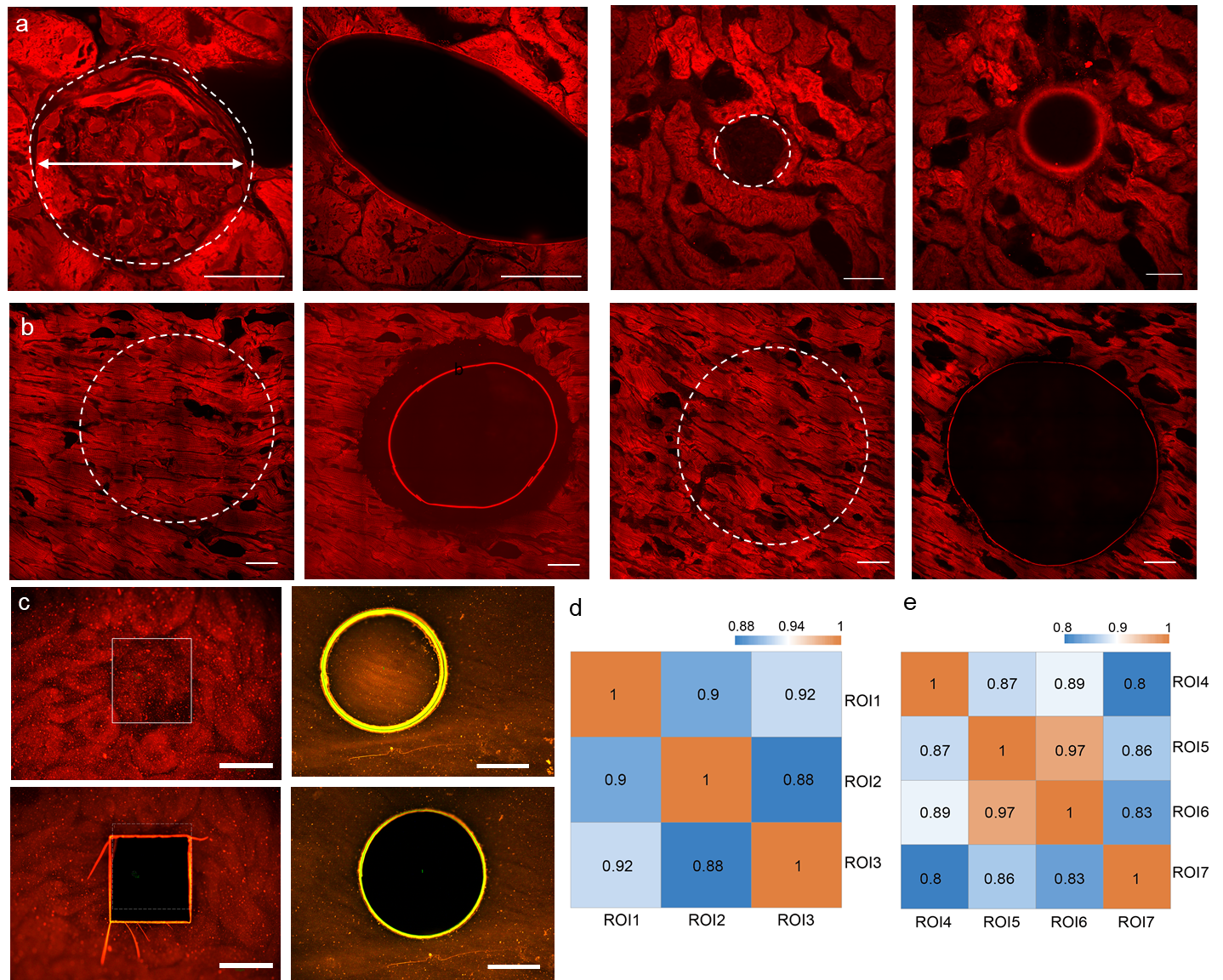

(a-b) Representative confocal single-plane fluorescence images of manual pre- and post-cut images of ROIs in mouse glomeruli and heart tissue stained with SyproRed post-expansion (n = 3 slices from 1 mouse for each tissue). (c) Representative brightfield fluorescence images of LCM pre- and post-cut images of ROIs in mouse (upper row) glomeruli and (lower row) heart tissue stained with SyproRed post-expansion (n = 3 slices from 1 mouse for each tissue). (d-e) Pearson correlation of protein identifications among ROIs of (d) single glomeruli and (e) heart samples shown in Fig.3. LEF: 5.79 (kidney tissue); 5.36 (heart tissue). Scale bars (physical size post expansion), 200 μm in a-c.

**Supplementary Figure 7.** Proteomic comparison between malignant melanoma (MM) and giant congenital melanocytic nevus (GCMN).

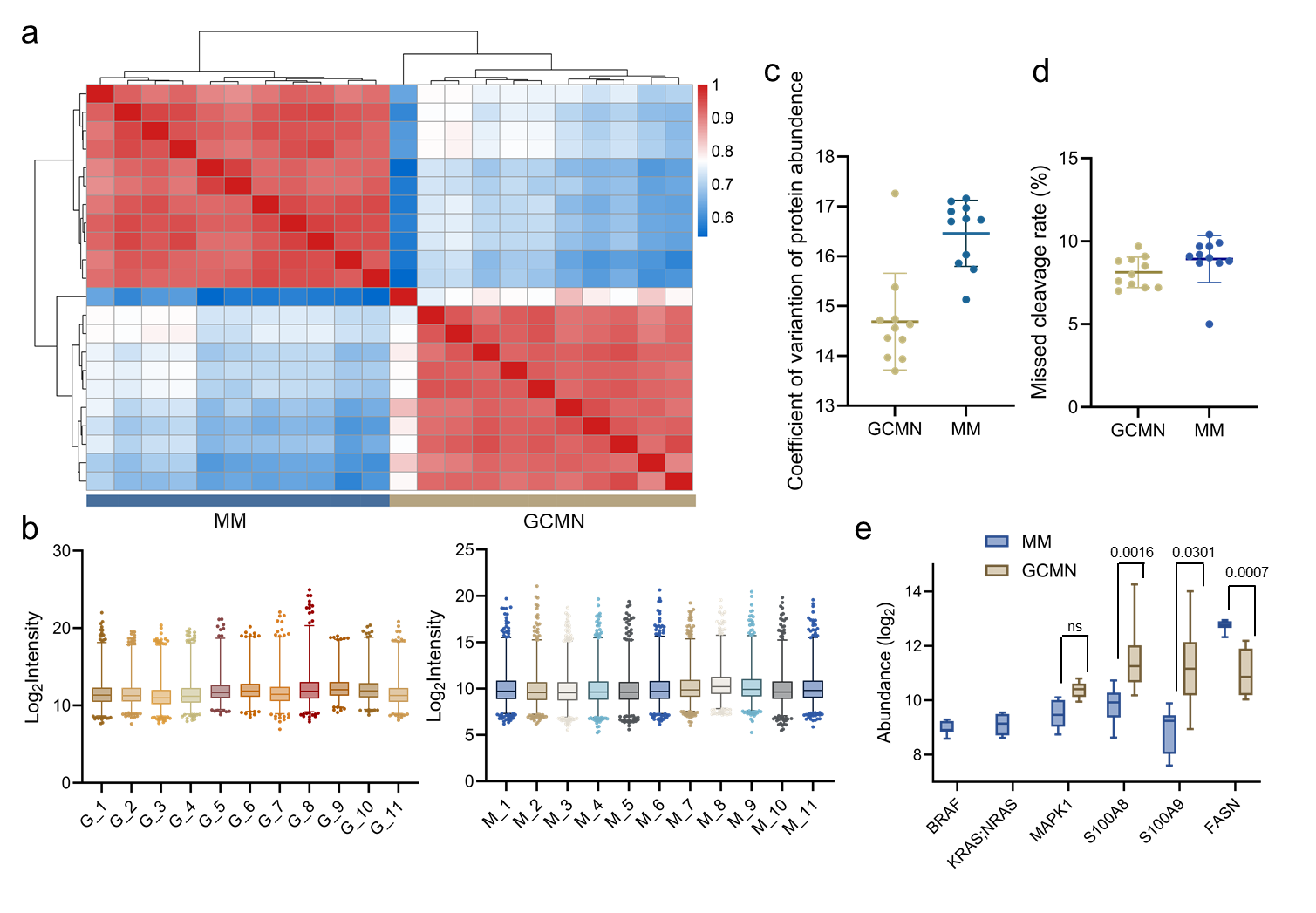

(a) Hierarchical clustering heatmap of Pearson correlation coefficients across MM and GCMN samples, shown in Fig. 4. (b) Quantitative intensity of protein identifications of MM and GCMN groups. (c) The coefficient of variation of quantified protein abundance for the samples shown in b. Dot, individual data point; bar, mean; error bars, SD. (d) Missed cleavage rate of samples shown in b. (e) Comparative distribution of protein abundance across the interested proteins in MM and GCMN samples.

**Supplementary Figure 8.** Epigenetic features associated with proteomics-downregulated genes in clinical skin samples.

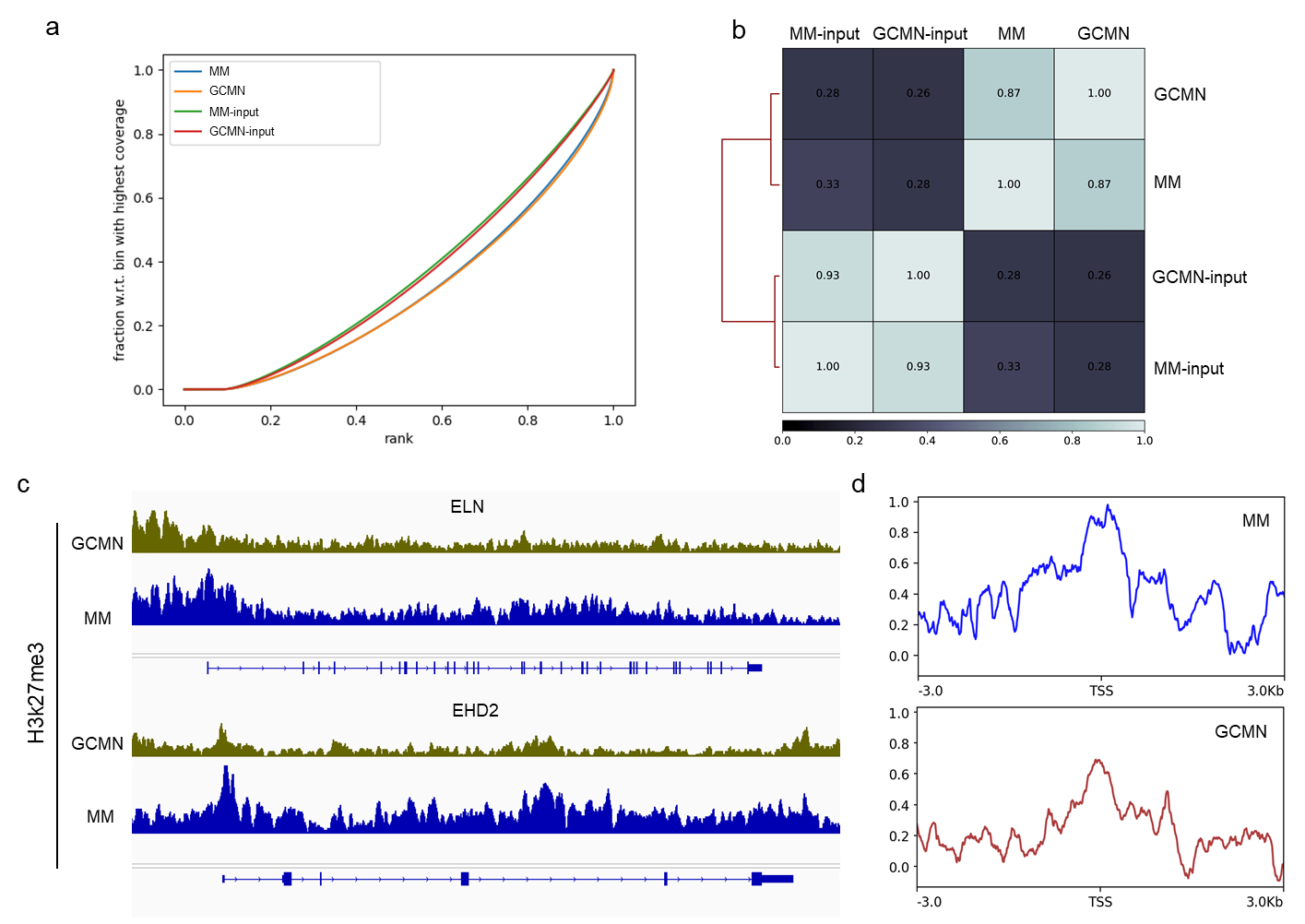

To investigate whether epigenetic repression contributes to the downregulation of proteins observed in GCMN and MM samples, we analyzed H3K27me3 ChIP-seq profiles in parallel with proteomics data. (a) Quality assessment showed the expected enrichment pattern of H3K27me3 ChIP-seq signal relative to input libraries. (b) Pearson correlation analysis revealed high reproducibility among biological replicates and clear clustering of samples by condition. (c) Genome browser visualization of two downregulated proteins, ELN and EHD2. (d) Metagene analysis centered on TSSs (±3 kb) of proteomics-downregulated genes in GCMN and MM samples.

**Supplementary Figure 9.** Validation of immunostaining of the mouse brain tissue post-expansion with microProteomEx protocol.

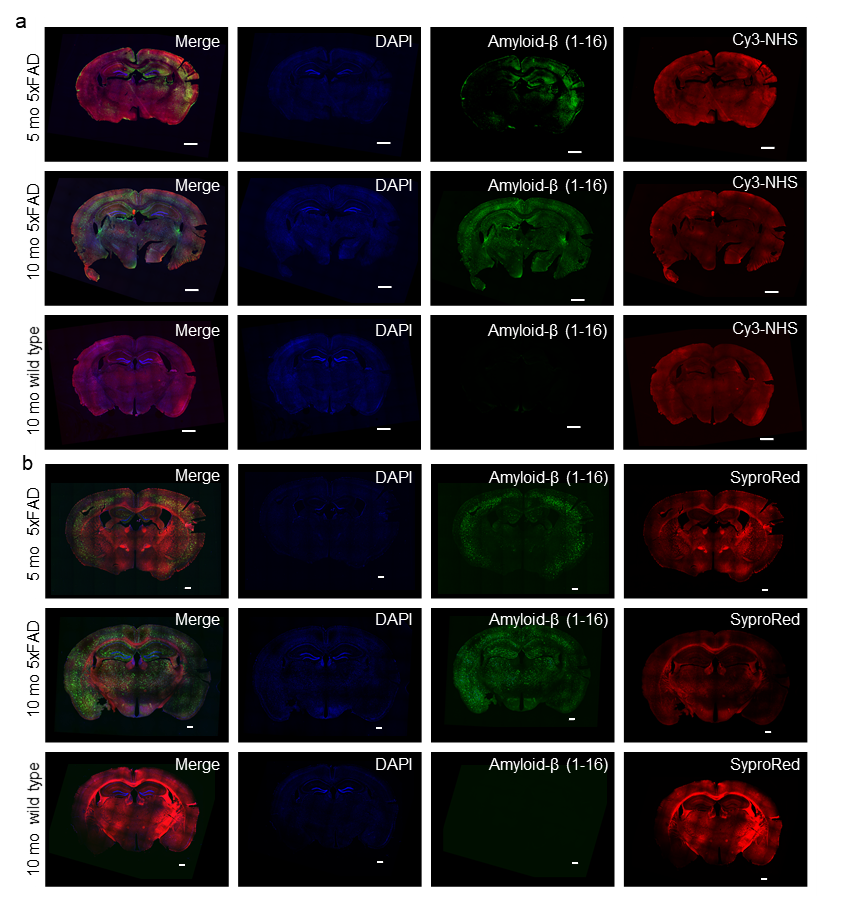

(a) Representative fluorescence images of brain tissue slices from 5-month-old female 5xFAD (5 mo 5xFAD), 10-month-old female 5xFAD (10 mo 5xFAD), and 10-month-old female C57BL/6J (10 mo wild type) mice immunofluorescently stained with anti-Amyloid-β (1-16) (green) and counterstained with DAPI (blue) and Cy3-NHS (red; n=3 slices from one mouse each). Scale bars, 1000 μm. (b) Representative fluorescence images of expanded brain tissue slices from 5-month-old female 5xFAD (5 mo 5xFAD), 10-month-old female 5xFAD (10 mo 5xFAD), and 10-month-old female C57BL/6J (10 mo wild type) mice immunofluorescently stained with anti-Amyloid-β (1-16) (green) and counterstained with DAPI (blue) and SyproRed (red) after processing with microProtemEx protocol (n=3 slices from one mouse each). Scale bar = 1000 μm (LEF=~2.5).

**Supplementary Figure 10.** SyproRed staining facilitates super-resolution imaging of the brain tissue samples processed with microProteomEx.

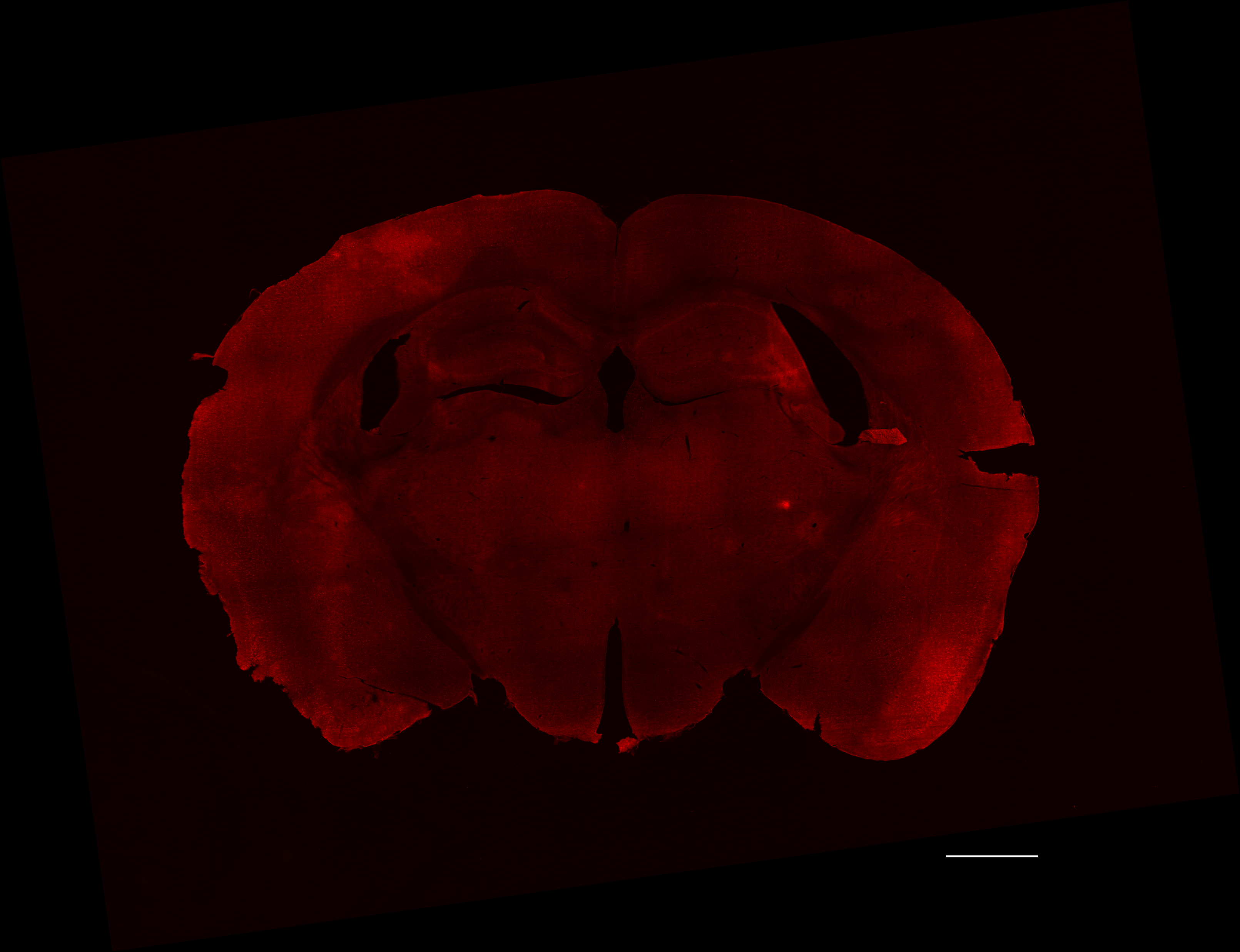

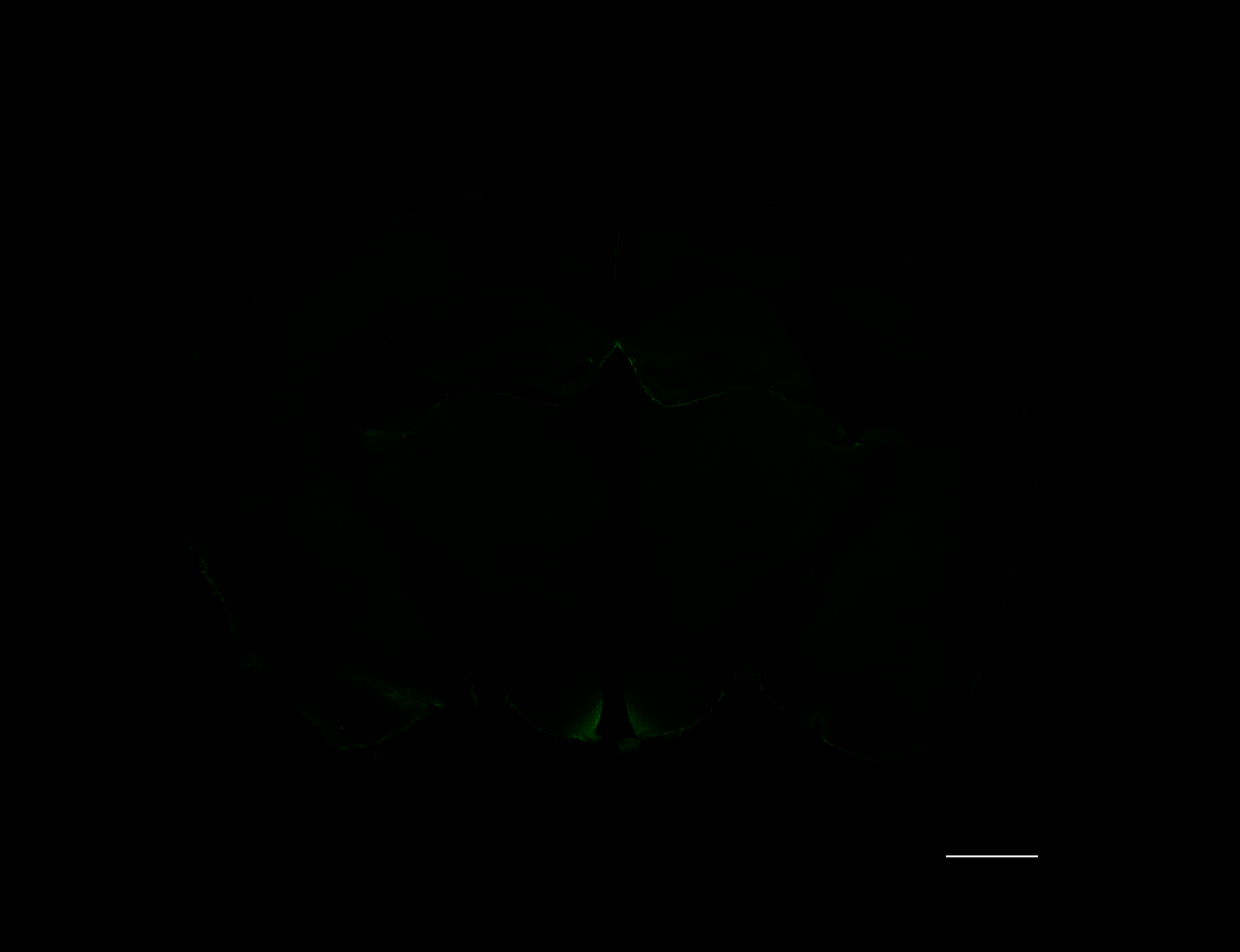

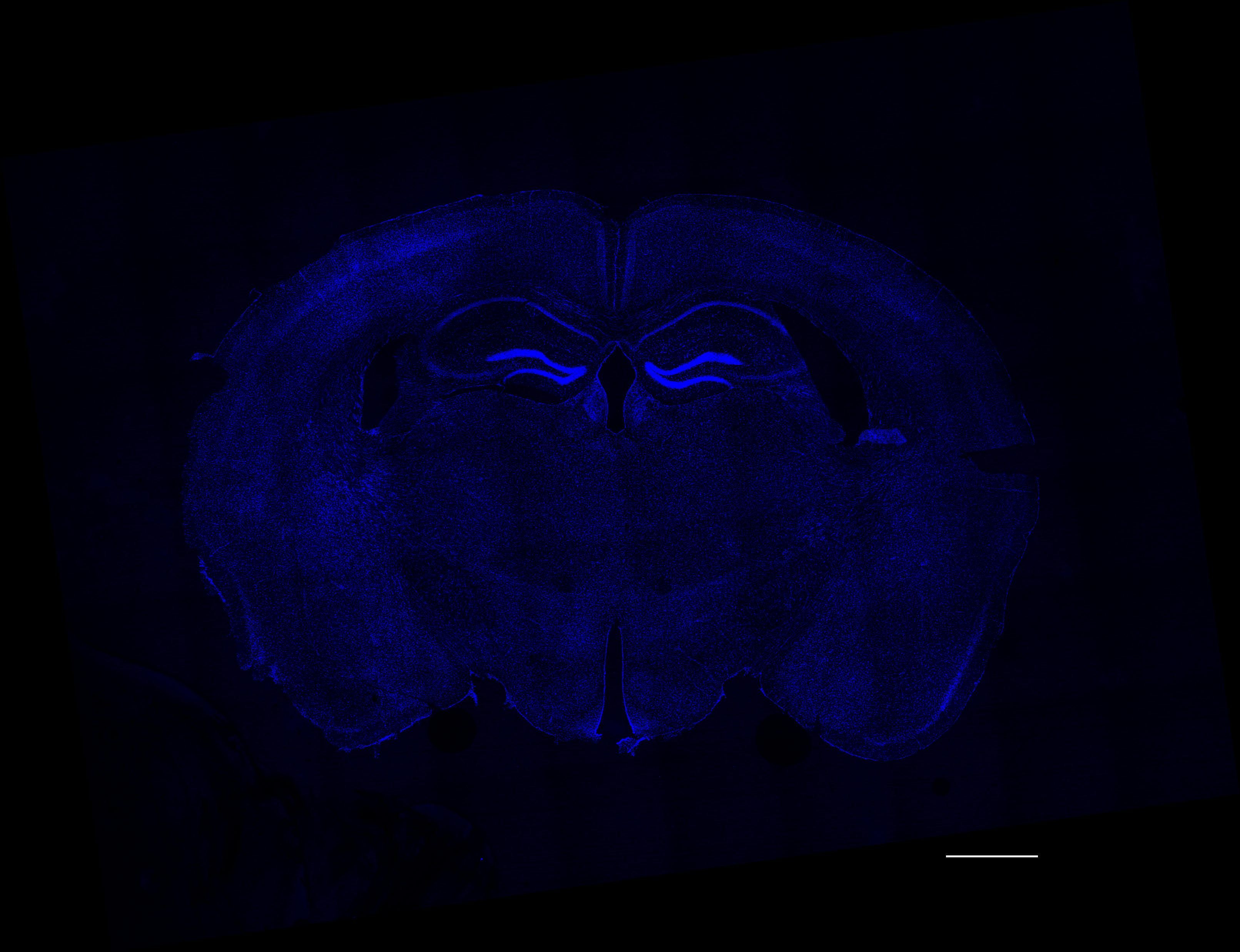

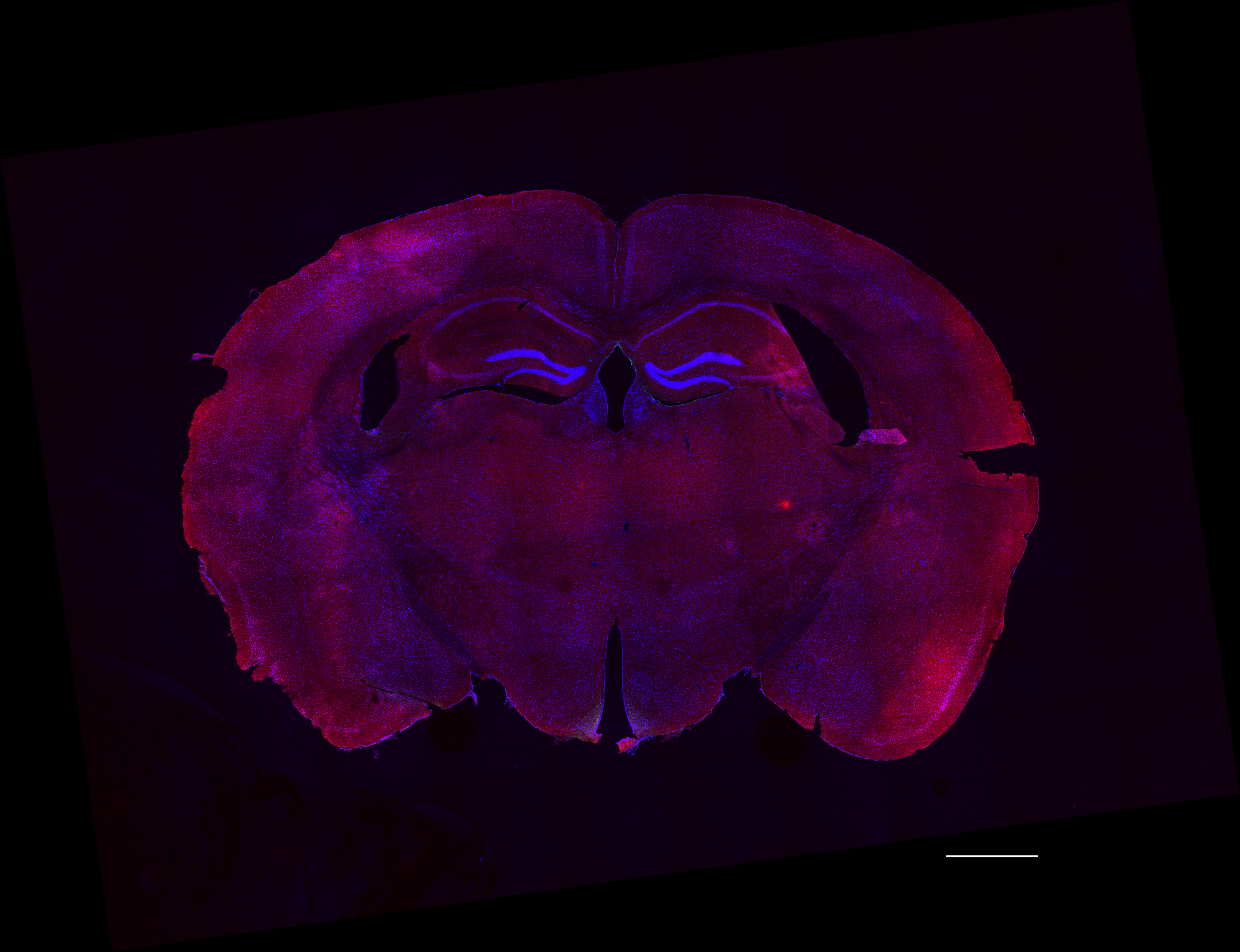

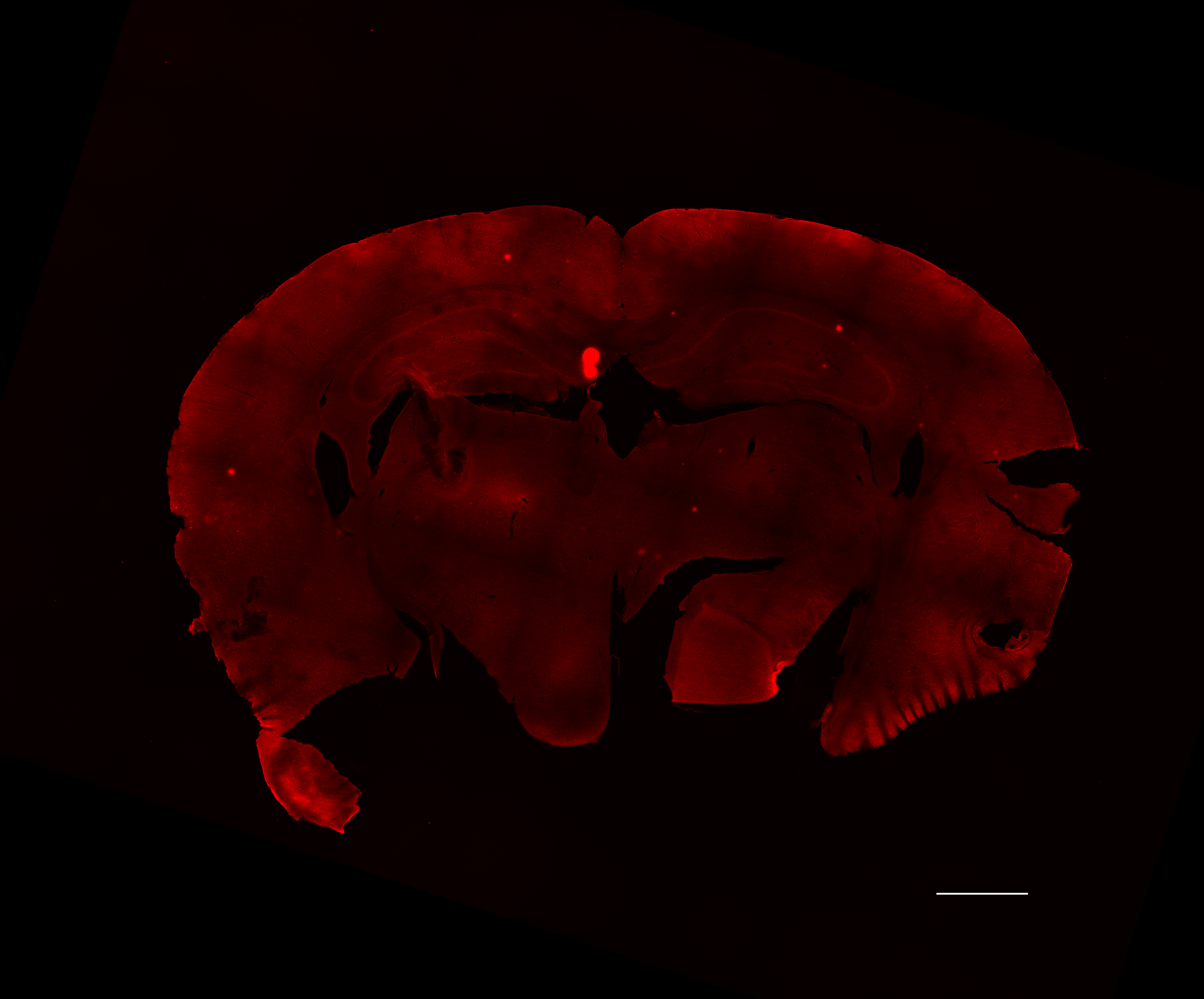

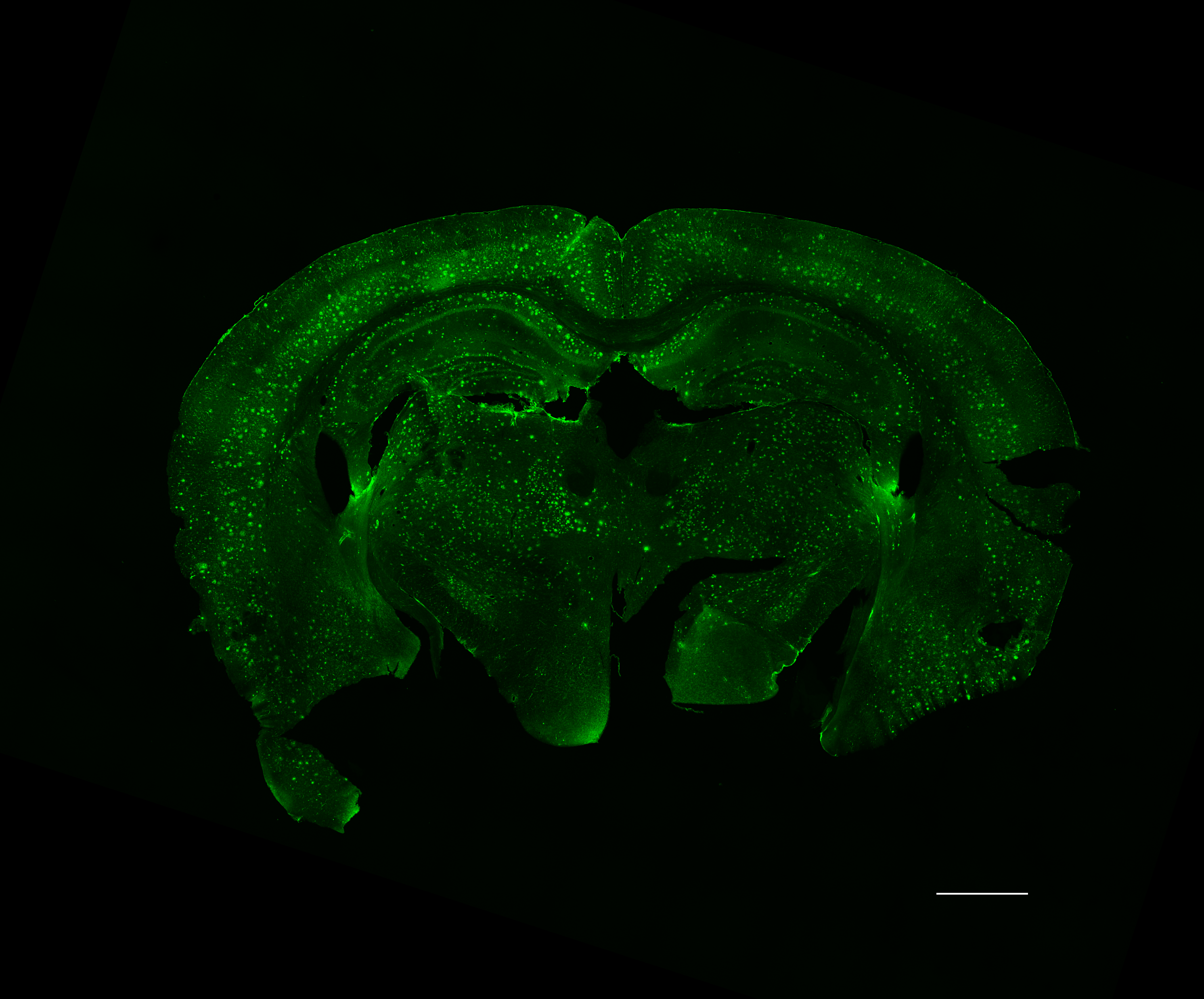

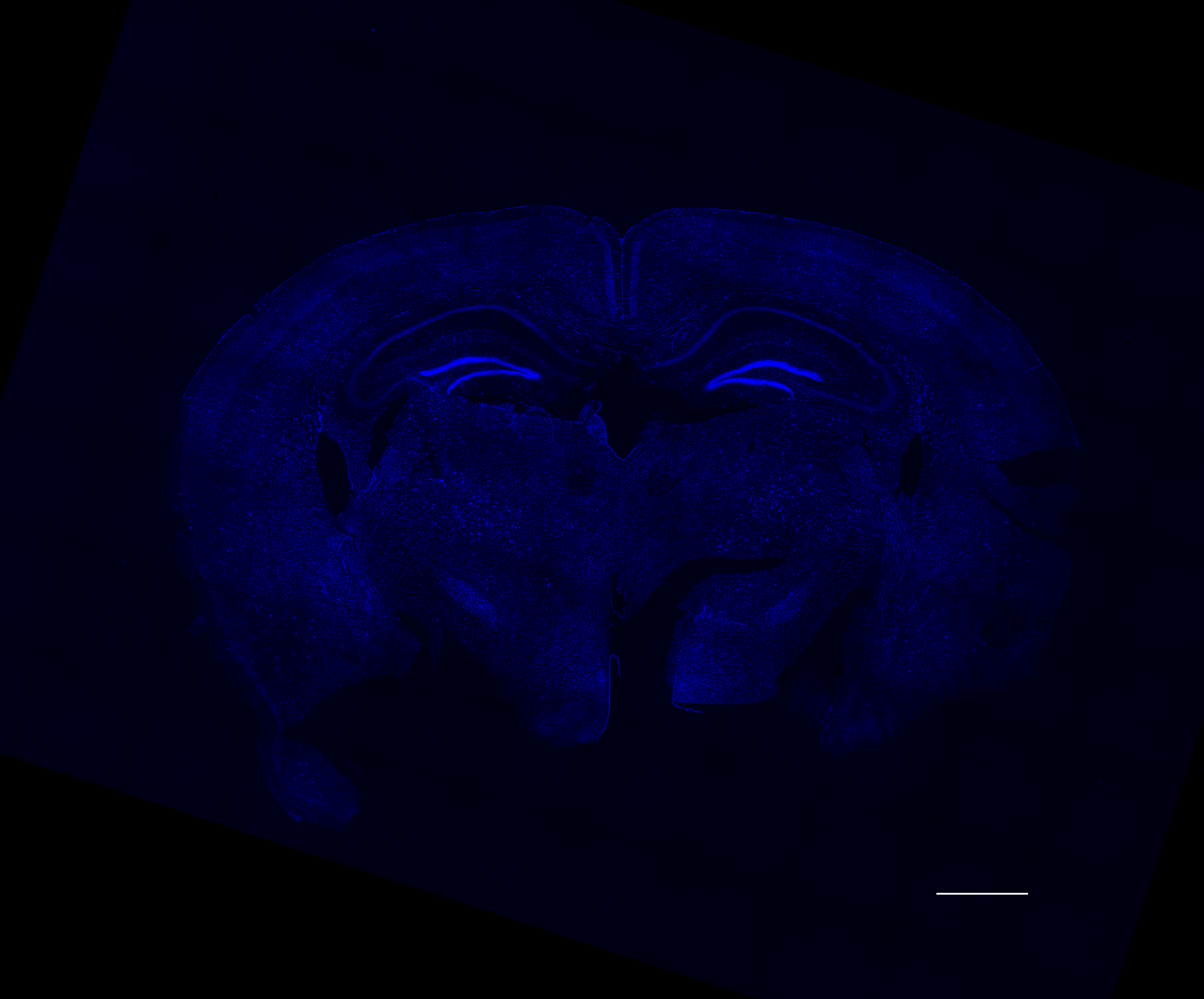

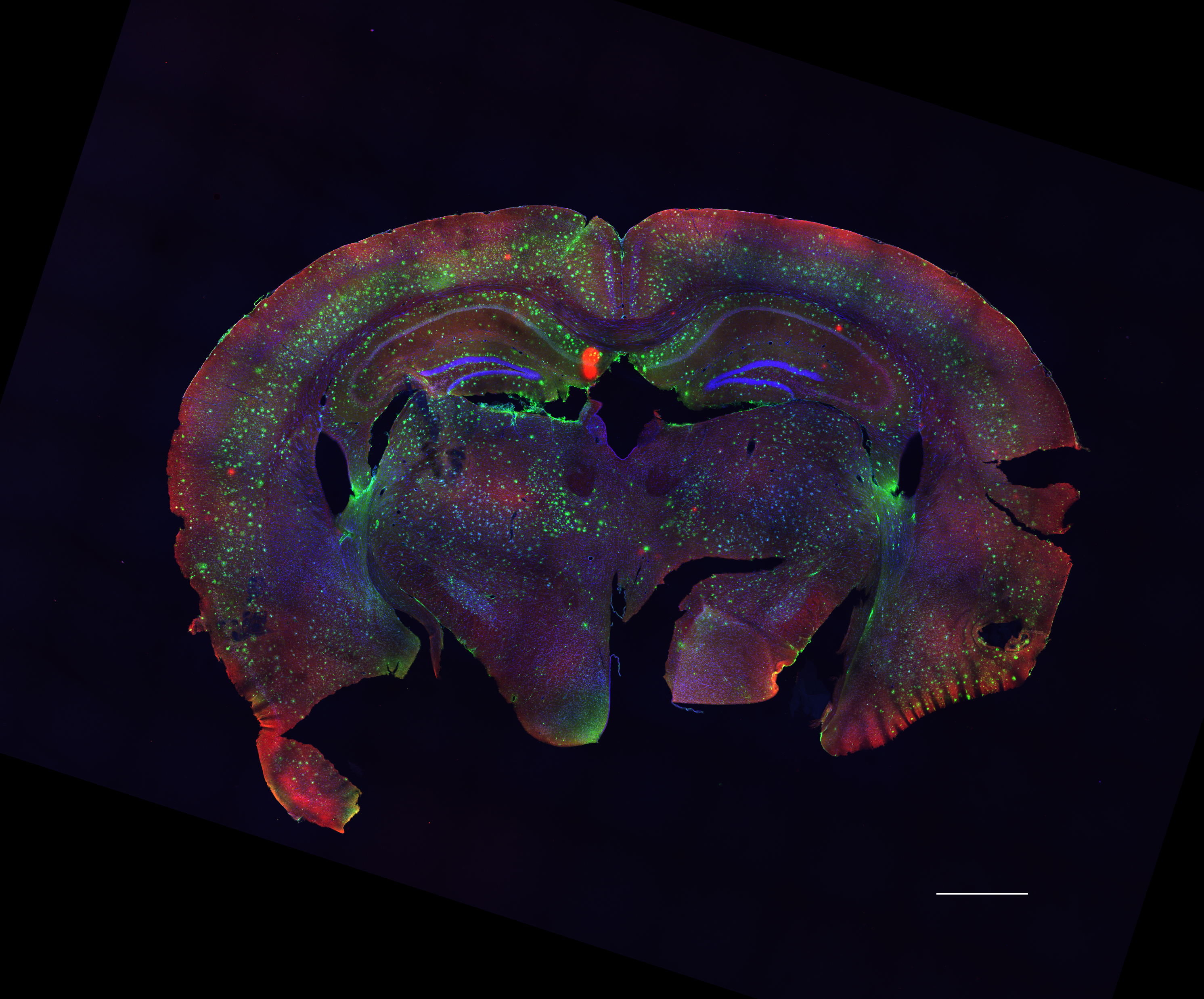

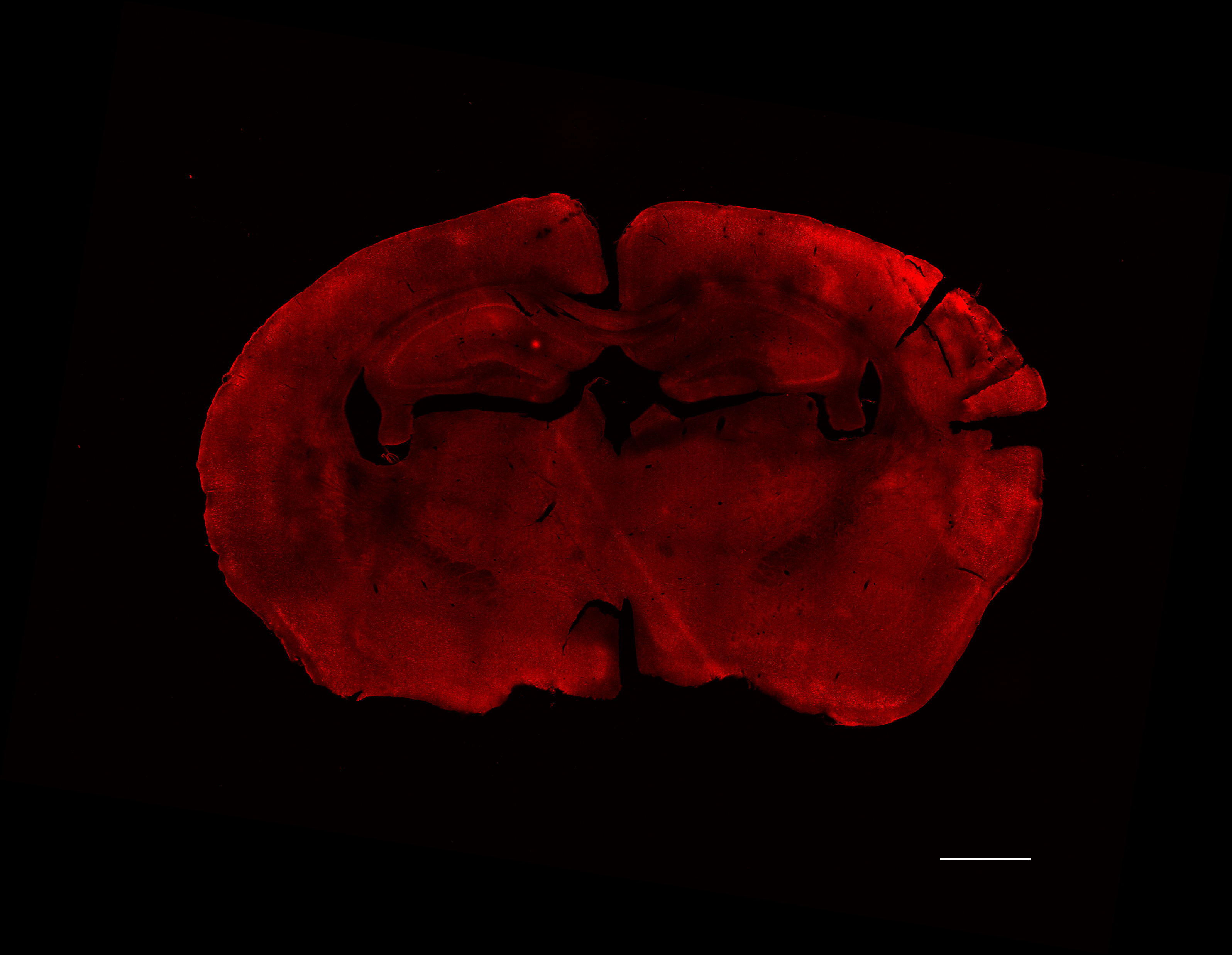

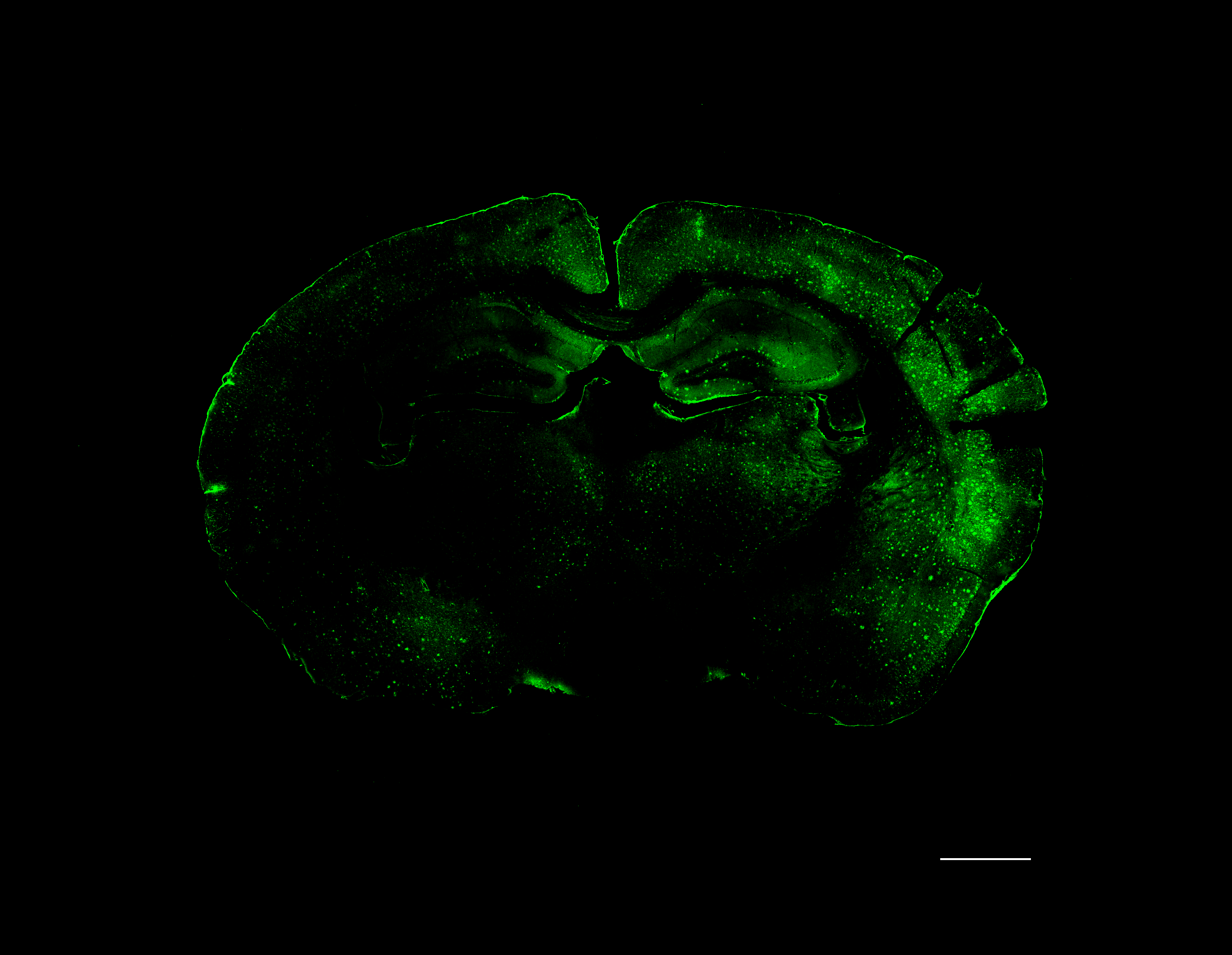

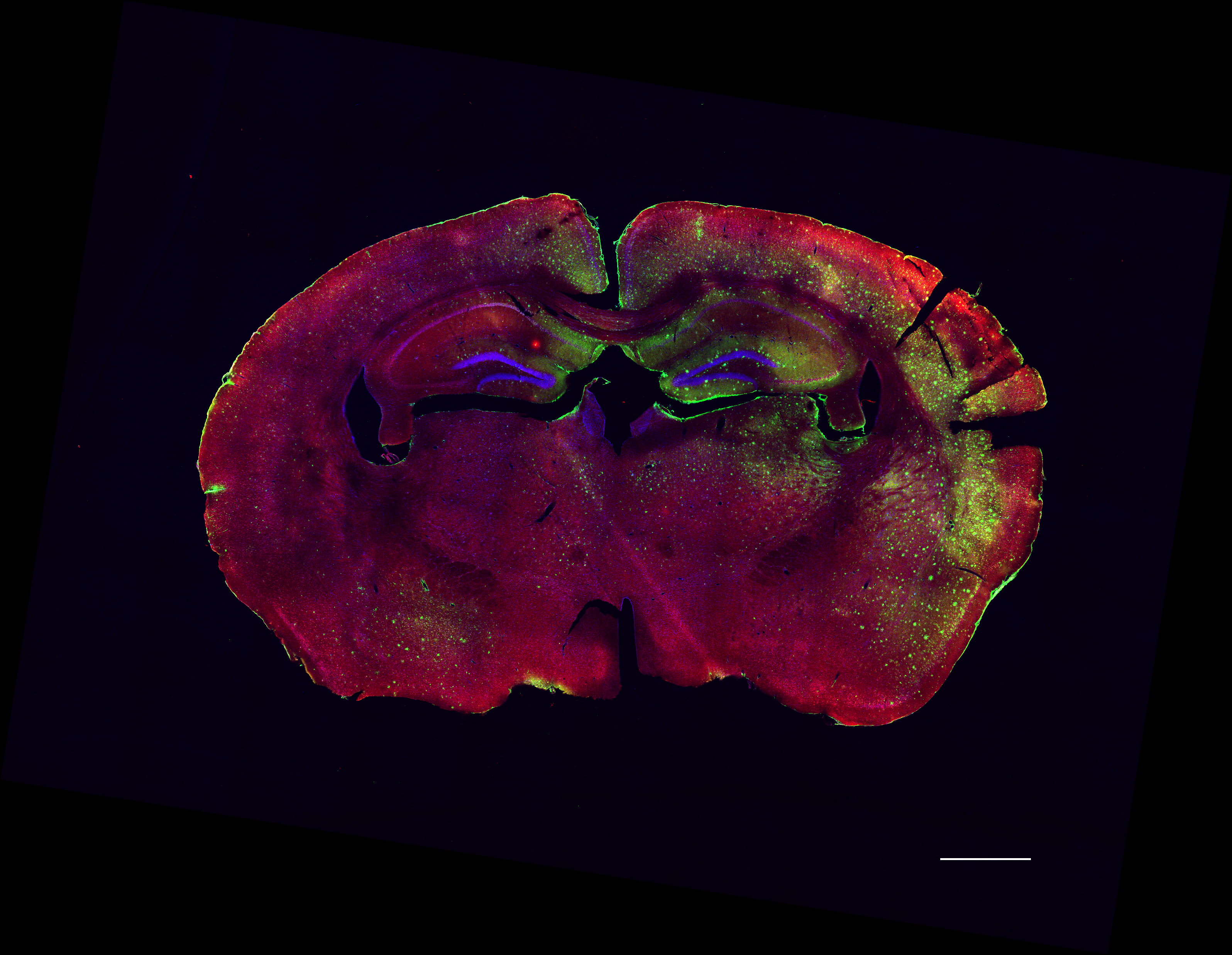

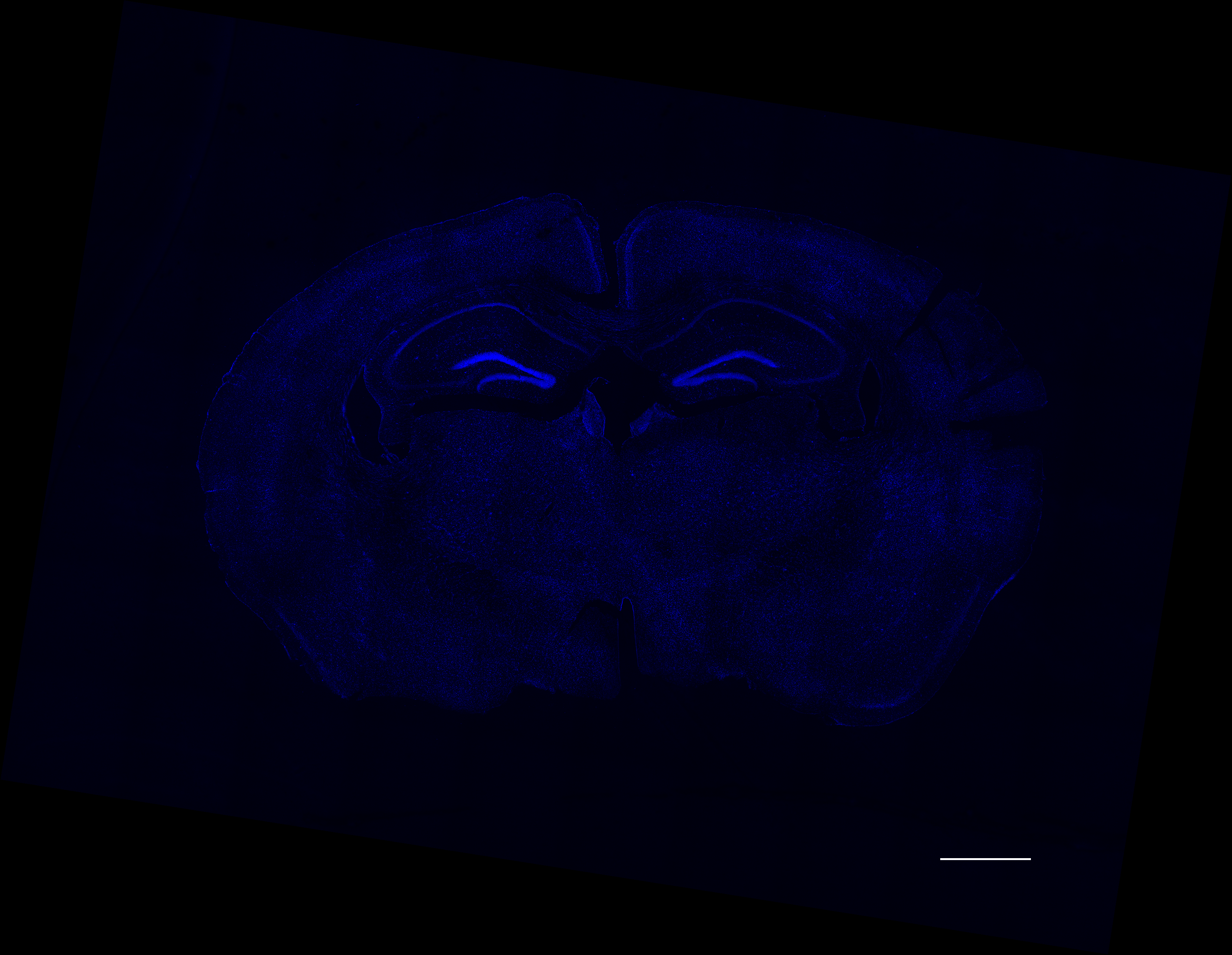

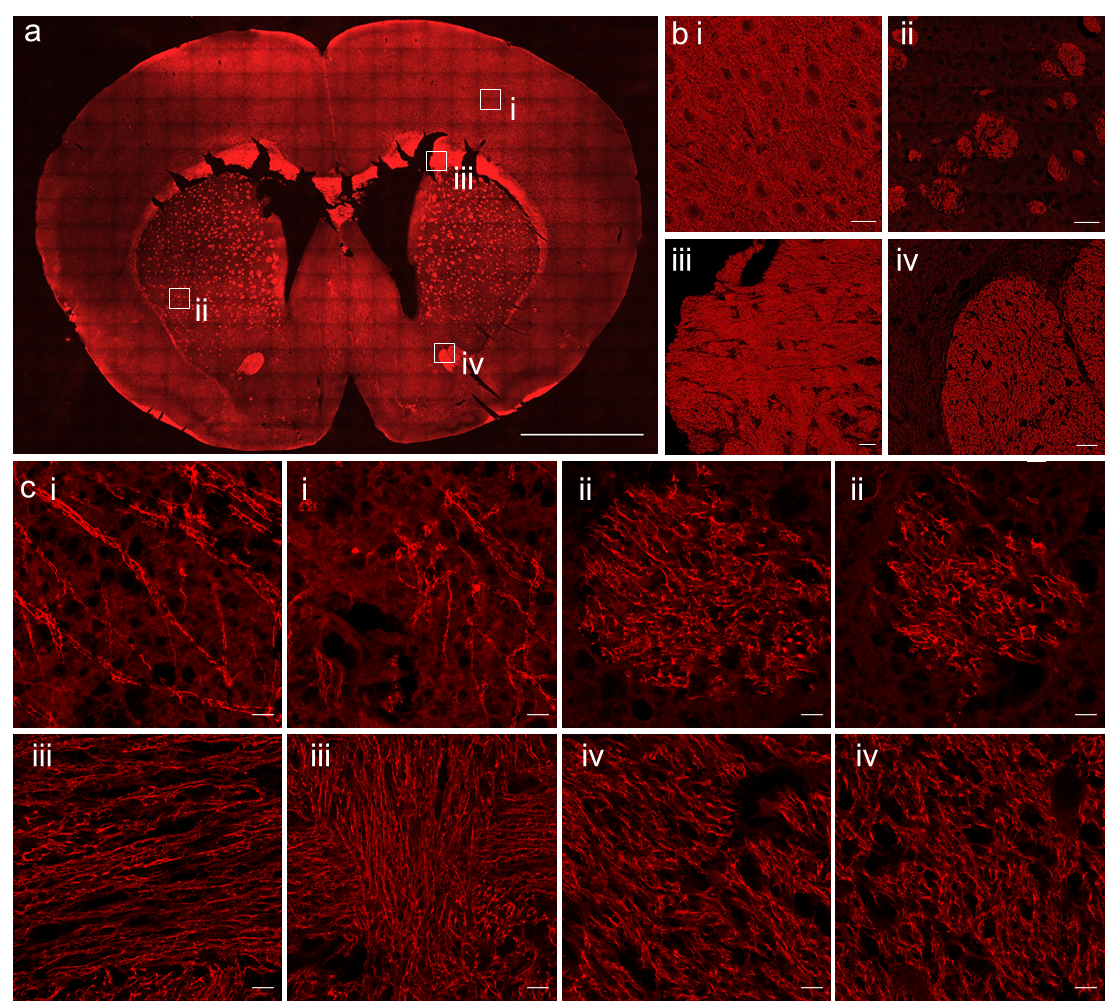

Merge

Merge

DAPI

DAPI

Old wild type

Old 5xFAD

Young 5xFAD

Cy3-NHS

Cy3-NHS

Amyloid-β (1-16)

Amyloid-β (1-16)

(a) Representative fluorescence image of post-expansion mouse brain processed with microProteomEx and stained with SyproRed (n = 3 slices from one mouse). Scale bar, 161.5 µm (LEF = 6.1 - 6.2 for all images throughout Supplementary Figure 9; physical size post-expansion, 1000 µm). (b) Single-plane confocal fluorescence images of different brain regions shown in boxes in panel a: i (cortex), ii (striatum), iii (corpus callosum), iv (anterior commissure). Scale bar, 16.2 µm (i, iii, iv (physical size post-expansion, 100 µm)), 32.3 µm (ii (physical size post-expansion, 200 µm)). Magnification, 40x (i, iii, iv (estimated resolution, 50 nm)), 20x (ii (estimated resolution, 76.2 nm)). (c) Representative single-plane super-resolution confocal fluorescence images of different brain regions as marked in panel b. Scale bar, 1.62 µm (i, ii, iii, iv (physical size post-expansion, 10 µm)), magnification, 40x (i, ii, iii, iv (estimated lateral resolution, 50 nm)).

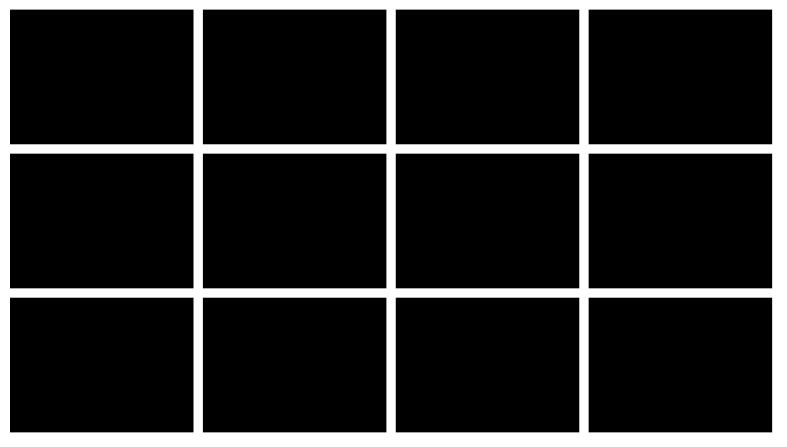

**Supplementary Figure 11.** Types of amyloid plaques identified in 5xFAD brain tissues.

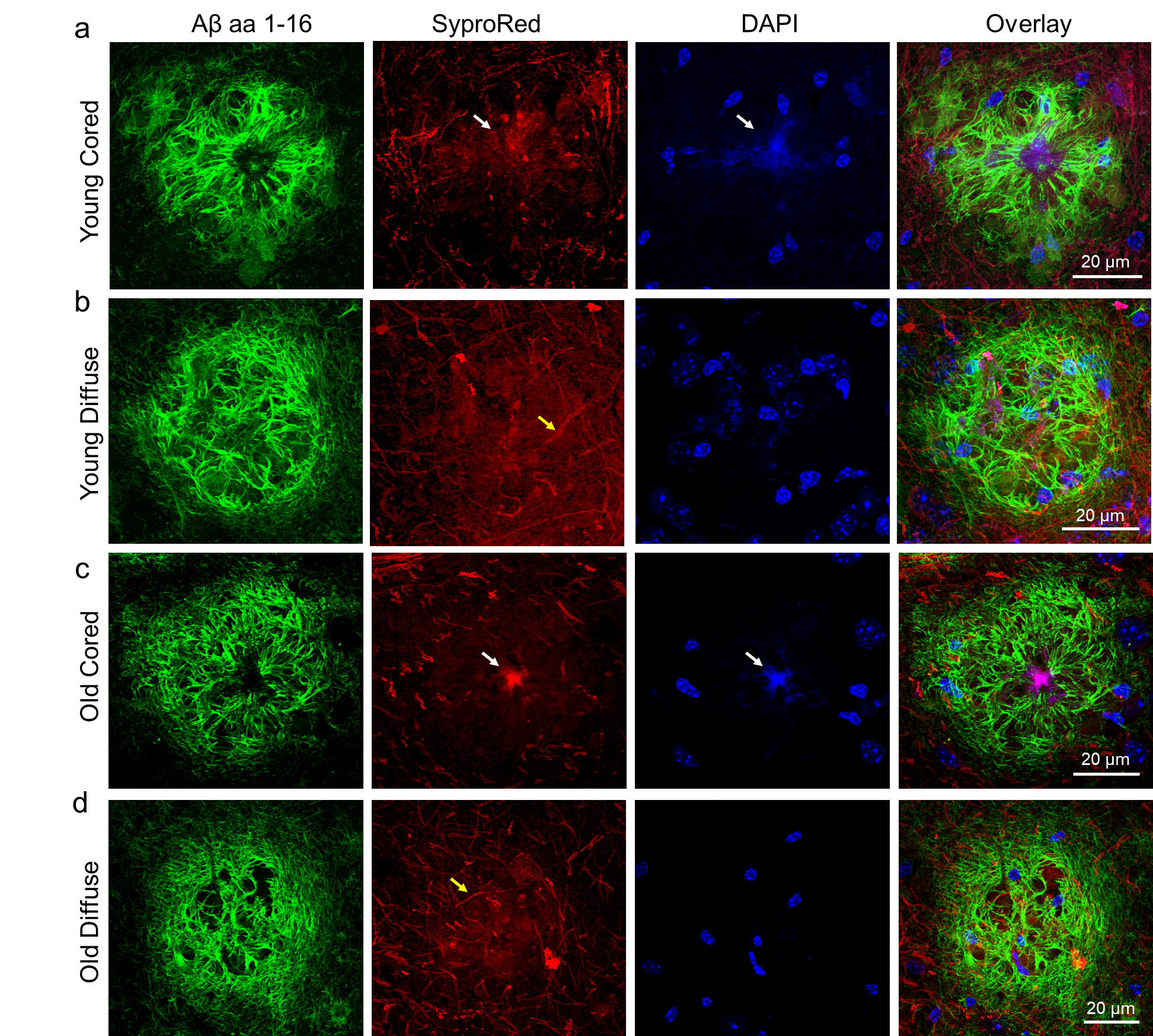

Representative maximum intensity projection confocal images of two types of amyloid plaques identified in 5xFAD brain tissue, immunostained with anti-Amyloid-β (1-16) (green) and counterstained with DAPI (blue) and SyproRed (red) post-expansion with mircoProteomEx protocol. (a) Representative images of cored plaque from a 5-month-old female 5xFAD mouse (n=25 plaques from 3 slices from one mouse). LEF=2.5. (b) Representative images of diffuse plaque from a 5-month-old female 5xFAD mouse (n=20 plaques from 3 slices from one mouse). LEF=2.5. (c) Representative images of cored plaque from 10-month-old female 5xFAD mouse (n=25 plaques from 3 slices from one mouse). LEF=2.5. (d) Representative images of diffuse plaque from 10-month-old female 5xFAD mouse (n=20 plaques from 3 slices from one mouse). LEF=2.5. White arrow in panels a and c indicates core of the plaque stained with SyproRed and DAPI. Yellow arrow in b and d indicates intact neurites going through the plaque.

**Supplementary Figure 12.** LCM isolation of selected plaques from brain tissue.

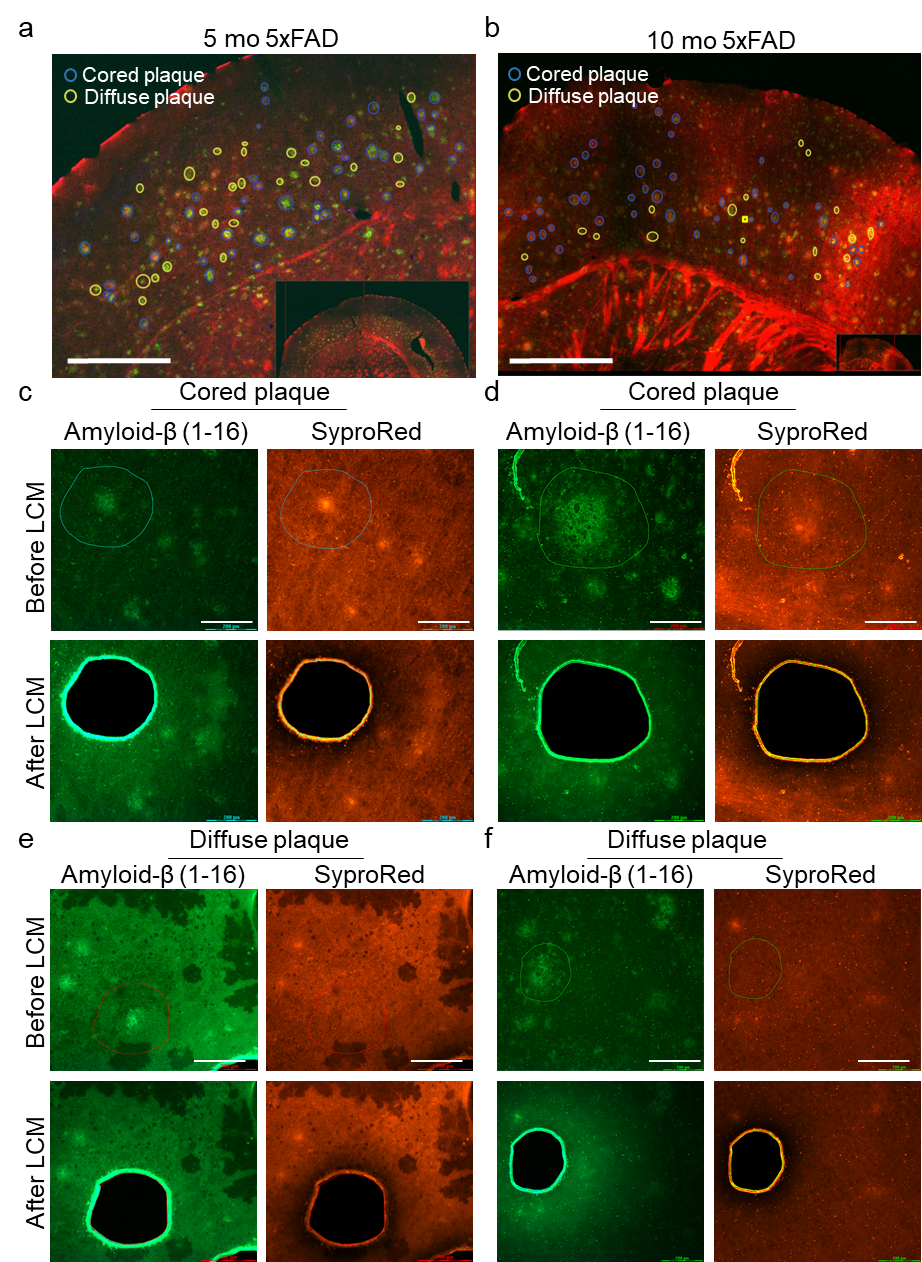

(**a**) Representative low magnification fluorescence image of cortex with marked cored and diffused plaques selected for LCM isolation (n=3 slices from one 5-month-old female 5xFAD mouse). Inset: low magnification imaging of cortex taken under LCM microscope in green and red channels. Scale bar, 500 µm; LEF=2.5. (**b**) Representative low magnification fluorescence image of cortex with marked cored and diffused plaques selected for LCM isolation (n=3 slices from one 10-month-old female 5xFAD mouse). Inset: low magnification imaging of cortex taken under LCM microscope in green and red channels. Scale bar, 200 µm; LEF=2.5. (**c-e**) Representative high magnification images of the individual (**c, d**) cored plaques and (**e, f**) diffuse plaques from (**c, e**) 5 mo and (**d, f**) 10 mo female 5xFAD mice before and after microdissection with LCM (n=~30 plaques from one mouse each). Scale bars, 200 µm; LEF=2.5.

**Supplementary Figure 13.** Reproducibility and variability assessment of protein identifications in plaque samples.
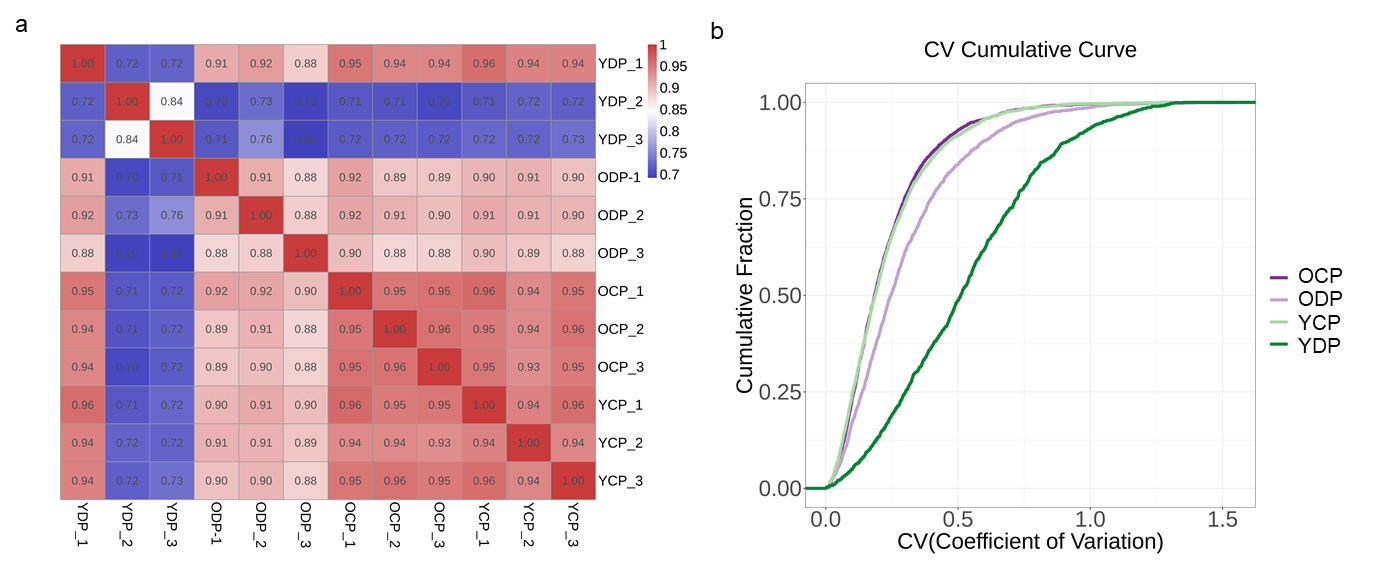

(a) Heatmap of Pearson correlation coefficients across two types of amyloid plaque samples, shown in Fig. 6. (b) The coefficient of variation cumulative curve for samples shown in a.

**Supplementary Figure 14.** Proteomic comparison across cored and diffuse plaque types in old and young AD mice.

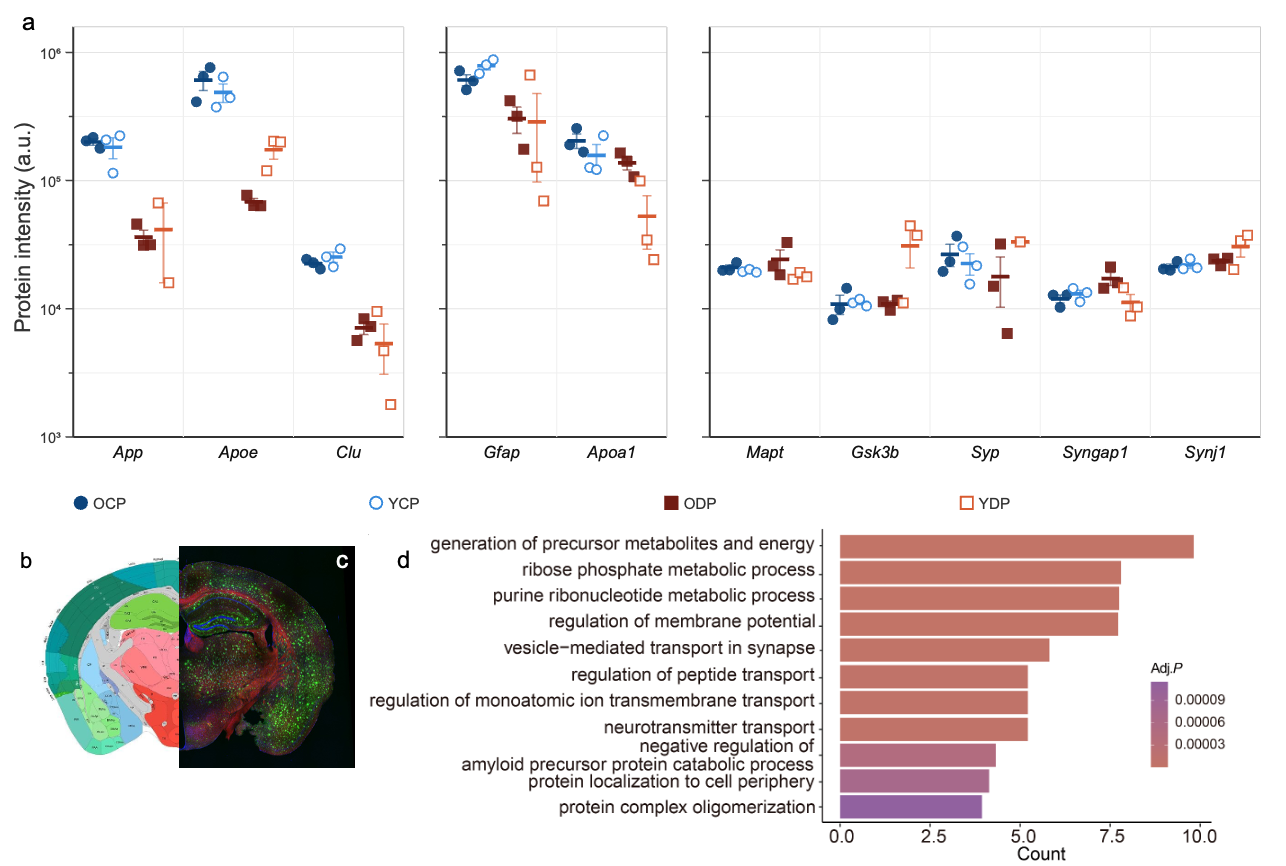

(a) Dot plots showing the intensity of individual proteins quantified by proteomics across four plaque-associated conditions: old cored plaque (OCP, filled blue circle), young cored plaque (YCP, open blue circle), old diffuse plaque (ODP, filled dark red square), and young diffuse plaque (YDP, open dark red square). Proteins are grouped by biological category: Aβ metabolism and clearance (App, Apoe, Clu; left), reactive astrogliosis and lipoprotein transport (Gfap, Apoa1; middle), and tau-related and synaptic proteins (Mapt, Gsk3b, Syp, Syngap1, Synj1; right). The Y-axis represents protein intensity in arbitrary units (a.u.) on a log₁₀ scale. Horizontal bars indicate the mean; error bars represent SEM (n = 2–3 plaques per condition). (b) Representative fluorescence image of a mouse brain section showing plaque distribution across anatomical regions. (c) Allen Brain Atlas reference overlay corresponding to the section shown in (b), with color-coded anatomical regions. (d) Gene ontology (GO) biological process enrichment analysis of proteins differentially enriched in YP versus YP. Bars represent the number of proteins annotated to each GO term; bar color indicates the adjusted P-value (Adj. P).

**Supplementary Movie 1.** Three-dimensional confocal rendering of expanded mouse liver tissue stained with SyproRed.

**Supplementary Movie 2.** Volumetric confocal rendering of an individual glomerulus in expanded mouse kidney tissue, visualized with SyproRed staining.

**Supplementary Movie 3.** Volumetric confocal rendering of intact cardiomyocytes in expanded mouse heart tissue, with clearly delineated sarcomere structures, visualized with SyproRed staining.

**Supplementary Movie 4.** Volumetric confocal rendering of the cytoarchitecture within the GCMN region of expanded tissue, visualized with SyproRed staining.

**Supplementary Movie 5.** Volumetric confocal rendering of the cytoarchitecture within the MM region of expanded tissue, visualized with SyproRed staining.

**Supplementary Movie 6.** Volumetric confocal rendering of single Aβ plaque within intact cortical tissue from a 5xFAD mouse model, following expansion, immunostaining with Aβ 1-16 antibody (green) and staining with SyproRed (red) and DAPI (blue).

**Supplementary Movie 7.** Volumetric confocal rendering of single Aβ plaque within intact cortical tissue from a 5xFAD mouse model, following expansion, immunostaining with Aβ 1-16 antibody (green) and staining with SyproRed (red) and DAPI (blue).

**Supplementary Movie 8.** Volumetric confocal rendering of immunostained Aβ plaques in expanded brain slices from 10-month-old (late-stage disease) female 5xFAD mice, processed using the microProteomEx workflow. The movie illustrates representative cored plaque morphologies targeted for laser capture microdissection (LCM), visualized with immunostaining with Aβ 1-16 antibody (green) and staining with SyproRed (red) and DAPI (blue).

**Supplementary Movie 9.** Volumetric confocal rendering of immunostained Aβ plaques in expanded brain slices from 10-month-old (late-stage disease) female 5xFAD mice, processed using the microProteomEx workflow. The movie illustrates representative coarse plaque morphologies targeted for laser capture microdissection (LCM), visualized with immunostaining with Aβ 1-16 antibody (green) and staining with SyproRed (red) and DAPI (blue).
